# Decoding epitope immunodominance in HIV Env using cryoEM and machine learning

**DOI:** 10.64898/2026.03.09.710672

**Authors:** Jan S Schuhmacher, Shuhao Xiao, Elise R Eray, Sharidan Brown, Alexander Zambrowski, Aryan Jain, Daniel Montiel Garcia, Gabriel Ozorowski, Wenwen Zhu, Katrina Saam, Tom G Caniels, John P Moore, Max Crispin, Rogier W Sanders, Srirupa Chakraborty, Bruno E Correia, Andrew B Ward, Aleksandar Antanasijevic

## Abstract

Viral surface glycoproteins, such as the HIV envelope protein (Env), present numerous antibody (Ab) epitopes, yet immune responses consistently focus on only a subset, a phenomenon known as immunodominance. Although structural studies have provided insights into Env antigenicity, our understanding of the molecular features that govern efficient Ab engagement remains incomplete, thereby limiting the predictive and rational design of vaccines. Here, we characterized the structural determinants of epitope immunodominance in HIV Env by integrating high-resolution cryoEM-based polyclonal epitope mapping (cryoEMPEM) across different clades with quantitative analyses of epitope topology, accessibility, and physicochemical properties. More than 70 new structures were resolved to assemble a library of >100 Env-antibody complexes. These data informed the development of a surface-centric, machine-learning model to predict relative **A**ntigen **S**urface **I**mmunodominance (ASI model). Comparison of ASI-predicted epitope sites with the specificities of Env-induced antibodies showed that the model accurately identifies immunodominant regions and highlights the structural features driving immune bias. Notably, immunogens redesigned based on model predictions successfully redirected Ab responses toward a normally subdominant epitope, demonstrating the potential of strategies coupling targeted assembly of focused structural libraries with machine learning to uncover complex molecular patterns and enable design of more effective vaccine antigens.

## INTRODUCTION

Recognition of foreignness is a central function of the immune system, mediated by evolved interactions between immune receptors and invading pathogens^1^. Humans harbor ∼10^9^ B cells, which further diversify in secondary lymphoid organs to produce antibodies (Ab) with high affinities (micromolar to picomolar in most cases) for discrete molecular elements on the antigen surface called epitopes^2,3^. Affinity maturation occurs through somatic hypermutation in the complementarity-determining regions (CDRs) of antibodies, producing paratopes with optimized geometry and physicochemical complementarity to these sites. This process is probabilistic, depending on both the diversity and competence of the host B-cell repertoire, as well as on the structural and chemical properties of individual epitopes^4^. As a result, different antigens and distinct regions within the same antigen vary in their likelihood of being targeted, leading to a hierarchical distribution of responses, a phenomenon known as immunodominance.

This is perhaps best exemplified in viral glycoproteins, such as the human immunodeficiency virus envelope (HIV Env). These antigens present a broad array of conformational epitopes, yet antibody responses are preferentially directed to variable (V) regions that continue to diversify under immune pressure^5^. In contrast, conserved (C) regions are targeted less frequently despite surface accessibility, indicating that epitope exposure alone does not guarantee effective affinity maturation^6,7^. As a result, immunodominance shapes viral evolution and challenges the design of broadly-protective vaccines. Prior attempts to link immunodominance in HIV Env to structural and chemical features of antigenic sites, stemmed from the established mechanisms of immune evasion^5,8,9^. Glycan-mediated molecular mimicry is the best-characterized example, in which N-linked glycans reduce the immune targeting of proximal peptidic elements via steric hindrance and central tolerance^10–14^. Multivalent Env presentation was also found to modulate epitope accessibility, suppressing responses to basal sites while enhancing responses to apex-proximal epitopes^15–17^. Beyond these insights, the molecular principles governing immunodominance remain incompletely defined or anecdotal.

This limited insight has hindered the development of computational methods to predict or rationally engineer immunodominance of conformational epitopes in HIV and other viral antigens. Structural data are essential for understanding epitope targeting, but the scale and diversity of B-cell responses present a major challenge. In HIV, a single immunization or infection can generate thousands of Env-specific Abs with unique sequences^18,19^, far exceeding the throughput of structural biology approaches. To date, there are ∼200 annotated structures of Ab-Env complexes in the SAbDab, most of which comprise broadly-neutralizing antibodies (bnAbs) or their precursors targeting conserved subdominant epitopes^20,21^. As a result, existing monoclonal Ab structures are not optimal inputs to achieve systems-level understanding of immune response. High-throughput methods such as peptide microarrays^22^ and phage display^23^ provide alternatives for epitope mapping. Still, they are largely limited to linear epitopes, which represent only a minor fraction of antigenic landscape in most antigens^24^.

In this study, we addressed the limitations of available structural data by applying cryoEM-based polyclonal epitope mapping (cryoEMPEM) to characterize polyclonal responses and map immunodominant antigenic space across six HIV Env antigens from different clades and geographic origins. By analyzing polyclonal antibodies from prior rabbit and macaque immunization studies, we assembled a library of over 100 high-resolution maps with antibodies bound to unique epitopes. These data enabled a quantitative analysis of the structural and physicochemical determinants of immunodominance and served as the foundation for a machine learning (ML) model to predict epitope targeting in HIV Env. We validated the model on an existing HIV vaccine candidate and demonstrated its utility in engineering immunodominance in BG505 SOSIP trimers through targeted amino-acid modifications. Overall, our study integrates focused structural characterization of immunodominant antibody responses with predictive modeling, providing new tools and conceptual frameworks for rational vaccine design.

## RESULTS

### Diverse HIV Env immunogens present distinct antigenic profiles

To map immunodominant epitopes across different Env antigens, we employed cryoEMPEM^25^. This approach enables the structural characterization of Ab-antigen complexes using polyclonal antibodies derived from immune sera, thereby allowing the simultaneous reconstruction of multiple electron microscopy maps that capture the most prevalent classes of antigen-specific antibodies. Serum samples from prior rabbit and macaque immunization studies were processed as described in the **Methods** and illustrated in **Fig. 1A**. Owing to the size and complexity of their B-cell repertoires and their ability to generate Abs with long complementarity-determining region 3 (CDR3) loops, these species serve as optimal models for human-like epitope targeting, even in cases of heavily glycosylated antigens such as Env^27–29^.

**Figure 1.**
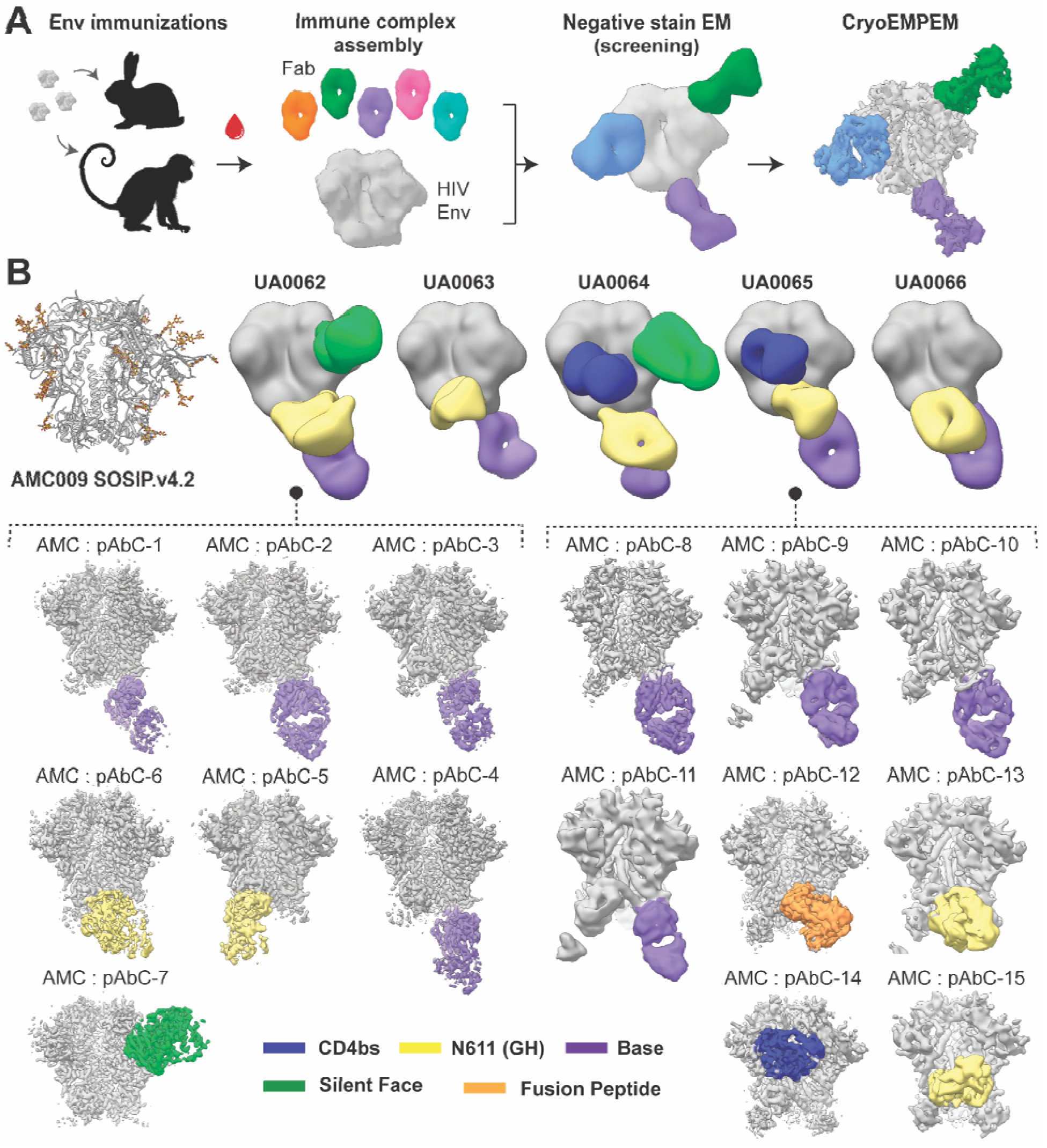
Recovering immunodominant antibodies to HIV Env using EMPEM. (**A**) Workflow used to identify dominant classes of Env-specific antibodies from immunized animals. (**B**) Structure of AMC009 SOSIP.v4.2 (PDB: 6VO3, top left) and composite negative-stain EMPEM reconstructions from rabbit polyclonal sera (top row; animal IDs shown above). CryoEMPEM maps of AMC009 SOSIP.v4.2 immune complexes with antibodies from rabbits UA0062 and UA0065 are shown below. The antigen is colored gray, and Abs are colored by epitope specificity.

We characterized immunodominant epitopes of representative Env antigens from HIV-1 clades A, B, and C (**Table 1**), as well as a consensus sequence of the M subtype (ConM). CH505(B) denotes a chimeric construct in which variable loops from CH505 Env were grafted onto the BG505 backbone^30^. In the context of stabilized native-like ectodomain constructs (SOSIP), these antigens have been extensively evaluated as vaccine candidates^31–36^. Pairwise sequence identities range from 74-85%, with the greatest divergence localized to the surface-exposed variable loops (V1-V5; **fig. S1**). Serum samples were obtained following three immunizations with the corresponding Env antigen, yielding detectable antigen-specific Ab titers in all animals analyzed^16,17,37,38^. Polyclonal Abs were prepared as fragment antigen binding (Fab) using a previously established protocol^39^, complexed with the immunizing Env antigen, and subjected to electron microscopy (**Fig. 1A**).

**Table 1:** Information on the Env antigens and immune samples.

| HIV Strain | Clade | Construct | Immunization | Animal model |
| --- | --- | --- | --- | --- |
| BG505 | A | SOSIP, MD39 | Various<sup>26,30,40</sup> | Rabbit, Macaque |
| B41 | B | SOSIP.v4.1 | C0045-15<sup>41</sup> | Rabbit |
| AMC009 | B | SOSIP.v4.2 | C0048-15<sup>37</sup> | Rabbit |
| 16055 | C | SOSIP.v8.3 | C0038-19<sup>42</sup> | Rabbit |
| CH505(B) | A/C | SOSIP.v8.1 | C0038-19<sup>30</sup> | Rabbit |
| ConM | Consensus | SOSIP.v7 | Various<sup>16,43</sup> | Rabbit, Macaque |

The immune complexes were initially screened by negative-stain electron microscopy (nsEMPEM), and low-resolution reconstructions capturing distinct Ab specificities were generated. The resulting composite figures of the AMC009 SOSIP.v4.2 antigen in complex with structurally distinct antibodies, detected in each serum sample (animal), are shown in **Fig. 1B** (top), with corresponding data for B41 SOSIP.v4.1, and the CH505(B) SOSIP.v8.1 chimera in **fig. S2A and S2B**, respectively. Equivalent datasets for ConM SOSIP.v9, BG505 SOSIP (multiple versions), and 16055 SOSIP.v.8.3 were obtained from previous studies^16,17,26,30,40,43–46^, and the summary of epitope targeting data is presented in **fig. S2C**.

Analysis of per-animal datasets enabled mapping of distinct epitope specificities while accounting for inter-individual variability in antibody responses. Epitope definitions follow established conventions in the field^21,26^ and are detailed in **Table S1**. Certain epitopes, including the Base and N611 (GH), are frequently targeted across animals immunized with the same antigen and, notably, across different HIV Env antigens (**fig. S2**). In contrast, other sites, such as the Silent Face (SF) or CD4 binding site (CD4bs), are targeted less consistently and are only observed in 40% of animals immunized with AMC009 SOSIP.v4.2 (**Fig. 1B**). Although intra-group differences reflect the influence of B-cell repertoire and stochasticity in affinity maturation, the repeated targeting of immunodominant epitopes across animals and Env antigens points to epitope-intrinsic features as the primary determinant of antibody engagement.

This observation is supported by cross-species comparisons of epitope targeting (**fig. S2C**). For both ConM and BG505 SOSIP antigens, the main immunogenic epitopes detected in humans^44,45^ were also observed in nsEMPEM data from rabbit^16,46^ and macaque^26,43^ immunizations. Although some specificities and the distribution of antibody responses differed between species, the repeated detection of the same dominant epitopes across human and other animal models further emphasizes the contribution of epitope features to immunodominance. This pattern is not unique to HIV Env and similar findings have been reported for other viral glycoproteins, including influenza hemagglutinin^47–49^.

### High-resolution epitope mapping by cryoEMPEM

To enable precise epitope assignment and comparison between antigens, a subset of immune complexes was subjected to high-resolution cryoEMPEM analysis. Based on the nsEMPEM screen, 1-2 polyclonal samples per antigen encompassing all detected Ab specificities were selected (**Table S2**), amounting to nine immune complex samples imaged. Microscope and imaging details are provided in **Extended Data - Table 1**, while representative raw micrographs and 2D class averages are shown in **fig. S3.** The resulting data was processed using a focused classification workflow previously described^26^ and illustrated in **fig. S4**. From these datasets, 72 maps were generated, each comprising a single Env antigen in complex with distinct polyclonal antibodies as Fab fragments (**fig. S5-S13**), referred to as structurally unique polyclonal antibody classes (pAbC). Relevant map statistics, including particle number and resolution, are presented in **Extended Data - Table 2**, with the corresponding Fourier Shell Correlation (FSC) plots in **fig. S14-S18**. Examples of maps reconstructed for AMC009 SOSIP.v4.2 in complex with polyclonal antibodies from rabbits AU0062 and AU0065 are shown in **Fig. 1B** (bottom).

For analyses of epitope targeting, we incorporated previously published cryoEMPEM data of BG505 SOSIP MD39, BG505 SOSIP.v5.2 N241/N289, and BG505 SOSIP.v5.2 (7S) in complex with polyclonal Abs from rhesus macaques, comprising 35 maps and models of structurally distinct BG505 Abs^26,30,40^. Combined with the newly generated data, this yielded a total of 107 pAbC maps, ranging from 7 to 35 depending on the antigen (**Fig. 2A**). These maps unveiled the immunodominant antigenic surfaces for each antigen and, in most cases, provided overlapping coverage of targeted epitopes. Exceptions were the CD4bs and the SF, for which only a single map was recovered (**Fig. 2B**). Global map resolutions ranged from 2.9 Å to 9.9 Å, with an average of ∼4.5 Å, enabling interpretation at the level of individual amino acids necessary for accurate epitope definition.

**Figure 2.**
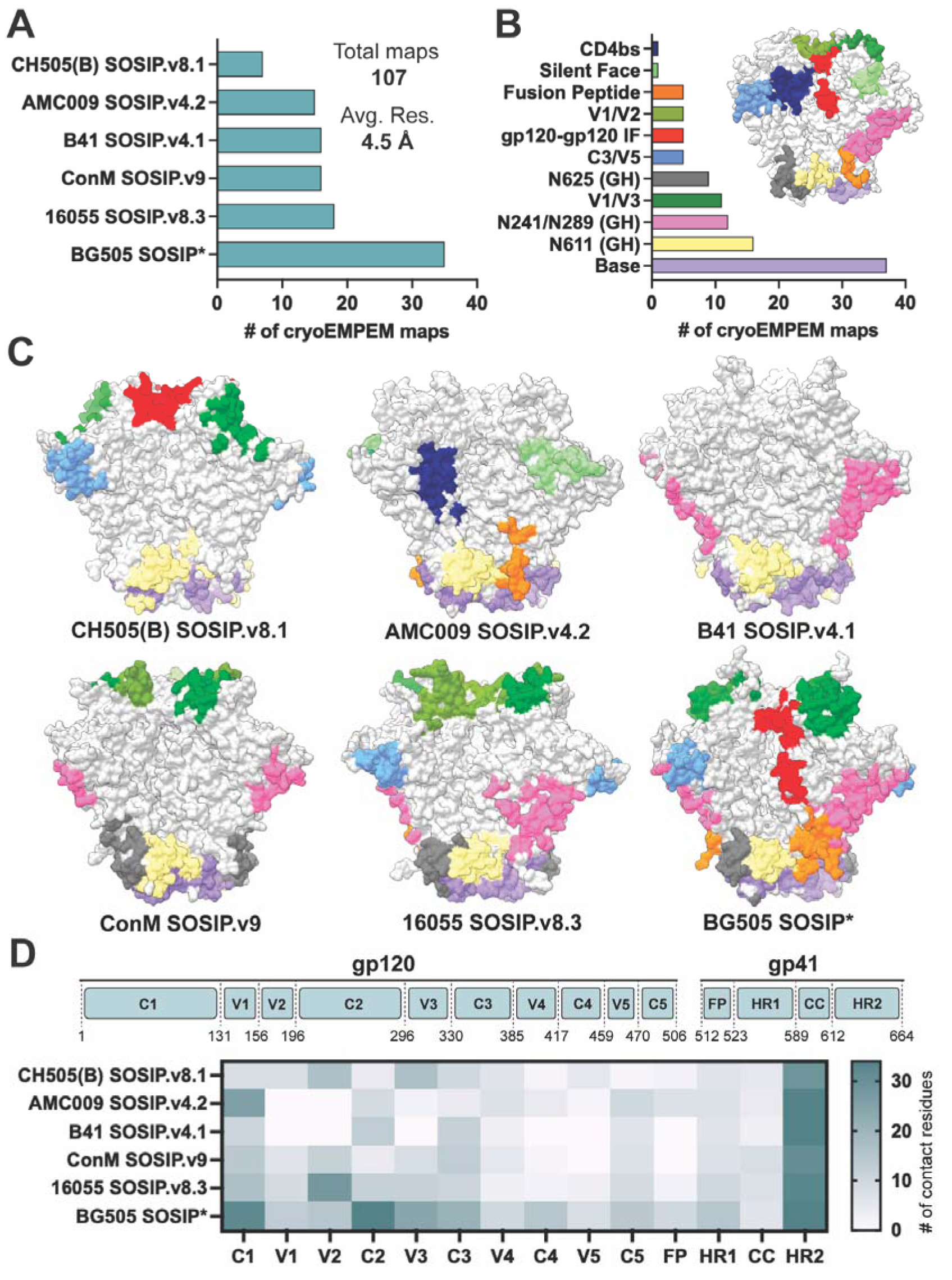
Overview of cryoEMPEM analyses and epitope specificity across HIV Env antigens. (**A**) Number of cryoEMPEM maps reconstructed for each Env antigen. (**B**) Classification of cryoEMPEM maps according to the epitope specificity of the recovered polyclonal antibodies. (**C**) Immunodominant epitopes across different HIV Env antigens. Epitopes are colored according to the scheme shown in panel B. N-linked glycans are omitted for clarity. (**D**) Heat map showing the number of non-redundant amino acid contacts mapped to distinct continuous regions of HIV Env. The coloring scheme is shown on the right. The corresponding amino acid ranges within gp120 and gp41 are indicated at the top.

Of the newly generated pAbC maps, 65 had global resolution below 8 Å and were of sufficient quality to perform antigen docking and modeling of the Ab backbones (**figs. S19-S23**), enabling identification of general epitope regions engaged by each antibody. Relevant model statistics are presented in **Extended data - Table 3**. Together with previously published BG505 SOSIP data, this yielded a combined set of 100 input models and maps (**Extended data - Table 4**). However, precise assignment of epitope contacts is hindered by the absence of amino acid side chains in the paratopes of reconstructed models, due to the inherent lack of sequence information for polyclonal antibodies. Therefore, standard tools for contact assignment based on atomic distances could not be used. To address this, we employed a surface-based approach in which the molecules are represented as discretized surface meshes, compartmentalizing the density map into regions corresponding to the antigen, Ab, and glycans. This strategy is adapted from MaSIF^50^, which uses MSMS software to generate solvent-accessible antigen surfaces (see **Methods**). Map density above a user-defined threshold was transformed into spatial coordinates, which were then used to assign epitope-paratope contacts, defined by a 3 Å distance cutoff between antibody coordinates and the nearest antigen surface vertex. This enabled automated, unbiased assignment of epitope contacts across all Env antigens (**Fig. 2C**).Structural analyses indicated that, for most Env constructs, large portions of the trimer surface remained subdominant and elicited antibody responses below the detection limit of cryoEMPEM (**Fig. 2C**). Antibody engagement was therefore restricted to a subset of epitope space, with partial overlap in the regions targeted across Env antigens. The most prominent examples of shared immunodominance were the Base and regions proximal to N611 and N625, which together accounted for more than 60% of reconstructed pAbCs (**Fig. 2B**). Antibodies targeting these sites converged on similar structural elements, primarily residues within HR1, HR2 and C1 elements (**fig. S24A**), engaging corresponding sets of amino acids in different Envs. For each epitope, the HxB2-numbered contacts are listed in **Extended data - Table 4**.

Other commonly targeted regions, including V1/V3, V1/V2, and C3/V5, exhibited only partial positional overlap in different antigens. In contrast to the Base and N611/N625 sites, these epitopes varied substantially across Envs due to differences in loop length, sequence composition, and distribution of potential N-linked glycosylation sites (PNGS), yielding distinct molecular surfaces for antibody engagement. For example V1/V3 epitope was not targeted in the context of AMC009 SOSIP.v4.2 or B41 SOSIP.v4.1, and was engaged via different amino acid contacts and angles of approach in other Envs (**Fig. 2C, fig. S24B**). Accordingly, antibodies elicited by the six antigens contacted different numbers and combinations of residues within constant and variable loops, yielding unique heat maps of epitope contacts (**Fig. 2D**).

Antibodies recognizing bnAb-like epitopes, including the CD4bs, SF, and the Fusion Peptide (FP), were also observed but more sporadically (**Fig. 2B**). Notably, AMC009 SOSIP.v4.2 elicited detectable responses to all three epitopes in multiple animals (**Fig. 1B, top**). Structural and sequence comparison of AMC009 with other Envs in this study revealed three unique mutations in the D-loop (residues 275-283), which is part of the CD4bs, and the absence of a PNGS at position 411, a glycan site that partially shields the SF epitope in other Envs (**fig. S24C**). Finally, the FP-targeting antibody appears to engage its epitope conditionally when the proximal N611 PNGS is unglycosylated, creating a glycan hole (GH). In all cases, local sequence and glycan differences contributed to formation of unique immunodominant epitopes in AMC009, that are absent in other antigens.

Altogether, the comparative analysis of epitope engagement indicated that antibody responses preferentially targeted a limited fraction of the Env antigenic space, while much of the trimer surface remained poorly targeted. Further, subtle sequence-level differences, including changes in PNGS distribution, were sufficient to measurably redirect antibody engagement across Envs. These findings motivated a comprehensive, quantitative analysis of molecular contacts and their relationship to immunodominance.

### Glycan context and amino acid composition impact epitope immunodominance

The conformational epitopes in Env are composed of surface-exposed residues drawn from multiple discontinuous sequence segments. We first assessed the properties of peptidic elements within immunodominant epitopes and compared them with subdominant antigenic sites, including bnAb-targeted epitopes from published structures (**Extended data - Table 4**). Analysis of antibody-antigen contacts showed that ∼90% of reconstructed pAbC targeting immunodominant epitopes interacted with ∼10-30 amino acid residues (**Fig. 3A**), consistent with prior structural analyses of conformational epitopes^51,52^. Further, these Abs largely avoided N-linked glycans, with ∼75% contacting only 0-4 monosaccharides (**Fig. 3B**). In contrast, bnAbs preferentially target conserved subdominant sites that are extensively shielded with glycans. Depending on the epitope, bnAbs engage up to 48 amino acids while contacting on average 5-9 monosaccharides and, in some cases, more than 15.

**Figure 3.**
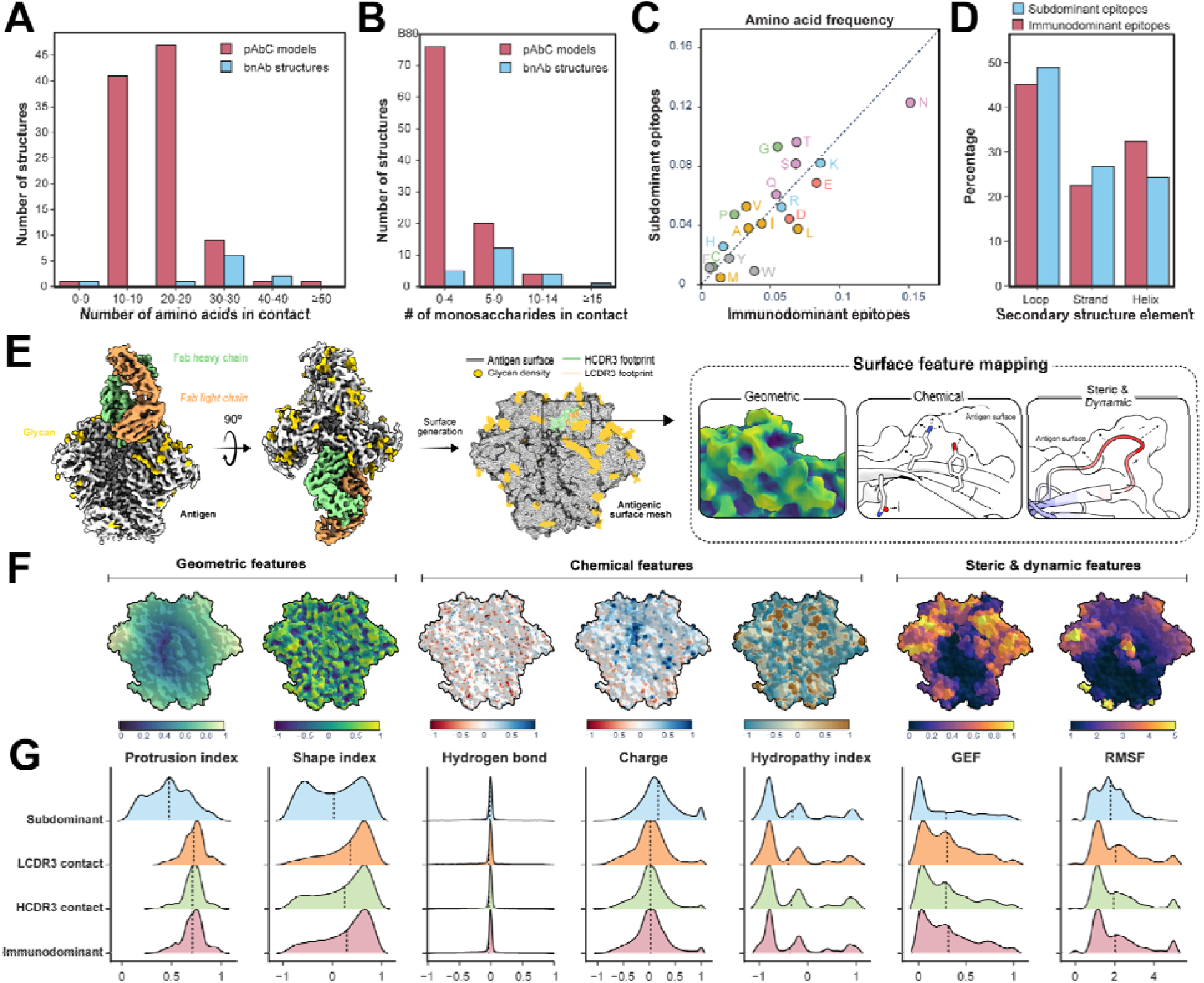
(**A**) Comparison in the number of amino acids in contact with the bnAb versus pAbC models. (**B**) Comparison in the number of monosaccharides in contact with bnAb versus pAbC models. (**C**) Differences in the frequency of 20 amino acid types at the residue level between immunodominant and subdominant sites. (**D**) Secondary structure elements of the immunodominant residue vs subdominant residues. (**E**) Computational framework for compartmentalizing density maps and generating surface features for HIV Env. (**F**) Examples of HIV Env surfaces colored by the values of corresponding geometric, chemical, and steric/dynamic features. (**G**) One-sided violin plots showing distribution of feature values among vertices, with the immunodominant epitope category further divided into contacts by LCDR3 (orange) and HCDR3 (green).

Next, we compared the amino-acid composition of immunodominant epitopes and subdominant surface regions across all tested antigens (**Fig. 3C**). Previous analyses of linear B-cell epitopes have shown that public antibody responses preferentially target viral peptides enriched in charged and aromatic residues^23^. Our data are broadly consistent with this trend but highlight a pronounced overrepresentation of hydrophobic residues (e.g., W, M, and L) within immunodominant sites, whereas charged (e.g., D, E) and polar residues (e.g., N) showed only moderate enrichment (**fig. S25**). In particular, W showed consistently higher abundance in immunodominant epitopes across all six Env antigens (**fig. S26**), likely reflecting its capacity to support diverse energetically favorable interactions, including π-π and cation-π stacking, hydrophobic contacts, and hydrogen bonding. In contrast, subdominant regions had higher content of smaller polar and hydrophobic residues (G, P, V, S, T), consistent with their lower average energetic contributions to antibody-antigen interactions^53,54^.

Despite differences in amino-acid composition and residue-specific contributions to secondary structure elements, we observed no major quantitative differences in the overall secondary structure content between immunodominant and subdominant sites (**Fig. 3D**). All detected epitopes contained loop (coil) regions, with variable contributions from α-helices and β-strands. The high loop content on the surface (∼45–50%) is a common feature of HIV Env and other viral antigens and is thought to facilitate immune escape^55–57^. Compared with helices and strands, loops exhibit greater tolerance to mutations, particularly at solvent-exposed positions with limited steric constraints^58^, allowing sequence variation with reduced impact on viral fitness. However, antibody responses directed against highly flexible loop regions (e.g., parts of V1-V4) may be less readily resolved by cryoEMPEM. Thus, epitopes dominated by highly flexible linear segments may be underrepresented in the current dataset.

Overall, per-residue contact analysis showed that antibodies targeting immunodominant epitopes engage ∼10-30 residues, predominantly in loop regions, and preferentially target sites with lower glycan density that are enriched for specific amino acids.

### The contribution of epitope geometry, dynamics, and chemical features

Next, we evaluated the geometric, chemical, and dynamic properties of Env epitopes using a surface-centric framework illustrated in **Fig. 3E**. Geometric features included the protrusion index (PI), for quantifying surface exposure at the domain level (i.e., distance from the center-of-mass), and the shape index (SI), which describes the local curvature. Chemical features characterized the availability of interaction-relevant groups, including hydropathy, continuum electrostatics, and hydrogen-bond donor/acceptor potential. To assess how local flexibility of peptidic and glycan elements influences epitope immunogenicity, we generated models of fully glycosylated Env antigens and performed molecular dynamics (MD) simulations^59^. From these, we extracted the Glycan Encounter Factor (GEF) and per-residue root-mean-square fluctuation (RMSF), representing steric hindrance by glycans and local flexibility, respectively. These values were mapped to the nearest surface vertices for all six antigens, with example visualizations for BG505 SOSIP MD39 shown in **Fig. 3F**. Using the epitope definitions from **Fig. 2C**, surface vertices were classified as immunodominant or subdominant and further partitioned by contacts to heavy- and light-chain CDR3 loops (HCDR3 and LCDR3). This comprehensive dataset enabled comparison of geometric, chemical, and dynamic feature distributions across categories based on modality and median values (**Fig. 3G**).

The most pronounced differences between immunodominant and subdominant sites were observed in geometric features (**Fig. 3G**, left). Immunodominant sites exhibited higher average PI values, indicating greater protrusion from the antigen center of mass. This is consistent with empirical observations from immunizations using multivalent Env displayed on nanoparticles^60,61^. In these studies, a regular array of Env trimers preferentially enhanced targeting of epitopes near the trimer apex, which are highly protruding when on nanoparticles, while reducing responses to epitopes proximal to the trimer base, which lie closer to the center of mass. Immunodominant epitopes also exhibited higher median SI values, with a single peak in the positive SI range, reflecting a predominance of convex surfaces. Notably, a subset of epitopes, including the Base (**fig. S27**), contained localized concave regions that were engaged by antibody HCDR3 loops. This is evidenced by a secondary peak at negative SI values among HCDR3-contacted surface vertices. Antibodies targeting these sites often had extended HCDR3 loops to access recessed surface pockets, as shown for Base-directed pAbCs in **fig. S28**.

Analysis of surface chemical features revealed comparable median values and distributions between immunodominant and subdominant sites (**Fig. 3G**, center), but substantial per-epitope variability was observed, particularly for electrostatic potential and hydropathy (**fig. S27**). On average, the immunodominant epitopes showed a modest shift toward more negative electrostatic values, consistent with enrichment of D and E residues, identified by contact analyses (**Fig. 3C**). Although immunodominant sites featured a greater content of W, L, and M, this was not reflected in bulk hydropathy scores, likely because averaged surface metrics dilute residue-specific contributions. Overall, these results indicate that chemical features alone do not strongly distinguish immunodominant from subdominant regions across the Env surface.

Finally, analysis of steric and dynamic features from MD simulations revealed differences in distribution modality without significant shifts in median values between immunodominant and subdominant epitopes (**Fig. 3G**, right). The RMSF distribution for immunodominant surface vertices showed a prominent peak at ∼1 Å, corresponding to conformationally stable regions. At the same time, all sites also included highly dynamic residues with RMSF > 2 Å, which become ordered upon antibody engagement. While it usually entails an entropic cost, this Ab-induced structuring effect is not uncommon, particularly at highly flexible epitopes such as the fusion peptide^62^. However, the most immunodominant epitopes, such as the Base, N611 (GH), and N625 (GH), are primarily composed of structured residues with RMSF < 2Å (**fig. S27**).

Surprisingly, GEF analysis showed that subdominant surface elements had median scores comparable to those of immunodominant epitopes, indicating similar levels of glycan coverage. Although this appears counterintuitive given the well-established role of glycan shielding in limiting epitope accessibility, several Env regions fail to elicit detectable antibody responses despite minimal glycan occlusion. The largest such region lies in the central portion of the molecule, including parts of the CD4bs, and comprises residues from C1 (46-84, 97-106) and HR1 (545-564), flanked by glycans at positions N262, N276, and N637 (**fig. S29**). These findings suggest a complex relationship between surface molecular features and epitope immunodominance that cannot be approximated by a single parameter.

A multivariate analysis integrating multiple surface features was therefore performed to identify non-obvious correlations associated with immunodominance (**fig. S30**). Two such comparisons revealed trends consistent with experimental observations. Bivariate analysis of PI and GEF showed that the majority of immunodominant epitopes exhibited high average protrusion (PI > 0.5) and low average glycan shielding (GEF < 0.5). In contrast, subdominant sites were typically associated with increased glycan density (GEF > 0.5) and/or reduced protrusion (PI < 0.5). In the latter case, proximity to the antigen center of mass likely reduced antibody engagement due to partial shielding by the protein’s quaternary structure. Further, joint evaluation of RMSF and PI further indicated that ∼90% of immunodominant surfaces combined protruding elements (PI > 0.5) with limited flexibility (RMSF < 1.5 Å). When these criteria were applied to the abovementioned C1/HR1 subdominant site, the exceptionally low glycan shielding (GEF = 0.01) may be offset by poor protrusion (PI = 0.39) and elevated flexibility (RMSF = 2.2 Å), providing a plausible explanation for the lack of detectable antibody targeting (**fig. S29**).

Other associations can also be observed through bivariate analyses, although with greater overlap in surface features between immunodominant and subdominant epitopes, making it challenging to unambiguously establish their relative contributions. Thus far, these data indicate that immunodominant epitopes are characterized by a combination of compositional, geometric, chemical, and dynamic properties, supporting the use of multivariate approaches to identify robust correlates of epitope immunodominance.

### Machine learning on antigenic surface features to predict immunodominance

To examine how surface features collectively relate to immunodominance and enable prediction, we developed a machine-learning framework that captures non-linear interaction patterns among antigenic surface properties. Input descriptors were constructed by concatenating features that exhibited distributional differences between surface labels (**Fig. 3**). These included two geometric features (SI and PI), glycan shielding (as distance to the nearest PNGS), chemical composition and properties (contacting amino acids). Using these descriptors as inputs, we trained an XGBoost classifier (REF) to differentiate between vertices labeled as immunodominant or subdominant. The model’s probabilistic output, termed the **A**ntigenic **S**urface **I**mmunodominance (ASI) score, represents a relative measure of the likelihood that a given surface site is immunodominant (**Fig. 4A**).

**Figure 4.**
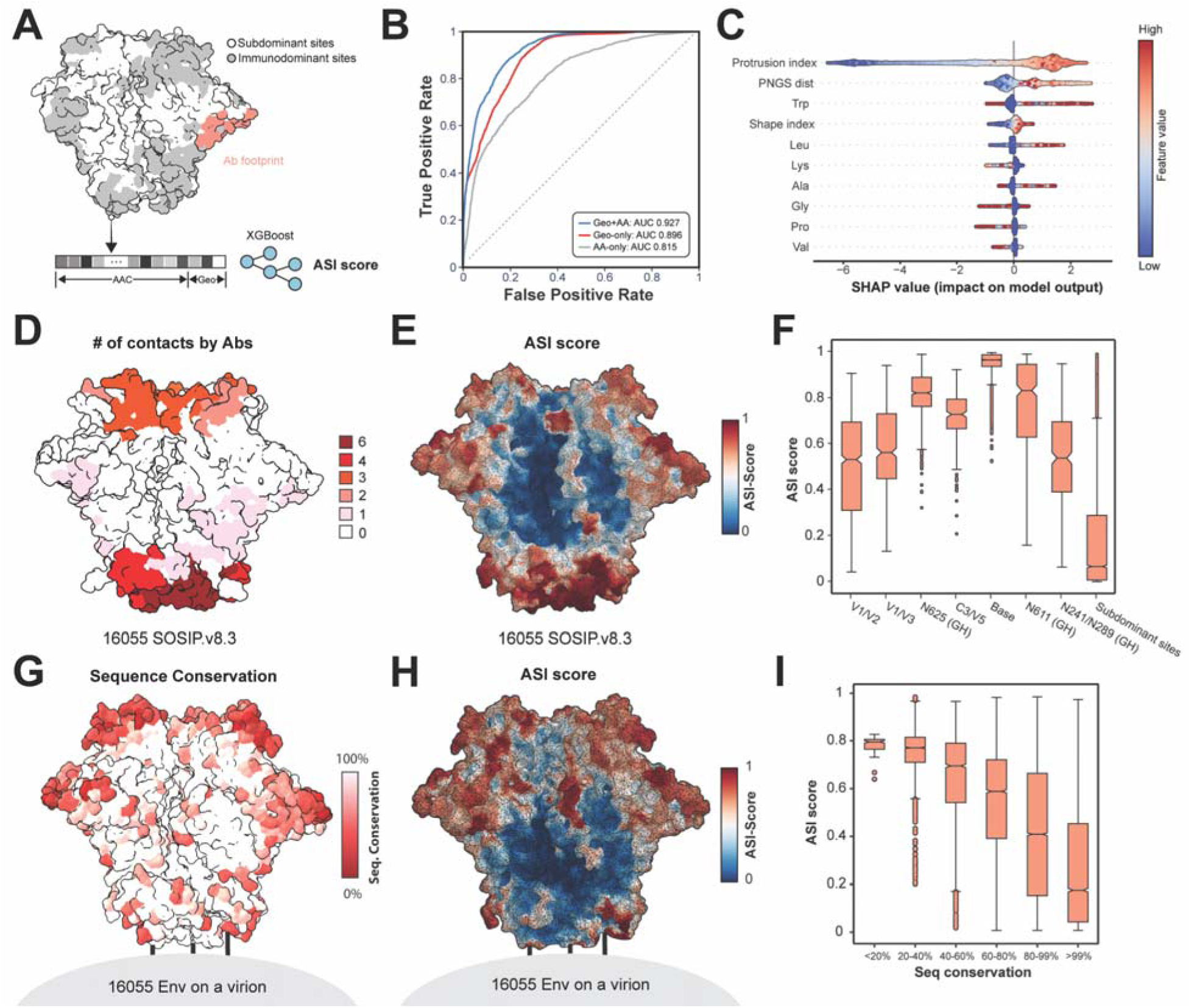
Predicting immunodominance with a machine learning model. (**A**) Schematic of XGBoost classifier training and ASI score prediction. (**B**) Receiver operating characteristic (ROC) curves for models trained using geometric features and amino acid composition combined, or each feature individually. (**C**) SHAP summary plot showing the distributions of the top 10 features ranked by importance. (**D**) 16055 SOSIP.v8.3 model with residues colored by the number of contacts with distinct pAbCs (n = 18). (**E**) ASI prediction mapped onto the full surface of the 16055 HIV Env model. (**F**) Distributions of ASI scores across immunodominant and subdominant sites in 16055 SOSIP.v8.3. Boxes indicate interquartile ranges; whiskers denote distribution limits. (**G**) Sequence conservation among 16055-like Indian-origin HIV strains mapped onto the model of the native 16055 ectodomain; the viral membrane is shown in gray. (**H**) ASI prediction for the 16055 HIV Env AlphaFold2 model coupled to a membrane-mimicking element. (**I**) Distribution of ASI scores from panel H across surface vertices stratified by sequence conservation (panel G). Boxes indicate interquartile ranges; whiskers denote distribution limits.

To benchmark the model’s performance, we tested it using the epitope features of the 16055 SOSIP.v8.3, the most evolutionarily distant Env antigen in the dataset (**fig. S1**). Model performance was evaluated using the receiver operating characteristic area under the curve (ROC AUC) to compare different input features. Notably, the model incorporating the geometric features, PNGS distance and amino acid composition achieved the highest predictive power on the test dataset (ROC AUC = 0.927, **Fig. 4B**). In contrast, models trained with only one of these feature types performed worse, as evidenced by their lower ROC AUC scores, indicating that all parameters are critical for accurate modeling of immunodominance.

To assess how individual features contribute to epitope targeting, we applied SHapley Additive exPlanation (SHAP)^64^. The top 10 features showing the strongest correlations are shown in **Fig. 4C** with the complete list in **fig. S31A**. Consistent with prior analyses, surface sites with higher PI values contributed positively to ASI scores, whereas lower PI values had the opposite effect. SHAP also revealed complex feature interactions. For example, when conditioning on SI, protrusion modulated rather than uniformly amplified the positive effect of convexity, such that less protruding convex sites (low PI, high SI) could contribute more than highly protruding ones (high PI, high SI; **fig. S31B**). The model further captured the influence of glycan shielding that aligned with empirical evidence. Protruded vertices more than 15 Å from the nearest PNGS contributed positively to ASI scores, whereas distances below 10 Å exerted a net negative contribution (**fig. S31C**). Additional positive contributors included higher local frequencies of specific amino acids, particularly W, L, and Y.

Next, we evaluated ASI score accuracy by comparing experimental and predicted values for 16055 SOSIP.v8.3, the test antigen. Residues were first colored by the number of distinct antibodies contacting them (**Fig. 4D**), providing a coarse, relative measure of targeting frequency in the structural dataset. Predicted ASI scores were then mapped onto the same antigen for comparison (**Fig. 4E**). The model is normalized to a 0-1 range, with higher values indicating greater likelihood of belonging to an immunodominant epitope. Overall, strong qualitative agreement was observed, as cryoEMPEM-defined epitopes co-localized with regions assigned higher predicted scores. Quantitative per-epitope comparison of ASI values across surface vertices is shown in **Fig. 4F**. Median scores for immunodominant epitopes ranged from 0.53 to 0.96, with the 75% of values above 0.3 in each case. The observed variability within individual epitopes is expected, reflecting heterogeneity in local surface properties. Subdominant regions exhibited markedly lower scores (median = 0.07), with 75% of values below 0.3, demonstrating that the algorithm reliably distinguishes immunodominant from subdominant epitopes. While the relative ranking of immunodominant sites was only partially captured, the model successfully predicted the Base and proximal N625 (GH) as highly immunodominant, despite underestimating the potential of apex-proximal epitopes (V1/V2 and V1/V3). This partial agreement is encouraging given the limited structural training dataset and the unique sequence features of 16055. We anticipate that expanding the training set will further improve resolution of inter-epitope relationships and refine hierarchical predictions.

Finally, we assessed whether ASI-mapped immunodominant epitopes could identify major immune pressure points on Env. Clade C HIV Env sequences of Indian origin were retrieved from the LANL^67^ database, selecting sequences isolated after the original 16055 virus and sharing ≥95% identity within Env. For the 30 sequences meeting these criteria, per-residue conservation analysis revealed the V1, V2, and V4 loops at the trimer apex as the most variable regions, consistent with strong immune selection (**Fig. 4G**). ASI prediction was then applied to a model of the native 16055 Env ectodomain coupled to a membrane-mimicking surface element (**Fig. 4H**). Incorporation of the membrane mimic led to markedly reduced ASI scores at base-proximal epitopes compared to 16055 SOSIP.v8.3, showing that the algorithm captured the shielding effect of the viral membrane despite not being trained on this feature. Regions with high ASI scores co-localized with areas of low sequence conservation, as supported by the quantitative analysis in **Fig. 4I**. Surface vertices adjacent to residues with the lowest conservation exhibited the highest median ASI scores, whereas more conserved regions showed progressively lower values. The ASI value distribution was particularly narrow in the most diversified regions (conservation < 40%), showing high predictive precision. Together, these findings suggest that ASI-mapped immunodominant epitopes can be used to infer regions that are most likely to mutate under Ab-mediated immune pressure.

### Predicting immunodominance in Env antigens after glycan engineering

Glycan engineering through targeted removal or addition of PNGS sites is a well-established strategy to modulate epitope exposure across diverse viral antigens^12–14,68,69^. In the context of HIV Env, this approach has been successfully implemented in multiple vaccine candidates, including BG505 SOSIP GT1.1^70^, eOD-GT8^71^, RC1^72^, and ApexGT^73^, often as part of efforts to enhance the elicitation of precursor Abs against conserved epitopes. Here, we assessed the predictive power of the ASI by evaluating its ability to identify newly exposed epitopes generated by glycan remodeling, using BG505 SOSIP GT1.1 as a model. This construct promotes CD4bs-directed responses through removal of surrounding glycans at positions N185e, N185h, N197, N276, N386, and N462, together with sequence modifications, including substitutions within residues 276-281 of the D-loop and deletion of residues 185d–187 in the flexible V2 loop (**Fig. 5A**). ASI predictions were performed on BG505 SOSP with and without GT1.1 mutations using AlphaFold2-generated models. To account for local confidence in epitope conformation, we incorporated an option to scale the ASI values using the predicted local distance difference test (pLDDT)^65^ which penalizes low-confidence regions from being assigned high ASI scores based on potentially wrong conformations (See **Methods**). Comparison of ASI scores before and after GT1.1 modifications revealed marked local increases at the sites of removed glycans (**Fig. 5B**). The median ASI score around the D-loop rose to 0.72 and extended across a broader CD4bs surface, predicting this region as a favorable antibody target upon immunization.

**Figure 5:**
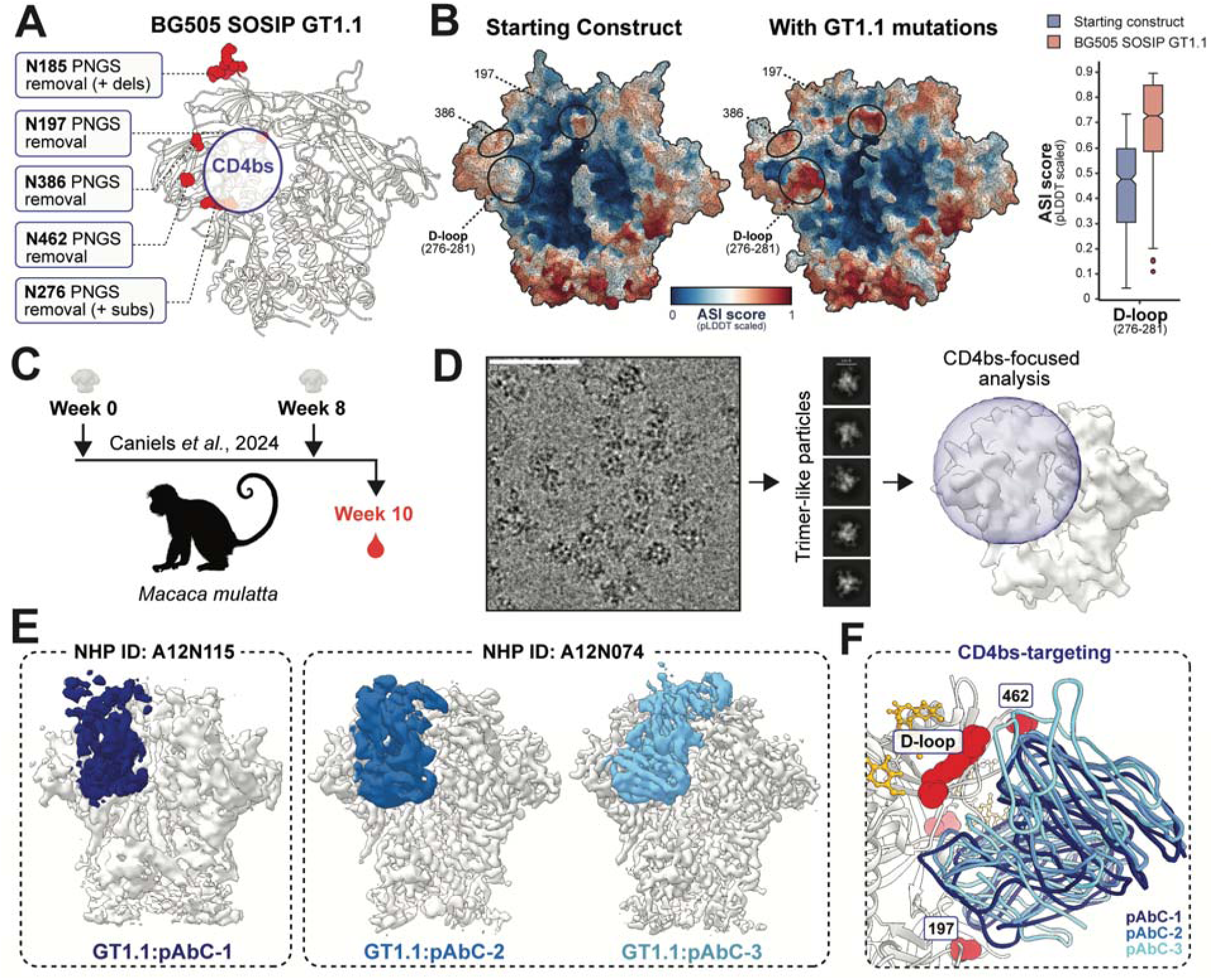
(**A**) Glycan remodeling sites in BG505 SOSIP GT1.1. (**B**) Predicted pLDDT-scaled immunodominance scores for BG505 SOSIP before and after GT1.1 mutations. (**C**) Schematic of macaque immunization with BG505 SOSIP GT1.1 as previously described^74^. (**D**) Workflow for structural analysis of CD4bs antibodies in non-human primates (NHPs). Left: representative micrograph and 2D classes of immune complexes with A12N0074 polyclonal antibodies. Right: reference map and spherical mask used for classification. (**E**) Electron microscopy reconstructions of CD4bs-targeting pAbC from macaque sera. Antigen is shown in gray, and Abs in shades of blue. (**F**) Overlay of pAbC structures (blue) bound to BG505 SOSIP GT1.1 (gray), with glycans shown in yellow and GT1.1 mutations as red spheres.

Previous studies have confirmed the capacity of the BG505 SOSIP GT1.1 antigen to induce Abs to the CD4bs epitope in macaques and humans^70,74^. To test if these Abs engage the local area predicted to be immunodominant, we selected polyclonal Ab samples from two rhesus macaques, immunized with this antigen as part of a previously completed pre-clinical evaluation of BG505 SOSIP GT1.1^74^. Each animal received 2 doses of the antigen at weeks 0 and 8, and immune sera were harvested at week 10 (**Fig. 5C**). Polyclonal Abs from these samples were purified, cleaved into Fab fragments, and complexed with the BG505 SOSIP GT1.1 antigen as described in the **Methods**. These immune complexes were subjected to cryoEMPEM analysis focusing on the CD4bs region (**Fig. 5D**). This approach allowed the recovery of three CD4bs-directed antibodies (**Fig. 5E**); one pAbC class binding to the CD4bs using antibodies from animal A12N115 (GT1.1:pAbC-1) and two structurally distinct pAbC fragments using antibodies from animal A12N0074 (GT1.1:pAbC-2 and pAbC-3). The representative EM data, including the corresponding FSC curves are shown in **fig. S32**.

All three antibodies bound the CD4bs epitopes, directly engaging residues 276-282 of the D2 loop (**Fig. 5F**), consistent with regions predicted to have favorable ASI scores. GT1.1 :pAbC-3 formed the most extensive contacts with this region via its HCDR3 and additionally interacted with C4/V5 residues 457-466, adjacent to the location of the mutated N462 glycan site. By contrast, GT1.1:pAbC-1 and GT1.1:pAbC-2 displayed inverted heavy and light chain orientations, spanning a broader CD4bs surface that included C5 and C1 elements and contacting residues 195-198 opposite the D2 loop (**fig. S33A-C**). Notably, these two antibodies exhibited highly similar backbone conformations (RMSD = 1.56 Å) and binding footprints, closely resembling the monoclonal antibody 21N13 (PDB: 8SW4) isolated from the same cohort of rhesus macaques^74^ (**fig. S33D**). This convergence indicates that the macaque antibody repertoire includes a relatively abundant class of precursors that can be efficiently driven toward CD4bs recognition by GT1.1 immunization, provided that impeding glycans are removed.

Collectively, these results demonstrate that the ASI framework enabled identification of changes in surface immunodominance arising from extensive N-linked glycan remodeling combined with minor sequence alterations. Importantly, the model extrapolated relationships between PNGS positioning and Ab targeting to identify newly exposed epitopes whose glycan configurations were not represented in either the training or test datasets.

### Engineering immunodominant epitopes through tailored amino acid composition

An additional insight from the combined structure meta-analysis (**Fig. 3D**) and SHAP interpretation of the XGBoost model (**Fig. 4C**) was that enrichment of certain amino acids, such as W, correlates with increased epitope engagement probability. We therefore hypothesized that introducing residues with favorable properties into otherwise subdominant epitopes could lead to detectable induction of response. To test, we engineered BG505 SOSIP with immunofocusing (IF) mutations within the CD4bs and FP epitopes. Candidate substitutions were derived from naturally occurring HIV Env sequence variants (LANL database^67^) and screened using an in-house script to identify segments enriched in W, Y, L, and charged residues. Selected segments were introduced into BG505 SOSIP, which was further modified with stabilizing mutations and glycan insertions at BG505-specific immunodominant sites to enhance native-like trimer content and reduce autologous reactivity. The full set of mutations incorporated into the resulting construct, termed BG505 SOSIP IF, is summarized in **Table S3**, with the positions of the IF substitutions mapped onto the Env trimer model in **Fig. 6A**.

**Figure 6.**
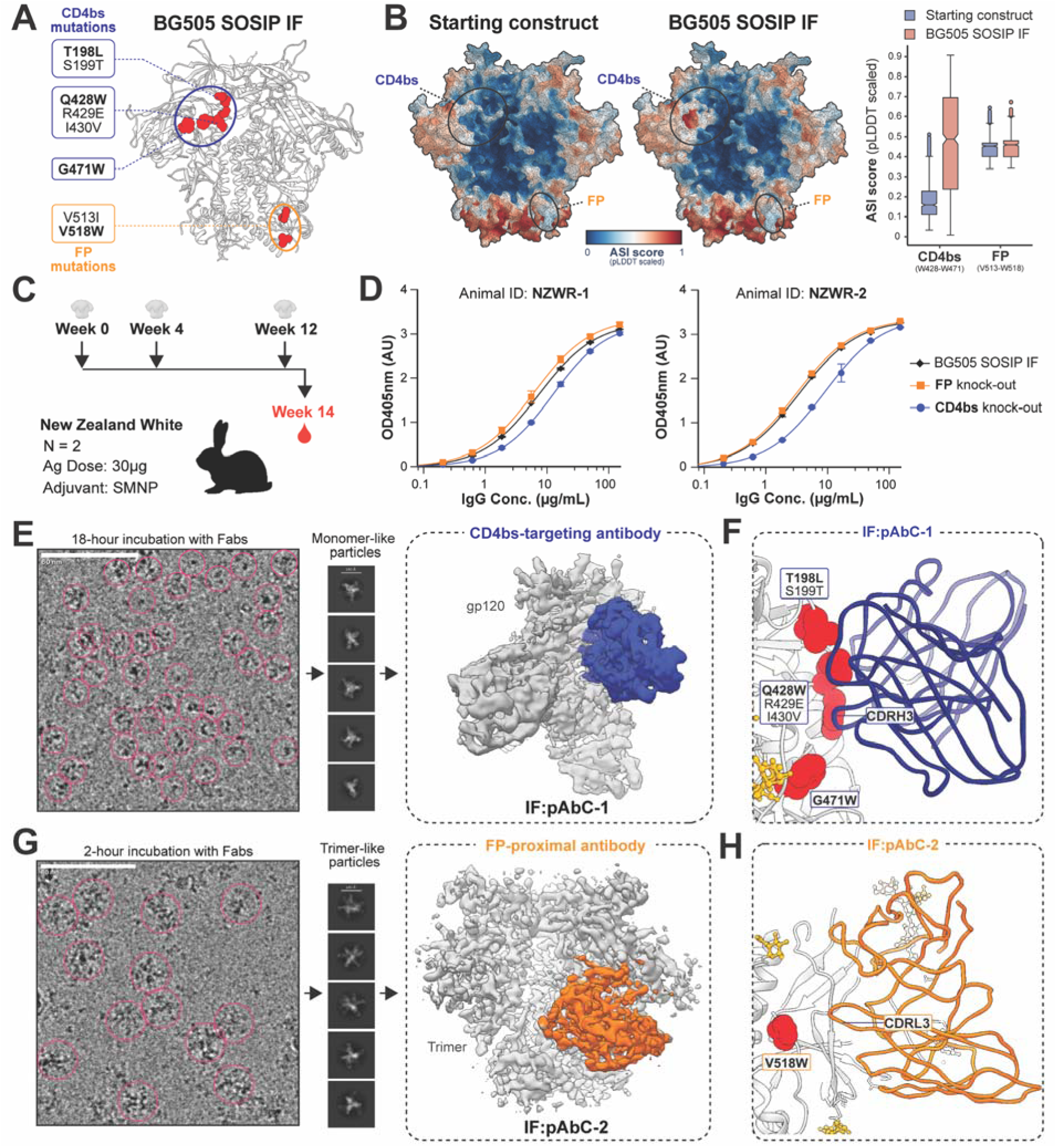
(**A**) Immunofocusing substitutions introduced into BG505 SOSIP IF; the CD4bs and FP are shown in blue and orange, respectively. (**B**) Predicted pLDDT-scaled ASI scores for BG505 SOSIP before and after immunofocusing mutations. (**C**) Schematic of rabbit immunization with BG505 SOSIP IF. (**D**) ELISA binding of rabbit polyclonal sera to BG505 SOSIP IF (black) and to revertant antigens in which CD4bs (blue) or FP (orange) mutations were restored to wild type. (**E**) Representative micrograph and 2D class averages of immune complexes purified after a standard, 18 h incubation, with reconstructed EM map showing pAbC binding to the CD4bs. Magenta circles in micrographs mark the particles. (**F**) Backbone structure of a CD4bs-targeting polyclonal antibody bound adjacent to the mutated FP residue (IF-pAbC-1). (**G**) Representative micrograph and 2D class averages of BG505 SOSIP IF immune complexes purified after 2 h incubation, with reconstructed EM map showing an antibody bound near the FP (orange). Magenta circles in micrographs mark the particles. Note the difference in particle size. (**H**) Backbone structure of IF:pAbC-2 (orange) with the immunofocusing mutation shown as red spheres.

Structural models of BG505 SOSIP with and without the IF mutations were generated using AlphaFold2^65^. The immunodominance probability for each surface site was then evaluated using the ASI framework (**Fig. 6B**). Comparison of the predicted scores for these two models revealed a localized increase at the interface of the newly introduced W428 and W471 residues in the CD4bs, indicating a higher probability of antibody engagement. In contrast, the predicted scores for the FP epitope were largely unchanged, likely reflecting the high flexibility of the fusion peptide (low pLDDT score) and significant content of hydrophobic residues, which limits the impact of amino-acid composition modifications on its immunodominance.

The BG505 SOSIP IF construct was expressed and purified as described in the **Methods**. nsEM analysis showed that >98% of the sample consisted of native-like trimers (**fig. S34A**). Monoclonal antibody binding was assessed by biolayer interferometry (BLI; **fig. S34B**), revealing preserved interactions with glycan supersite and V2-apex epitopes, as indicated by comparable binding to 2G12 and PGT145 bnAbs. In contrast, bnAbs targeting the CD4bs (e.g., VRC01, CH103, and CH235) and FP (e.g., VRC34) exhibited reduced binding, showing that although the IF substitutions came from native HIV sequences, they altered these epitopes in the BG505 context. For further evaluation of antigenicity, we performed glycan composition analyses by liquid chromatography-mass spectrometry (LC-MS; **fig. S34C**). These data confirmed >95% occupancy at PNGS surrounding the CD4bs (N197, N276, N386) and FP (N88, N611, N637), with a mixture of oligomannose- and hybrid-type glycans comparable to BG505 SOSIP constructs studied in the past^16,75^. These results indicate that the engineered epitopes maintain native-like glycan shielding, providing an optimal context to evaluate the influence of substituted amino acids on epitope immunodominance.

Next, we proceeded to evaluate the immunogenic behavior of BG505 SOSIP IF and determine if the engineered amino acids elicit a measurable antibody response. Two New Zealand White rabbits were immunized with this antigen co-formulated with the SMNP adjuvant^75,76^ at weeks 0, 4, and 10, and serum samples were collected at week 12 (**Fig. 6C**). Polyclonal IgG was isolated from the serum and analyzed by enzyme-linked immunosorbent assay (ELISA) to quantify antigen-specific responses (**Fig. 6D**). Both animals developed antibodies binding BG505 SOSIP IF with midpoint titers (EC_50_) of 7.4 ± 0.6 µg/ml and 3.5 ± 0.3 µg/ml for NZWR-1 and NZWR-2, respectively. When tested against a construct in which the CD4bs substitutions were reverted to the native BG505 sequence, EC_50_ values increased ∼2-3 fold, indicating that a fraction of the antibody response depends on these mutations, and consistent with engagement of the CD4bs. In contrast, no significant change in EC_50_ or maximum absorbance was observed when the FP substitutions were reverted, consistent with the absence of a substantial FP-specific response.

Polyclonal antibodies from both animals were pooled, digested into Fab fragments, and complexed with BG505 SOSIP IF using the same protocol as with other samples, including 18-hour incubation for assembly. The complexes were imaged on a Titan Krios (**Extended data - Table 1**), and processed as shown in **fig. S35A**. Analysis of micrographs and 2D classes indicated that the great majority of particles were smaller than expected for a BG505 SOSIP IF trimer (**Fig. 6E**). 2D-cleaned particles yielded a 3D map of gp120 bound to four antibodies. Despite the modest resolution of 4.8 Å, the map of sufficient quality to enable reliable model docking and reconstruction of antibody backbones (**fig. S36A,B**). Analysis of the refined model revealed that one antibody, IF:pAbC-1, engaged the CD4bs via its HCDR3 loop (**Fig. 6F**), with bridging EM density extending toward the engineered elements W428-I430V and L198-T199 (**fig. S36C**). The antibody approaches the CD4bs at an angle predicted to sterically clash with a neighboring gp120 protomer (**fig. S36D**). Further, its epitope footprint partially overlaps that of b12 (**fig. S36E**), a CD4bs bnAb known to induce Env opening and trimer destabilization in certain contexts^77,78^. We believe that binding of IF:pAbC-1 and related CD4bs antibodies within the polyclonal response contributed to the extensive trimer disassembly and accumulation of gp120 monomers observed in cryoEMPEM analyses. Thus, inclusion of IF mutations to the otherwise subdominant CD4bs was sufficient to recruit B-cell binding and induce detectable antibody response.

From these data, reliable reconstructions containing gp41 could not be obtained, precluding structural analysis of FP-specific antibodies. To reduce the observed trimer disassembly, immune complexes were prepared with a shorter 2-hour incubation prior to purification and grid application, which was previously shown to limit antibody-induced Env destabilization^80^. This second dataset showed a higher proportion of trimer-Fab complexes in EM images (**Fig. 6G**), allowing application of the standard cryoEMPEM workflow used for other immune complexes (**fig. S4**) to evaluate whether FP- or CD4bs-directed antibodies were present. Analysis revealed no CD4bs antibodies bound to intact trimers; however, several antibodies binding proximal to the FP region were identified (**fig. S35B**). Examination of the maps and models showed that their contacts were primarily with HR1/HR2 and C1/C2 elements associated with the N241/N289 (GH), N611 (GH), and Base epitopes (**fig. S37A**). Among them, IF:pAbC-2 approached the engineered FP residue, W518, within ∼5 Å (**Fig. 6H**), but this interaction appears structurally non-essential for binding, consistent with the ELISA results. Importantly, in the model this residue is packed against HR2, leaving it partially occluded within the trimer core (**fig. S37B**). This restructuring of the normally exposed FP epitope may have reduced its accessibility to antibodies compared to native BG505 SOSIP, providing a plausible explanation for the absence of a detectable response.

Collectively, these proof-of-concept experiments demonstrate that the ASI framework can be used to modulate Env immunodominance through targeted changes in amino-acid composition. At the same time, they highlight the importance of structural context, showing that engineered residues must remain exposed to influence antibody targeting and that highly flexible epitopes present additional design constraints.

## DISCUSSION

Characterizing the biases imposed by B-cell immunodominance can inform viral evolution and support rational vaccine design, yet the underlying molecular determinants remain poorly defined due to insufficient and suboptimal structural data. To address this gap, we applied cryoEMPEM and assembled a library of high-resolution HIV Env structures spanning multiple clades, in complex with polyclonal antibodies bound to immunodominant epitopes. We generated 13 datasets yielding 71 new structures and combined them with published BG505 SOSIP entries, resulting in 106 structures across the training, testing, and validation sets. Data collection and initial processing were relatively rapid (∼1–2 weeks per dataset), whereas model building, refinement, and validation remain rate-limiting; though continued development of ML-assisted tools^80^ is expected to accelerate these steps. This collection enabled systematic mapping of immunodominant sites and extraction of shared molecular features across antigenic settings, effectively overcoming limitations of available PDB data. In this context, our results demonstrate that current structural biology workflows can support the assembly of targeted structure libraries tailored to specific experimental questions.

Surface-focused meta-analysis of the immunodominant epitope space in our structural library revealed factors contributing to efficient antibody recruitment. The analysis pointed towards the fundamental importance of geometric properties, quantified through protrusion and shape metrics, while also confirming the established roles of glycan shielding^5,13^ and amino acid composition^82,83^. Intuitively, the importance of geometry can be understood using sites that are recessed or buried, which restrict antibody approach angles or impose requirements for longer CDR loops, collectively reducing the likelihood of targeting. However, this principle also addresses the observed preferential targeting of apex-proximal epitopes on viral glycoproteins (e.g., the influenza HA head domain) during natural infection or vaccination with virus- or nanoparticle-displayed antigens^17,49,84,85^. Apex-proximal sites are highly protrusive due to outward orientation on a particle, unlike isolated recombinant antigens, where apex and base protrusion are equivalent, resulting in less polarized antigenic landscape. The protrusion and shape index parameters can partially explain and quantify this geometric effect. However, the collective data indicated that immunodominance arises from the combined effects of geometry, chemistry, and glycosylation, with no single factor fully explaining the observed hierarchy of antibody targeting. The tailored nature of this dataset allowed direct comparison of immunodominant and subdominant sites, enabling estimation of their relative contributions and the derivation of a generalizable model.

To estimate the relative contributions of distinct structural and molecular features to immunodominance, we developed the ASI framework, which incorporates an XGBoost classifier trained on a cryoEMPEM-derived structural dataset. The model was rigorously validated using complementary interpretability tools (e.g., SHAP) and by testing its predictive performance across diverse antigens, including cases involving targeted glycan remodeling and amino-acid substitutions. Because the features used for training capture general properties of antigen surfaces, the approach is not limited to HIV Env and, in principle, applies to other viral antigens such as influenza, betacoronaviruses, and pneumoviruses, although its broader utility will require further validation. Existing computational approaches range from sequence-based linear epitope predictors to more advanced ML methods for protein-protein interaction prediction that leverage antigen structures and antibody sequence data^86–91^. While these tools can identify potential binding sites for individual monoclonal Abs with variable accuracy^92^, they are poorly suited to modeling polyclonal responses. In contrast, our ASI framework enables the identification of primary immune pressure points on viral antigens and supports *in silico* antigen engineering and the prediction of epitope targeting prior to experimental testing *in vitro* or *in vivo*. As such, it may improve early engineering and filtering of immunogens thereby reducing the number of animals used for preclinical evaluation of vaccine candidates.

This work introduces a new principle for engineering immunodominant epitopes through targeted modulation of amino acid composition. Although conceptually intuitive, this strategy has not been explored previously, in part due to a limited understanding of how residue composition influences immunodominance. Using either experimental structures or AlphaFold2 models, the ASI framework predicts how amino acid substitutions alter surface antigenicity and enables rational, targeted antigen design. While outcomes can be less predictable in highly flexible regions such as the fusion peptide, predictions were robust for structurally constrained epitopes, such as the CD4 binding site, where substitutions directly enhanced antibody engagement. Antigens engineered solely by amino acid composition can be used to prime responses toward the CD4bs and other bnAb epitopes without altering the glycan shield. These primed responses may then be guided toward broader cross-reactivity through stepwise reversion of engineered residues toward consensus sequences. Compared with reintroducing glycans, reversing point mutations is less disruptive to epitope-paratope interactions, potentially enabling more efficient and reliable shepherding of antibody responses. Therefore, the ASI model is complementary and applicable to ongoing HIV vaccine design efforts based on germline targeting^93^.

One limitation of the study was that our approach was antigen-centric and largely agnostic to the B-cell repertoire. Nevertheless, we analyzed samples from rabbits and macaques, in which prior studies have shown that immunodominant epitope hierarchies for HIV Env and other antigens closely resemble those observed in humans^47–49^. This consistency supports the idea that epitope-intrinsic properties (e.g., surface presentation, local topology, and favorable chemical features) are major drivers of immunodominance. At the same time, the underlying B-cell repertoire and its capacity to mature high-affinity antibodies against specific epitopes can modulate how individual sites are engaged, or whether they are targeted at all. A clear example is the mouse model, which has a more limited naïve repertoire and a reduced likelihood of generating the long HCDR3 loops required to engage heavily glycosylated antigens such as HIV Env^94,95^. Dissecting the interplay between epitope features and responding B-cell clones remains challenging due to the scale of the humoral response, but continued advances in cryoEMPEM-based antibody sequencing^96,97^ may make this increasingly tractable. We hope to incorporate additional species- and repertoire-based elements into future implementations of the ASI model.

## MATERIALS AND METHODS

### Clones and mutagenesis

Genes encoding CH505(B) SOSIP.v8.1, 16055 SOSIP.v8.3, B41 SOSIP.v4.1, AMC009 SOSIP.v4.2, ConM SOSIP.v9, and BG505 SOSIP GT1.1 were subcloned into PPI4 or pcDNA3.4 expression vectors. Sequence alignments of these constructs are shown in **fig. S1**. BG505 SOSIP IF (as well as the CD4bs/FP epitope revertants) were derived from BG505 SOSIP MD39 using site-directed mutagenesis (QuickChange Lightning, Agilent) and restriction-ligation cloning (restriction enzymes and Quick Ligation Kit, NEB) according to the manufacturers’ protocols. All sequences were verified by Sanger sequencing (Genewiz), and descriptions of the introduced immunofocusing (IF) mutations are provided in **Table S3**.

### Antigen expression and purification

All antigen constructs were produced as described previously^16^. Briefly, constructs were transfected into FreeStyle 293F cells using polyethyleneimine (PEI; Polysciences). For each liter of cells (0.8-1.2 × 10^6^ cells/mL), 250 µg of construct plasmid and 125 µg of Furin-encoding plasmid were used. Cells were cultured in FreeStyle 293 Expression Medium (ThermoFisher) for 6 days, then removed by centrifugation (9000 rpm, 1 h, 4°C). The supernatant was passed over an in-house-made PGT145 immuno-affinity column, extensively washed, and eluted with 3 M MgCl_2_, pH 7.4, into TBS buffer (20 mM Tris-HCl, 150 mM NaCl, pH 7.5). Eluate was concentrated using Amicon 100 kDa filters (Millipore) and buffer-exchanged into TBS. Antigens were then purified by size-exclusion chromatography (SEC) on a HiLoad 16/600 Superdex pg200 (Cytiva) equilibrated in TBS. Samples were reconcentrated to ∼1 mg/mL, aliquoted, flash-frozen in liquid nitrogen, and stored at -80°C. Protein concentration was measured using a NanoDrop spectrophotometer (Thermo Scientific) with E1% values calculated from each construct’s primary sequence (ProtParam).

### Monoclonal antibodies

Monoclonal Abs (e.g., PGT145, PGT151, 2G12) were produced by co-transfecting AbVec plasmids encoding the heavy and light chains into CHO cells cultured in ProCHO 5 medium (Lonza Biosciences). For each transfection, 500 µg of heavy chain and 250 µg of light chain plasmid were combined with a threefold mass excess of PEI and applied to 1 L of cells at ∼ 1 x 10^6^ cells/ml density. After 6 days, cells were removed by centrifugation (9000 rpm, 1 h, 4°C), and Abs were purified from the supernatant using CaptureSelect IgG-Fc (ms) resin (ThermoFisher). Column-bound Abs were eluted with 0.1 M glycine buffer (pH 3.0) and immediately neutralized with 1 M Tris-HCl (pH 8.0). Samples were concentrated using Amicon ultrafiltration units (30 kDa cutoff) and buffer-exchanged into TBS. Ab concentrations were determined with a NanoDrop spectrophotometer (Thermo Scientific) using E1% values calculated from the primary sequence (ProtParam). Aliquots at ∼1 mg/mL were stored at -80°C until use.

### Origins of serum and plasma samples

Serum and plasma samples used in this study were obtained from three primary sources. [1] For the initial characterization of immunodominant surfaces, serum samples were collected from previously completed immunization studies involving rabbit and rhesus macaque models. Details of the corresponding studies, sampling time points, and animal identifiers are provided in Table 1 and Table S2. [2] Polyclonal plasma samples from BG505 SOSIP GT1.1 immunizations were obtained from a previously completed rhesus macaque study^74^. Samples were collected at week 10, following two immunizations with BG505 SOSIP GT1.1. Animals A12N115 and A12N074 were selected based on preliminary screening indicating the presence of CD4bs-directed Abs. [3] Serum samples from rabbits immunized with BG505 SOSIP IF were generated specifically for this study; immunization protocols and sampling time points are described below.

### Isolation of polyclonal IgG from serum/plasma

Polyclonal Abs (IgG) were purified from serum or plasma using CaptureSelect IgG-Fc multispecies affinity resin (Thermo Fisher Scientific). After binding at 4°C under a flow of 0.5 ml/min, the column was washed with PBS (137 mM NaCl, 2.7 mM KCl, 10 mM Na₂HPO₄, 1.8 mM KH₂PO₄, pH 7.4), and bound Abs were eluted with 0.1 M glycine (pH 3.0), followed by immediate neutralization with 1 M Tris-HCl (pH 8.0). Eluted fractions were pooled and buffer-exchanged into PBS using Amicon centrifugal filters (30 kDa cutoff; Millipore). Ab concentrations were measured by spectrophotometry (NanoDrop Lite, Thermo Scientific). Starting material typically consisted of 2-5 mL of serum or plasma, yielding approximately 0.5-8 mg of polyclonal IgG per mL of input sample.

### Production of cleaved polyclonal Fab/Fc

IgG digestion was performed as previously described^39^. Briefly, IgGs were digested with crystalline papain (Sigma) for 5 h at 37 °C. Papain was first activated by incubation with 10 mM cysteine for 15 min at 37 °C in Tris-EDTA buffer (100 mM Tris-HCl, pH 7.5, 2 mM EDTA) at 1 mg/mL. Activated papain was then added to IgG in the same buffer containing cysteine, resulting in final concentrations of 20 µg/mL papain and 0.5 mg/mL IgG. The reaction was quenched by the addition of iodoacetamide to 30 mM. Fab/Fc fragments were separated from undigested IgG and papain by size-exclusion chromatography using a HiLoad 16/600 Superdex pg200 column (Cytiva) running in PBS. Fractions corresponding to Fab/Fc were pooled and concentrated to 2-4 mg/mL using Amicon centrifugal filters (10 kDa cutoff; Millipore). Typical recoveries ranged from 50% to 70% of the input IgG.

### Negative-stain-EM-based polyclonal epitope mapping (nsEMPEM)

Negative-stain electron microscopy (nsEM) was used to screen polyclonal immune complexes and identify optimal candidates for high-resolution cryoEM analysis. For 16055 SOSIP.v8.3, ConM SOSIP.v9, and BG505 SOSIP (multiple versions), nsEMPEM datasets were available from prior studies corresponding to the original immunization experiments^16,17,26^. Screening of B41 SOSIP.v4.1, AMC009 SOSIP.v4.2, and CH505(B) SOSIP.v8.1 immune complexes was performed in this study. Immune complexes were assembled by incubating 1 mg of purified polyclonal Fab/Fc from each animal with 20 µg of the corresponding SOSIP antigen for ∼18 h at room temperature. Complexes were separated from excess Fab by size-exclusion chromatography using a Superdex 200 Increase 10/300 column equilibrated in TBS, concentrated using Amicon ultrafiltration units (10 kDa cutoff), and immediately applied to carbon-coated 400-mesh Cu grids that had been glow-discharged for 30 s at 15 mA. Samples were adjusted to 50-100 µg/mL prior to grid application and stained with 2% (w/v) uranyl formate for 60 s.

nsEM data collection was performed as described previously^26^ using a Tecnai F20 transmission electron microscope operating at 200 kV. Images were recorded at a nominal magnification of 62’000, corresponding to a pixel size of 1.77 Å, with an electron dose of 25 e^-^/Å^2^ and a defocus of -1.5 µm. Micrographs were collected using a Tietz TemCam-F416 4k × 4k CMOS camera under control of the Leginon automated data acquisition software^98^. For each polyclonal immune complex (animal), 200–300 micrographs were collected, yielding five datasets per antigen for B41 SOSIP.v4.1, AMC009 SOSIP.v4.2, and CH505(B) SOSIP.v8.1, because there were five animals per group. Initial image processing, including particle picking and extraction, was performed using Appion^99^. Datasets were required to contain >100,000 particles to enable consistent comparisons; additional micrographs were collected if necessary. Particle stacks were then transferred to RELION^100^ for 2D and 3D classification. Following 2D classification, particles corresponding to SOSIP trimers, with or without visible Fab density, were subjected to 3D classification using the unliganded SOSIP trimer as the initial model. Classes exhibiting similar Fab-like densities were iteratively combined and reclassified until reliable immune complex reconstructions were obtained. Final particle subsets were refined by 3D auto-refinement in RELION using the unliganded SOSIP trimer as the starting model.

The resulting reconstructions were deposited in the Electron Microscopy Data Bank (EMDB) under the accession codes listed in the **Data Availability** section. Maps were visualized and segmented in UCSF Chimera^101^. For each immune complex, distinct Ab specificities targeting non-overlapping epitopes were segmented and combined with a docked Env reference model to generate the composite figures shown in the manuscript. Full particle stacks and 3D models used for Fab segmentation and figure generation are available upon request.

### cryoEM-based polyclonal epitope mapping (cryoEMPEM)

cryoEMPEM experiments were performed as described previously^26^. Polyclonal Fab/Fc samples were complexed with the corresponding SOSIP antigens at a 40:1 mass ratio to ensure comparable detectability of distinct Ab specificities across immune complexes. Assemblies typically contained 4-8 mg of Fab/Fc and 100–200 µg of SOSIP antigen and were incubated overnight (∼18 h) at room temperature to allow complex formation. For rabbit samples used to survey epitope targeting across antigens, the specific SOSIP-Ab combinations are listed in **Table S2**. Equivalent assembly conditions were used for BG505 SOSIP GT1.1 complexes with macaque samples (animals A12N115 and A12N074), as well as BG505 SOSIP IF with rabbit samples, except that Fab/Fc from NZWR-1 and NZWR-2 were pooled prior to analysis. Additionally, due to the fact that NZWR-1/2 induced disassembly of the BG505 SOSIP IF, the complexes were also produced following a shorter (2-hour) incubation of this antigen with the Fab. Therefore, two datasets were collected in the case of BG505 SOSIP IF, one with standard 18-hour assembly and one where assembly was allowed to proceed for 2 hours. In total, 13 polyclonal immune complexes were assembled across the training and validation datasets. Immune complexes were purified from unbound material by size-exclusion chromatography using a HiLoad 16/600 Superdex pg200 column (Cytiva) equilibrated in TBS. Fractions corresponding to immune complexes were concentrated using Amicon ultrafiltration devices (100 kDa cutoff) to final concentrations of 5-8 mg/mL and used directly for grid preparation.

CryoEM grids were prepared using the 300-mesh UltrAuFoil 1.2/1.3 or Quantifoil 1.2/1.3 grids (Quantifoil Micro Tools GmbH). Grids were glow-discharged for 30 s at 15 mA prior to sample application and prepared using a Vitrobot Mark IV (Thermo Fisher Scientific) at 100% humidity and 10 °C, with a blot force of 0, a wait time of 10 s, and blot times ranging from 3-7 s. Immediately before vitrification, lauryl maltose neopentyl glycol (LMNG) was added to each sample to a final concentration of 0.005 mM, and 3 µL was applied to the grid. Grids were plunge-frozen in liquid ethane cooled by liquid nitrogen and stored under liquid nitrogen until imaging. For each sample, 4-8 grids were prepared using varied blotting conditions. Grids were imaged at different facilities using either a Titan Krios or an Arctica microscope, and the relevant details are provided in **Extended data - Table 1**. Depending on the microscope, Leginon^102^ or EPU (Thermo Fisher) packages were used for automated data acquisition. Representative sample micrographs are shown **fig. S3**.

Data processing was done as previously described^26^ and illustrated in **fig. S4** Between 3’800 - 9’600 movie micrographs were collected per dataset (**Extended data - Table 1)**. Movie frames were aligned and dose-weighted using MotionCor2^103^, and the early processing steps were performed in the cryoSPARC package^104^. CTF parameters were estimated with CTFFIND4^105^. Immune-complex particles were picked using template-based picking and subjected to two rounds of 2D classification to remove contaminants and poorly aligned particles. In 12 out of 13 datasets, this procedure was applied to enrich for immune complexes containing SOSIP antigen in trimeric form. The exception was the BG505 SOSIP IF dataset collected following an 18-hour assembly. In this case, a significant concentration of immune complexes containing Fabs bound to gp120 monomers (formed from disassembled trimers) was detected and the 2D classification was used to select the monomeric particle subsets and reconstruct maps as illustrated in **fig. S35A**. There was only a small fraction of trimeric particles in this dataset, and they were not processed further.

Clean particle stacks were transferred to RELION for further processing. Following an additional round of 2D classification in RELION, particles corresponding to intact immune complexes were selected and refined in 3D using C3 symmetry with a soft mask around the antigen core. A low-pass-filtered BG505 SOSIP map was used as the initial model for all 3D steps to minimize model bias. Following the refinement of trimers, the particles were symmetry-expanded around the C3 axis to ensure that all epitope-paratope interfaces are overlaid onto every protomer. This step facilitates 3D classification, as the spherical maps can now be positioned against just a single epitope in a trimer. However, to avoid alignment of symmetry-related copies, particle alignment was constrained during subsequent classification and refinement. In the case of the monomer subset in the BG505 SOSIP IF dataset, all processing steps were done using C1 symmetry.

Epitope-focused sorting was performed using three rounds of focused 3D classification without image alignment (--skip_align, T = 16), each employing an 80 Å spherical mask positioned around each expected epitope (**fig. S4**). These classifications were carried out independently for each epitope cluster, with the number of classes adjusted based on Fab occupancy. Classes corresponding to structurally distinct Fab specificities were selected and refined separately using local angular searches only. The final round of 3D classification was performed using a full trimer-Fab complex mask, followed by 3D refinement and post-processing with the same solvent mask used during refinement. In some cases (e.g., BG505 SOSIP GT1.1 with A12N074 Fabs) excessive Fab densities from highly-occupied sites on trimer were subtracted from particles before the last round of refinement and postprocessing, to reduce the influence on image alignment. The resulting postprocessed maps were used for model building and deposition to EMDB. They are shown in **fig. S5-S13, S32, S35**. Detailed processing statistics are summarized in **Extended data - Table 2**, and the corresponding FSC curves are shown in **fig. S14-S18, S32, S35**.

### Model building into cryoEMPEM maps

Postprocessed maps generated in RELION were used for model building and refinement. As all antigens analyzed have previously reported structures deposited in the Protein Data Bank, these models were used as initial templates for the antigen components. Rigid-body docking was performed in UCSF Chimera^101^. A poly-alanine model of the immunoglobulin Fv region was docked into the Fab-corresponding density. Owing to structural differences between the heavy and light chains, including CDR2 and CDR3 lengths and the conformation of framework region 3, the Fv could be unambiguously oriented into the respective subdensities. The resulting antigen-Fab complex models were resampled against the corresponding density maps and imported into Coot^106^ for manual refinement. Antigen features were first adjusted to optimize agreement with the density, followed by manual construction of Ab CDR loops using poly-alanine backbones, as Ab sequence information is inherently unavailable. Identification of the CDR loop lengths and conformations was guided by the density constraints. Models were then subjected to relaxed refinement in Rosetta^107^, and model quality was assessed using Phenix Comprehensive Validation^108^ and EMRinger^109^ metrics. Manual and automated refinement cycles were repeated until optimal map fit and stereochemical metrics were achieved. The resulting model statistics are presented in **Extended data - Table 3**. Env antigens were numbered according to the HxB2 reference. Final models were used for downstream analyses and figure preparation and were deposited in the Protein Data Bank (PDB).

### Molecular surface generation and epitope definition

PDB models generated in the previous step were used to analyze epitope properties. Antigen proteins are preprocessed into triangulated mesh surfaces with MaSIF as previously described^50^. For each EM model, an optimal density threshold was determined by visual inspection, and only density points above this threshold were selected for further analysis. To automate the partitioning of the density map, density points within the antigen’s triangulated surface mesh, defined by the occupancy calculated with the Open3d Raycastingscene package^110^, were designated as antigen density. Density points within 1.5 Å distance to the glycan atoms are defined as glycan density. After excluding the antigen and glycan density, density points within 5 Å of the poly-alanine Ab model were defined as Ab density. Vertices on the antigen surface within 4 Å of the Ab density were defined as the epitope. Surface-exposed residues (i.e., RSA higher than 5%) with side-chain atoms within 3 Å to the epitope vertices were defined as epitope residues.

### Computation of surface features

Residues of the entire antigen are first grouped into a spatially connected cluster (*blob*) by identifying connected components in the adjacency graph using a distance threshold for the pairwise *Cα* Euclidean distances. Each blob is represented by its centroid, calculated as the mean of all coordinates, along with a covariance ellipsoid derived from the covariance matrix of its coordinates as previously described^111^. The ellipsoid’s orientation is determined by eigenvectors, while its axis lengths are scaled eigenvalues. The ellipsoid surface was then parameterized on a regular spherical grid (*θ*, *ϕ*) with *θ* ∈ |0,2*π*| and *ϕ* ∈ |0,*π*|. Each direction vector (*cos cos θ sin sin ϕ, sin sin θ sin sin ϕ, cos cos ϕ*) was scaled by the ellipsoid radii (*r_x_, r_y_, r_z_*), rotated by the covariance eigenvectors, and translated by the centroid *μ*. To quantify surface accessibility for each surface vertex *s*, a normalized protrusion index *PI* was computed as

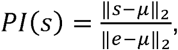

where *e* is the nearest point on the fitted ellipsoid surface. The PI is therefore normalized to values between 0 and 1, where 0 indicates proximity to the centroid, while 1 corresponds to areas that are at the furthest-most edge of the molecule.

Principal curvatures for each vertex are computed using the libigl library with the default setting^112^. Shape index *SI* is calculated based on the principal curvature values as previously described using the following formula:

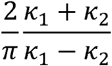

The SI is therefore normalized to values ranging from -1 to +1, with negative values corresponding to concave features and positive values corresponding to convex features.

Chemical features, including the hydropathy index, Poisson-Boltzmann continuum electrostatics, and the presence of free electrons and proton donors, are directly obtained from MaSIF preprocessing outputs.

### Model setup and MD simulations

CryoEM structures were completed with the missing residues modeled using MODELLER^113^. Full-length glycosylation was added following our previously established in-house method^114^. Atomistic molecular dynamics simulations for each structure were performed in AMBER software^115^ with CHARMM36^116^ and CHARMM36m^117^ forcefields. The structures were solvated in an explicit solvent water box with 25 angstrom padding, and a TIP3P^118^ water model modified for SHAKE^119^. The systems were neutralized with an ionic concentration of 150 mM KCl. Steepest-descent energy minimizations were conducted on the systems where constraints on restricted atoms were gradually removed as the systems approached an energy minimum. The first equilibration step involved constant volume and temperature conditions (NVT) for 2 ns. The second and third equilibration steps involved constant pressure and temperature conditions (NPT) for 10 ns each for a total of 20 ns. Position restraints on all non-hydrogen atoms were in place for the NVT equilibration, and these were relaxed to position restraints on the protein backbone alpha carbons (C𝛼) during the NPT equilibrations. During the production run, each of the systems was run for 1.2 μs, and the initial 200 ns was excluded from each trajectory to reach RMSD equilibrium and to adjust for initial instabilities for subsequent analyses. During the simulations, a Monte Carlo barostat was used to maintain an isotopic pressure of 1.01325 bar with compressibility of 4.5×10^-5^ bar^-1^, and a temperature of 310 K was maintained with a Langevin thermostat. The Particle Mesh Ewald method was used for determining energies^120^. AMBER recommended Hydrogen Mass Repartitioning^121^ was done, which allowed for a timestep of 4 fs during the simulation process, and the simulations were run in NVIDIA Tesla H100 GPU machines to allow for increased simulation bandwidth. After the simulations were completed, an RMS fit was applied to the C𝛼 _atoms of each residue in the gp41 region except for the disordered C-terminal tail.

### Root mean square fluctuations (RMSF)

RMSF provides a measure of the flexibility of individual residues within the protein structure, and it can be calculated through the following formula:

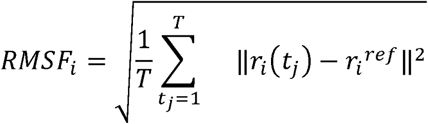

where T is the total time in the trajectory, t is the index time used for a calculation at a particular point, 𝑟_𝑖_ are the atomic coordinates in the i^th^ frame, and 𝑟_𝑖_ is the reference structure. These calculations were performed in MDTraj using the mean structure as the reference for the calculations, with the first 200 ns excluded to omit unstable protein dynamics. Additionally, the three protomer units for each structure were averaged together to yield an RMSF plot of the mean fluctuations across the three protomers.

### Glycan Encounter Factor (GEF)

Glycan shielding effect over the protein surface was calculated per-residue, based on the glycan encounter factor (GEF) score previously established by the authors^59^. This was calculated as the geometric mean of the probability that a probe approaching a surface residue would encounter glycan heavy atoms, in perpendicular and tangential directions. A probe size of 6 angstroms was chosen to mimic a typical hairpin loop. Calculations were performed in x-y-z cardinal directions, and their geometric mean was taken and normalized. Computational modeling calculations were implemented with VMD 1.9.3^122^ and Python.

### Machine learning on protein surface features to predict ASI score

For each surface vertex *i*, an amino-acid fingerprint vector *f_i_* ∈ *R*^21^ was constructed to encode the local chemical environment contributed by nearby antigen residues. The fingerprint consists of 20 standard amino-acid channels, plus an additional channel that represents proximity to PNGS by identifying the N-linked glycan sequon (N-X-S/T).

For a given vertex *i*, the *n* = 8 nearest antigen residues were identified, indexed *j* = 1, …, 8, with corresponding distances *d_ij_*. Contributions were accumulated for residues within a distance cutoff *d_cut_* = 7.0 Å. The contribution of residue *j* to amino-acid type *k* was weighted by its relative solvent accessibility *RSA_j_* and inversely by distance:

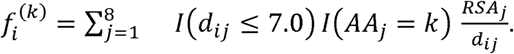

An additional PNGS channel was defined using a broader distance cutoff *d_PNGS_* = 15 Å:

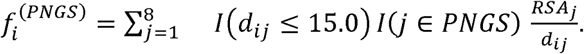

The resulting vector *f_i_* = (*f_i_*^(1)^, …, *f_i_*^(20)^, *f_i_*^(*PNGS*)^) defines the amino-acid composition feature representation for vertex *i*.

To train a model that takes surface features of each vertex as input and predicts its immunogenicity, we first create a descriptor by concatenating all pre-computed features. *PI*, *SI*, and the Euclidean distance to the nearest PNGS site were used as the geometric features, which were concatenated with the amino-acid composition vector, resulting in a 24-dimensional feature descriptor that serves as the model’s input.

In each of the 100 pAbC models, positive labels are assigned to the vertices in direct contact with the Ab density (as defined in **Molecular surface generation and epitope identification**). Only the vertices that are distant from epitope residues from any of the models from the same antigen group are labeled with negative labels. For training purposes, negative-label sites were randomly selected to match the number of positive-label sites in each cryoEM model.

Using the 24-dimensional feature descriptors as the input, an XGBoost classifier model (n_estimators=500, learning_rate=0.1) was trained to predict the labels for each vertex, and the probability of the positive label was used as the ASI score. Ablation of geometric and amino acid composition features was also performed, yielding descriptors with 21 and 3 dimensions, respectively. To understand the global impact of each feature in the descriptors on the final prediction outcome, their Shapley values were calculated using the SHAP Python package^64^, providing a global interpretation of their positive or negative impact on the ASI score.

### Modeling ASI score to AlphaFold2 models and experimental structures

AlphaFold2 models of HIV Env antigens (**fig. S1**) were generated using ColabFold (https://github.com/sokrypton/ColabFold.git) and formatted into HxB2 residue IDs. The molecular surface of each antigen structure, derived from AlphaFold2, PDB, or EMPEM models, was generated. The ASI score was then computed for each surface vertex as described above. For AlphaFold2-generated models, structural reliability was incorporated by multiplying the ASI value of each vertex by the pLDDT score of the nearest residue. For HIV Env antigens derived from EMPEM models, the corresponding AlphaFold2-generated antigen served as a reference, and the pLDDT value for each residue was mapped onto the equivalent residue in the EMPEM structure. In this way, ASI scores across different conformational epitopes were calibrated according to the local confidence of the underlying structural model. Because pLDDT values range from 0 to 100, this procedure downweights regions with greater conformational uncertainty. For experimental structures determined by electron microscopy, model-to-map metrics (e.g., cross-correlation or Q scores) can be applied per residue and serve as analogous measures, provided that the atomic model has been built accurately

### Design of BG505 SOSIP IF

To engineer amino acid composition within the CD4bs and FP epitopes, we used the AnalyzeAlign tool from the LANL HIV database^67^ to generate aligned FASTA files for defined Env segments spanning the FP (residues 512–523) and CD4bs (residues 198-199, 428-431, and 455-461). Searches were performed using the Filtered Web database, comprising ∼7,600 unique HIV Env sequences. Aligned sequence segments were downloaded and analyzed using an in-house script that treated each FASTA file as text input and quantified the frequency of selected amino acids (W, Y, F, L, I, D, E, R, K) within each segment. For each fragment, the top 100 scoring sequences featuring the highest level in the selected amino acids were further ranked based on tryptophan (W) content, as meta-analysis indicated strong enrichment of this residue in immunodominant epitopes. Final sequences were selected for all four fragments (one FP and three CD4bs segments) and introduced into the BG505 SOSIP MD39 background. The resulting construct additionally incorporated N-linked glycan modifications in the base, N611 (GH), N241/N289 (GH), and C3/V5 epitopes, which have previously been shown to reduce targeting of these sites^12,123,124^. A complete list of all mutations and their rationale is provided in **Table S3**. The CD4bs and FP epitope revertants used in the ELISA experiments were generated by reverting the mutations in the corresponding epitope back to native BG505. Mutagenesis, sequence verification, protein expression, and purification of BG505 SOSIP IF and the epitope-reverted constructs were performed as described above.

### Negative-stain EM of the designed BG505 SOSIP IF

BG505 SOSIP IF was diluted to 20 µg/mL and applied to glow-discharged (15 mA, 30 s) carbon-coated Cu grids (CF300-Cu-50, Electron Microscopy Sciences) as described previously^26^. Grids were negatively stained using 2% uranyl formate with a double-staining protocol (15 s followed by 60 s). Data were collected on a Thermo Fisher Talos L120C G2 microscope at 120 keV, 57’000 x magnification (2.44 Å/pixel), with an electron dose of 20 e^-^/Å^2^ and defocus range -1.0 to -4.0 µm in 0.5 µm steps. Images were recorded with a Ceta S 16M CMOS camera using EPU automated collection software (Thermo Fisher). Particle picking and 2D classification were performed in CryoSPARC^104^ using the blob picker and template-based workflows. The quantitative analysis was performed as described before^125^.

### Biolayer interferometry

The experiments were performed as described previously^16^. Kinetics buffer (DPBS supplemented with 0.1% BSA and 0.02% Tween-20) was used to prepare Ab and antigen solutions. IgG Abs were diluted to 5 µg/mL, while BG505 SOSIP IF and BG505 SOSIP MD39 antigens were adjusted to 500 nM. Binding measurements were collected using an Octet Red96 instrument (ForteBio). Abs were captured on anti-human IgG Fc (AHC) biosensors. Association and dissociation steps were both set to 300 s. Raw data were processed with Octet Data Analysis v9.0, including background subtraction using buffer-only controls, y-axis alignment to the baseline preceding association, and interstep correction between association and dissociation. Data were exported and plotted in Excel. Experiments were performed in duplicate to assess reproducibility; a single representative measurement is shown.

### Glycan composition analysis using LC-MS

BG505 SOSIP IF was prepared for mass spectrometry by denaturation (50 mM Tris/HCl, pH 8.0, 6 M urea, 5 mM DTT), reduction and alkylation with iodoacetamide, and buffer exchange into 50 mM Tris/HCl (pH 8.0). The sample was digested overnight with trypsin, chymotrypsin, or α-lytic protease (1:16 w/w). Resulting peptides were dried, extracted, and re-suspended in 0.1% formic acid prior to nanoLC-ESI-MS on an Orbitrap Eclipse HPLC-MS system with stepped HCD fragmentation. Glycopeptide fragmentation spectra were processed with Byos (Protein Metrics Inc.) and manually validated for correct b/y ions and diagnostic oxonium ions. Data were searched against a Glycan/peptide library to determine glycoform composition and site occupancy, and the relative abundance of each glycoform was calculated from extracted chromatographic areas. This workflow categorizes glycans as oligomannose, hybrid, or complex types based on composition and is consistent with previously described site-specific glycosylation analysis strategies used for HIV-1 Env immunogen evaluation^126^.

### Immunization experiments with BG505 SOSIP IF

The rabbit immunization study was conducted in accordance with protocols approved by the Institutional Animal Care and Use Committee (IACUC) at The Scripps Research Institute, San Diego, United States (protocol no. 21-0012-2). Rabbits were housed, immunized, and bled in compliance with the U.S. Animal Welfare Act and other applicable federal statutes and regulations, and in accordance with the *Guide for the Care and Use of Laboratory Animals* of the National Research Council. The study was additionally reviewed and approved by the Animal Research Ethics Committee (AREC) of the Swiss Federal Institute of Technology Lausanne (EPFL).

Immunologically naïve New Zealand White rabbits were housed at the Animal Models Core Facility at The Scripps Research Institute. Two animals were immunized intramuscularly in the hind-leg quadriceps with 30 µg of BG505 SOSIP IF formulated with 225 µg of SMNP adjuvant^76,77^. Each rabbit received three doses at weeks 0, 4, and 12. Blood was collected at weeks 0 and 6 (20 mL each), and a terminal bleed (50 mL) was performed at week 14. Plasma was separated immediately and stored at -80 °C until use.

### Enzyme-linked immunosorbent assays

BG505 SOSIP IF, CD4bs knock-out, and FP knock-out trimers were diluted to 3 µg/mL in 0.1 M NaHCO₃ and coated onto clear flat-bottom 96-well immuno plates (Thermo Fisher Scientific) at 50 µL per well for 2 h at room temperature. Plates were blocked overnight at 4 °C with 300 µL per well of PBS containing 10% BSA and 0.05% Tween-20. Unbound antigen was removed by three washes with TBS supplemented with 0.1% Tween-20. Purified IgG from rabbit sera NZWR-1 and NZWR-2 was added in three-fold serial dilutions starting at 150 µg/mL (30 µL per well) and incubated for 2 h at room temperature. Plates were washed three times with TBS + 0.1% Tween-20, followed by incubation with goat anti-rabbit IgG secondary Ab (Jackson ImmunoResearch, 111-055-003) diluted 1:5,000 in PBS containing 1% BSA for 1 h at room temperature. After three final washes, colorimetric detection was performed using 1-Step PNPP substrate (Thermo Scientific), and absorbance at 405 nm was measured on a Tecan Infinite microplate reader. All trimers and control antigens were coated on the same plate. To ensure that differences in EC₅₀ values were not due to antigen quality, ELISAs were also performed with the monoclonal Ab 2G12, which showed comparable binding to all three constructs and to the dual-epitope knock-out probes (EC₅₀ values within 18% of each other).

## Data availability

Three-dimensional maps and atomic models from the EM analysis have been deposited in the Electron Microscopy Data Bank (http://www.emdatabank.org/) and the Protein Data Bank (http://www.rcsb.org/), respectively. Accession numbers are provided in Extended Data Tables 2 and 3. Negative-stain EM maps have been deposited in EMDB under accession ID EMD-56907, EMD-56914, EMD-56922, EMD-56923, EMD-56924, EMD-56925, EMD-56927, EMD-56943, EMD-56942, EMD-56941, EMD-56940, EMD-56935, EMD-56927. The ASI prediction software will be deposited to https://github.com/xiaosh9527/antigenic_surface_immunodominance.git. All entries will be released upon publication of the manuscript. Additional raw and processed data will be made available upon reasonable request.

## Acknowledgments

We would like to express sincere gratitude to all the funding bodies for making this research possible. AA was supported by the American Foundation for AIDS Research (amfAR) Mathilde Krim Fellowship in Basic Biomedical Research [grant number 110413-73-RKVA]. AA and ABW are also funded by the NIH grant AI136621. Glycan analysis was supported by the Gates foundation CAVD (INV-070116). This project also received funding from the National Institute of Allergy and Infectious Diseases of the NIH award number P01 AI110657 (to A.B.W., J.P.M., and R.W.S.) and the Gates Foundation through grants INV-063951 and INV091003 (to R.W.S. and J.P.M.).

We thank Florence Pojer, Kelvin Lau, and Yoan Duhoo from the EPFL Protein Production and Structure Core Facility for advice, monoclonal Ab production, and cryo-EM grid preparation. We are grateful to the teams at the Interdisciplinary Centre for Electron Microscopy (CIME) and the Dubochet Center for Imaging (DCI) in Lausanne for assistance with cryo-EM data collection and processing. We further acknowledge Hannah L. Turner at the Scripps Research Institute for her help with electron microscopy experiments performed at that facility. A portion of this research was supported by NIH grant R24GM154185 and performed at the Pacific Northwest Center for Cryo-EM (PNCC) with assistance from Sean Mulligan. We also thank the IT departments of the School of Life Sciences at EPFL (Philippe Borel) and the Scripps Research Institute (Jean-Christophe Ducom) for computational support. Finally, we express our deepest gratitude and dedicate this work to Bill Anderson for his many years of training and steadfast support to users at the electron microscopy center at The Scripps Research Institute.

## Author Contributions

JS, SX, AA, ABW, RWS, BEC, JPM, MC, and SC conceptualized the study, designed experiments, and analyzed the data. JS, SX, AA, ERE, SB, AZ, DMC, GO, WZ, TGC, planned and performed the experiments. JS, SX, and AA assembled the figures and wrote the initial draft of the manuscript. ABW and BEC provided extensive feedback during the early stages of manuscript assembly. All authors provided input and contributed to the final version of the manuscript.

## Declaration of Interests

All authors declare no competing interests.

## Notes

### Competing Interest Statement

The authors have declared no competing interest.

