## Supplementary Information for "Decoding epitope immunodominance in HIV Env using cryoEM and machine learning"

### SUPPLEMENTARY MATERIAL

**Table S1:** Epitope definitions for HIV Env

| **Epitope** | **Color** | **Description** |
| --- | --- | --- |
| **Base** |  | Epitopes at the bottom of the trimer, comprising C1 (N-term) and HR2 (C-term) regions. In the context of virions these areas normally face the viral membrane. |
| **N611 (GH)** |  | Epitopes surrounding residue N611 in HR2 with minor contribution from HR1. N611 glycosylation is occasionally incomplete leading to the formation of a glycan hole (GH) in a subset of molecules. |
| **N625 (GH)** |  | Epitopes surrounding residue N625 in HR2. N625 glycosylation is occasionally incomplete leading to the formation of a glycan hole (GH) in a subset of molecules. |
| **N241/N289 (GH)** |  | Epitopes proximal to the N241 and N289 residues. These glycosylation sites are absent in some strains leading to the formation of a glycan hole (GH). |
| **V1/V2** |  | Epitopes at the trimer apex comprising elements of V2 and V1 loops with minor contribution of V3 loop elements in certain cases. V2 glycans (e.g., N160, N185) shield these epitopes. |
| **V1/V3** |  | Epitopes at the trimer apex comprising elements of V1 and V3 loops with minor contribution of V2 loop elements in certain cases. V1 glycans (e.g., N156, N133, N137) shield these epitopes. |
| **C3/V5** |  | Epitopes at the edge of the gp120 blade distal from the C3 axis comprising the C3 and V5 elements flanked by N-linked glycans at positions N355 and N465. |
| **gp120-gp120 interface** |  | Epitopes comprising elements of the V3-tip (or crown), C1 (resi: 61-71) and C2 (resi: 207-209), flanked by the N197 and N262 N-linked glycan sites. |
| **Silent Face (SF)** |  | Epitopes formed primarily by conserved C2, C3 and C4 loop elements and covered by several N-linked glycans (e.g., N295, N332, N411, N448) |
| **CD4 binding site (CD4bs)** |  | Epitopes surrounding the receptor binding site, formed primarily by the conserved C1, C2, C3, C4 and C5 loop elements and flanked by the N-linked glycans at N276, N197 and N386 |
| **Fusion peptide (FP)** |  | Epitopes incorporating the fusion peptide (resi: 512-523) and the surrounding C1, C2, HR1 and HR2 elements. The site partially overlaps with the N611 and N241/N289, but includes FP contacts. |

**Table S2:** Information regarding the Env and polyclonal samples subjected to cryoEMPEM characterization^16,17,37,38^.

| **Construct** | **Immunization** | **Animal model** | **Time point** | **Animal ID** |
| --- | --- | --- | --- | --- |
| B41 SOSIP.v4.1 | C0045-15 | Rabbit | Week 22 | r1645, r1646 |
| AMC009 SOSIP.v4.2 | C0048-15 | Rabbit | Week 22 | UA0062, UA0065 |
| 16055 SOSIP.v8.3 | C0038-19 | Rabbit | Week 22 | r2463, r2464 |
| CH505(B) SOSIP.v8.1 | C0038-19 | Rabbit | Week 22 | r2474 |
| ConM SOSIP.v9 | C0171-017 | Rabbit | Week 22 | r2381, r2382 |

**Table S3:** Mutations introduced into the BG505 SOSIP IF construct (locations are based on HxB2 numbering).

| **Background** | **Rationale** |
| --- | --- |
| BG505 SOSIP MD39 | Stabilized BG505 construct with well-characterized immunogenic properties. For details see Steichen et al., 2016^127^. |
| **Mutations** | **Rationale** |
| T198L, S199T, Q428W,  R429E, I430V, G471W, | Mutations within the CD4bs intended to increase immunodominance of this epitope. Engineered for the purpose of this study. |
| V513I, V518W | Mutations within the FP intended to increase immunodominance of this epitope. Engineered for the purpose of this study. |
| S519F | Reversal of a stabilizing mutation in the FP region of MD39 to the original BG505 sequence^127^. |
| C605T, A662C | Switching from the SOS mutation in SOSIP to an alternative mutation to crosslink gp41 to gp120 which was found to improve trimer stability^128^. |
| P240T, S241N | Introduction of a PNGS at position N241 to cover a glycan hole that exists in BG505 but is absent in most HIV Env sequences^12,124^. |
| T290E, P291S | Introduction of a PNGS at position N289 to cover a glycan hole that exists in BG505 but is absent in most HIV Env sequences^12,124^. |
| T465N | Introduction of an additional native PNGS at position N465 to shield an immunodominant epitope in the C3/V5 region of BG505^12,124^. |
| R500N, K502T | Introduction of an artificial PNGS site in the Base epitope to shield this highly immunodominant epitope. Engineered for the purpose of this study. |
| S613T | Introduction of T into the N611 PNGS sequon to improve the processing and increase the occupancy of glycan N611^124^. |

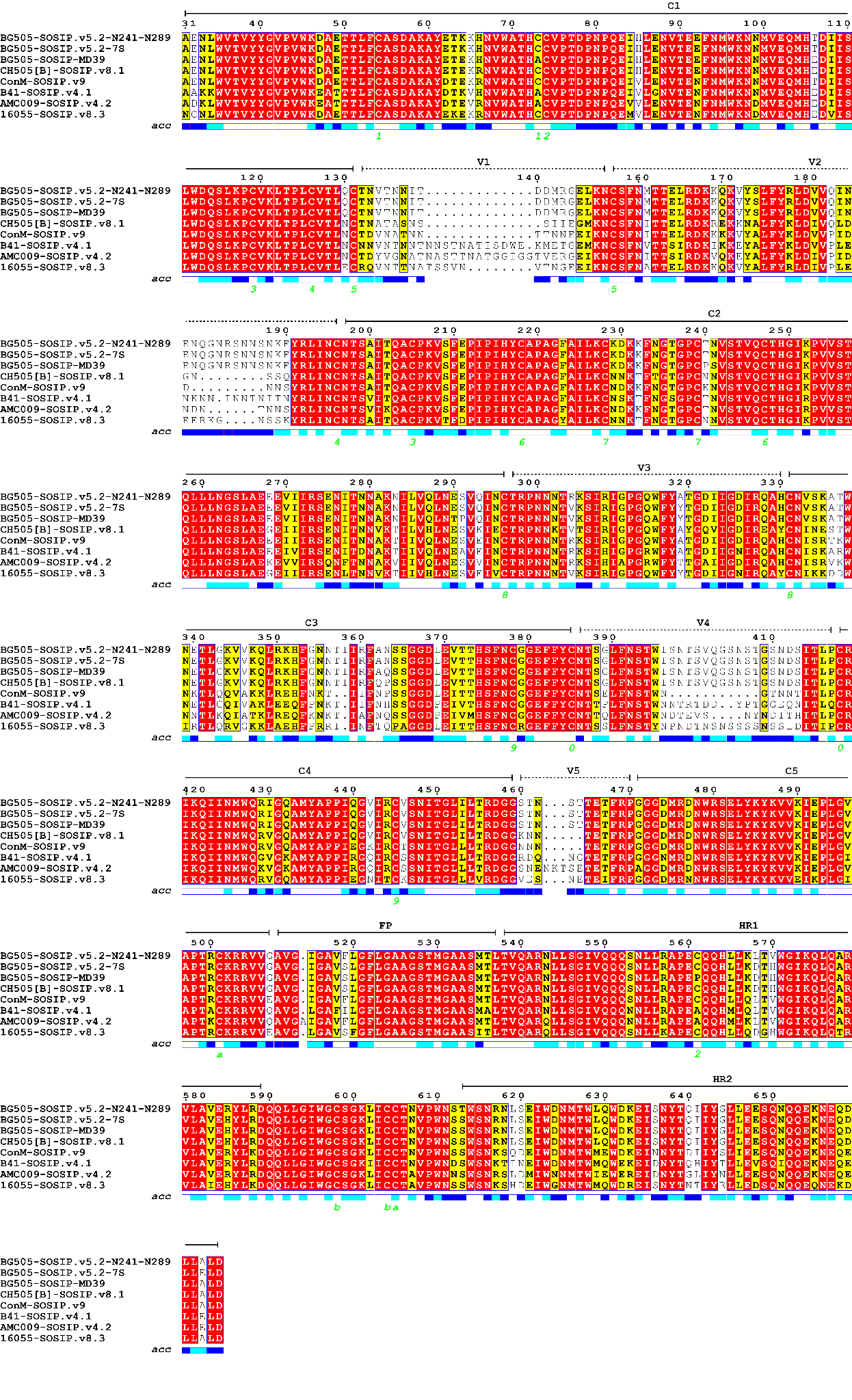

**Fig S1.** Sequence alignment of HIV Env immunogens used in this study.

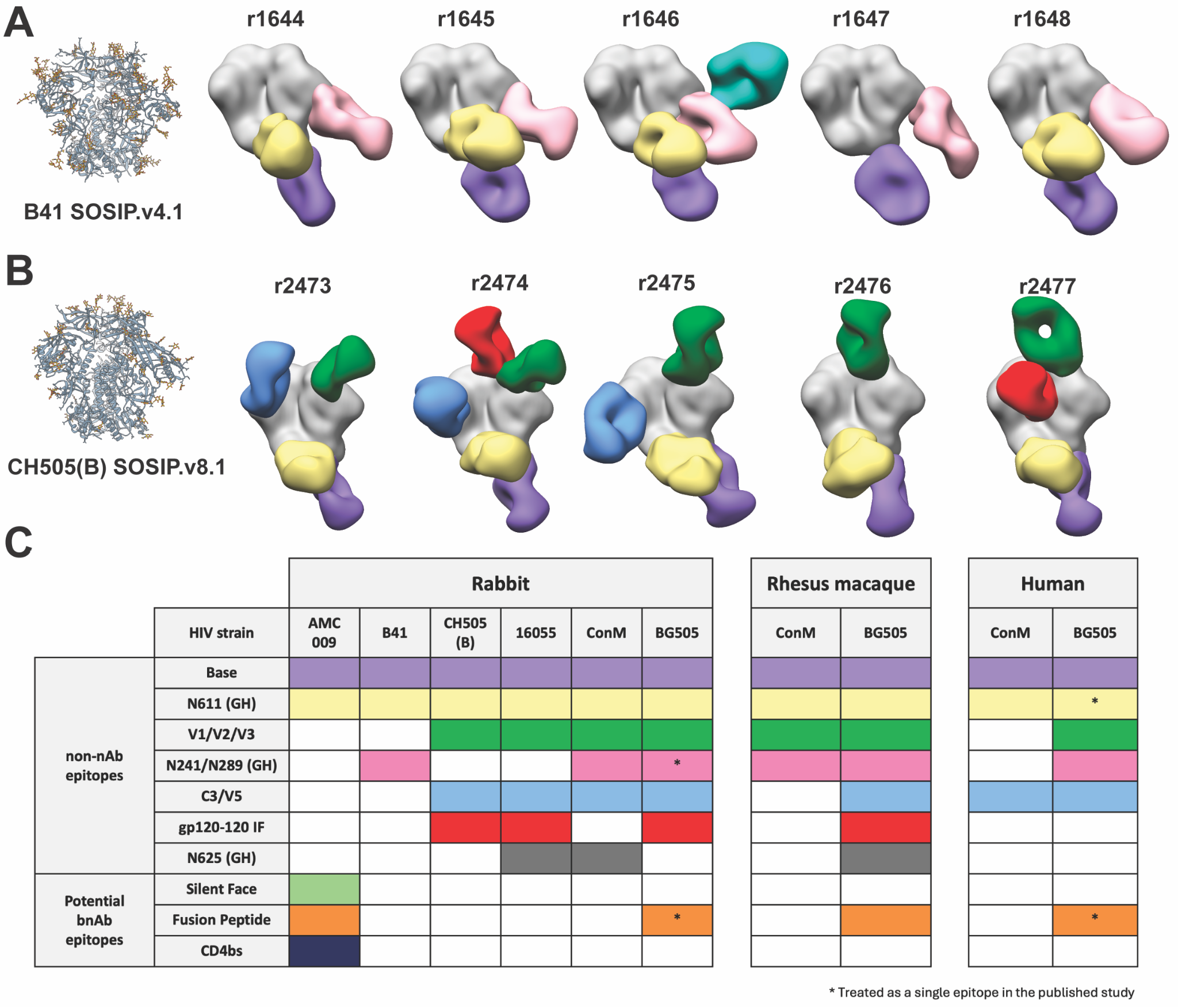

**Fig S2. Negative-stain EMPEM results.** Composite figures showing (**A**) B41 SOSIP.v4.1 and (**B**) CH505(B) SOSIP.v8.1 in complex with polyclonal antibodies isolated from immunized rabbits. The identification number of the animal from which the antibodies were derived is indicated above each panel. Antigens are shown in gray, and antibodies are colored according to their epitope specificities (see panel C for the color scheme). In addition to Env-directed antibodies we also detected an anti-immune-complex antibody in animal r1646, shown in aquamarine green/blue color. This antibody was already discussed elsewhere^30^. (**C**) Summary of epitope-targeting analyses performed by negative-stain EMPEM across different antigens and animal models. Colored entries indicate successful detection of antibodies targeting the corresponding epitope on the given antigen. The first three columns (AMC009, B41, and CH505(B)) present new data reported in this study, whereas data shown in the remaining columns were adapted from previously published studies^16,26,30,40,42–46^.

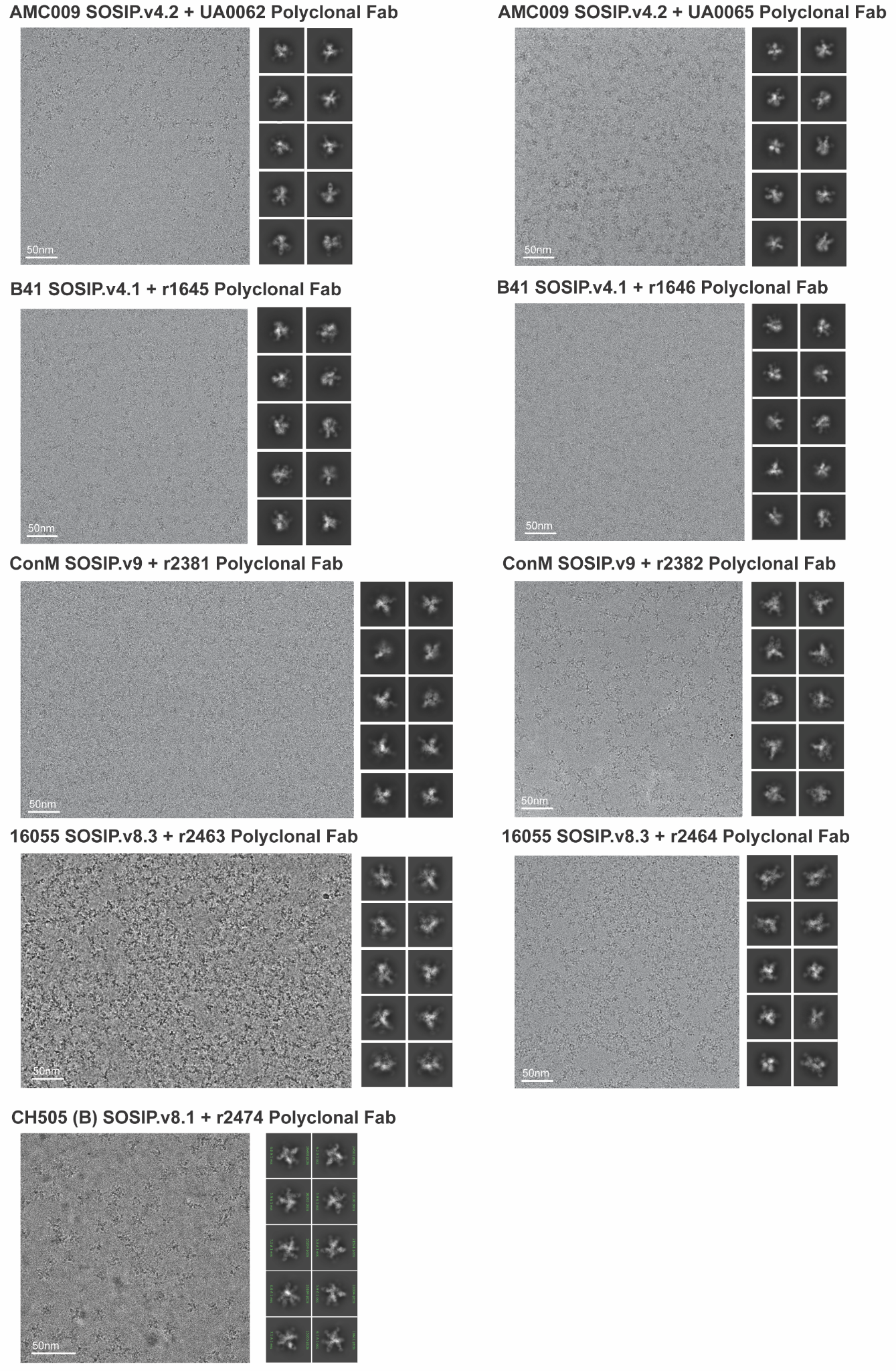

**Fig S3. Examples of raw cryoEMPEM data.** Representative raw micrograph and 2D class averages for the cryoEMPEM datasets collected in this study.

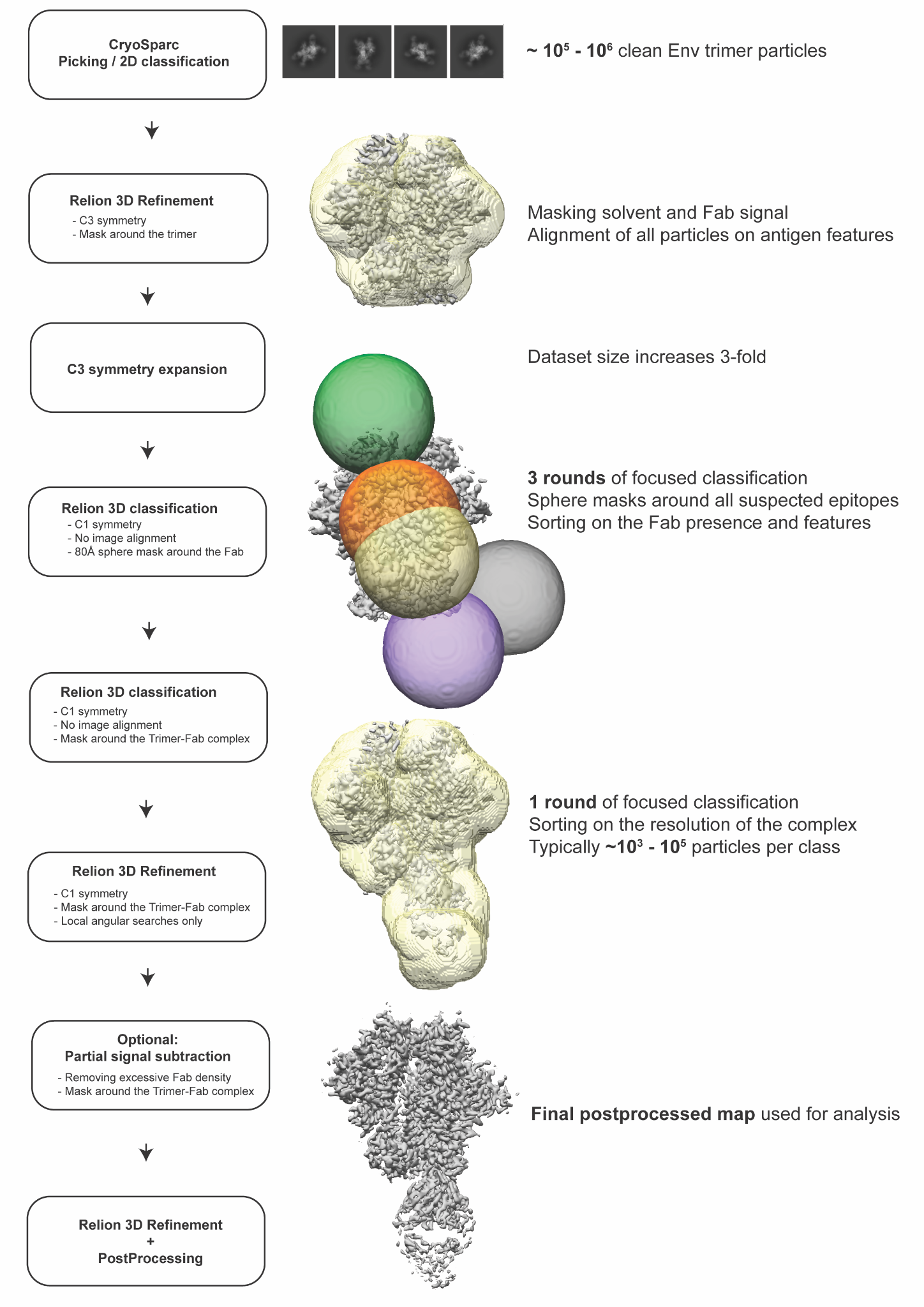

**Fig S4. Workflow employed to process cryoEMPEM data and generate 3D maps.**

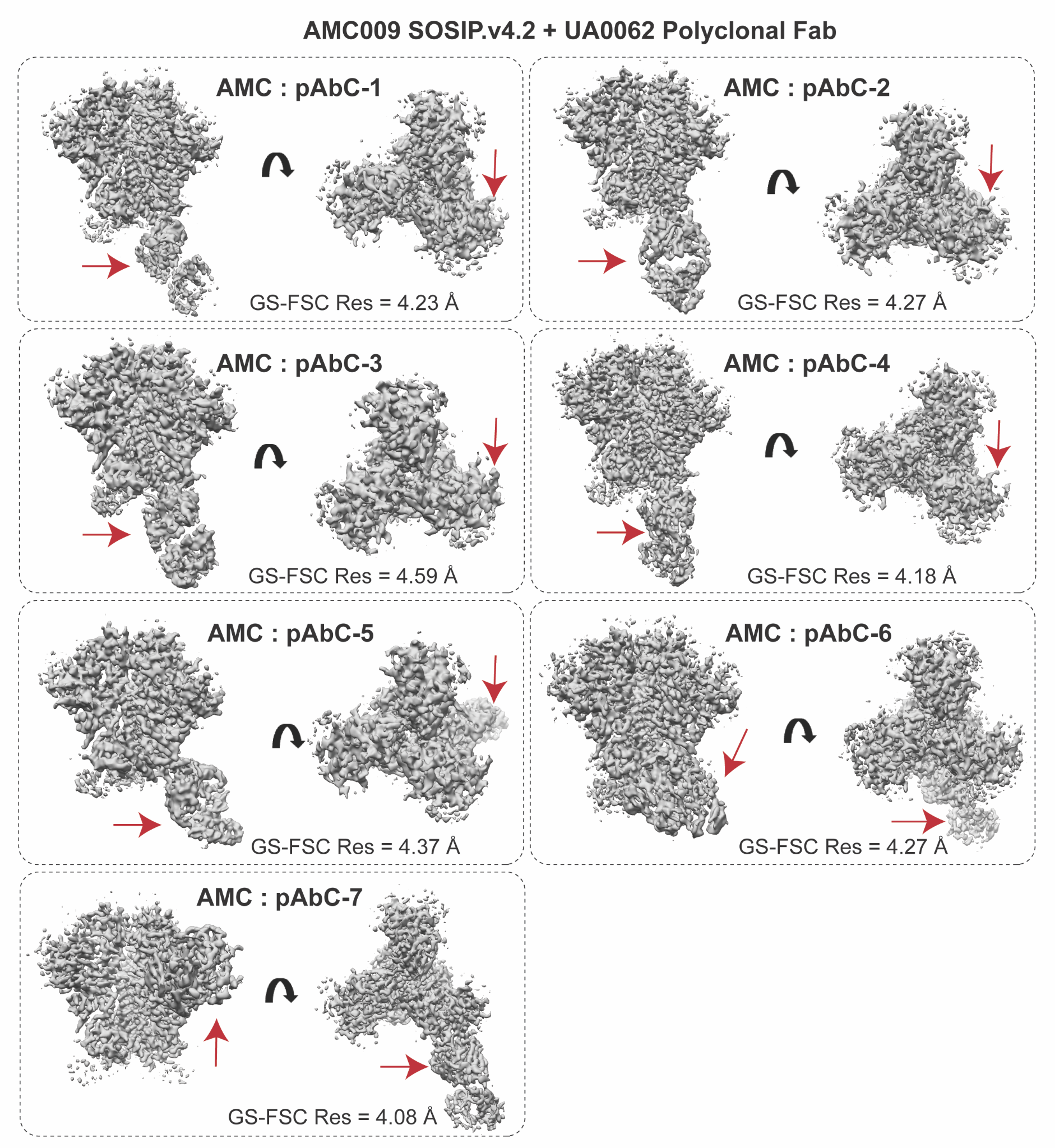

**Fig S5.** EM density maps recovered by cryoEMPEM of AMC009 SOSIP.v4.2 in complex with UA0062 polyclonal antibodies (as Fab). Red arrows point towards the Fab-corresponding density.

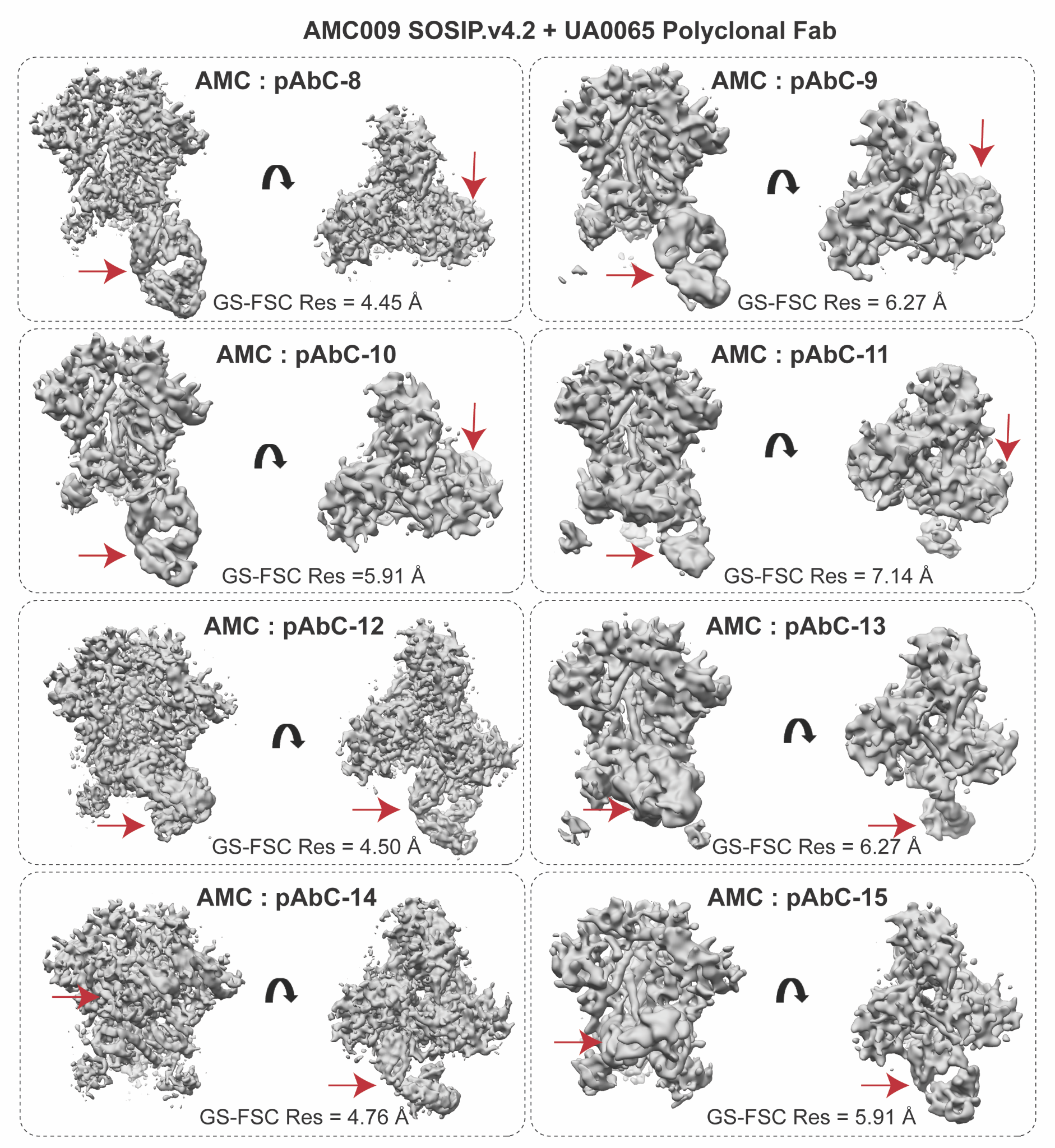

**Fig S6.** EM density maps recovered by cryoEMPEM of AMC009 SOSIP.v4.2 in complex with UA0065 polyclonal antibodies (as Fab)**.** Red arrows point towards the Fab-corresponding density.

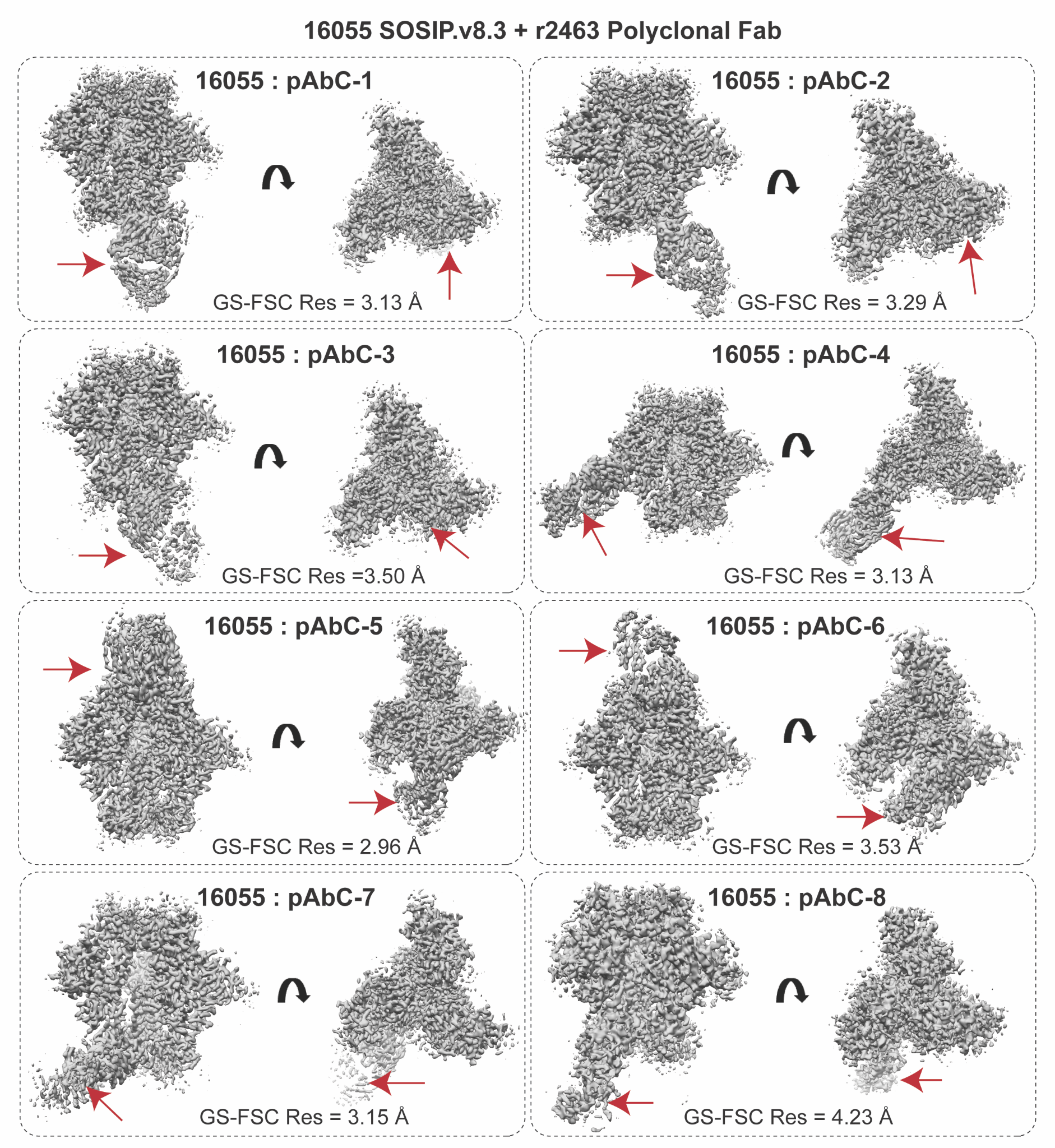

**Fig S7.** EM density maps recovered by cryoEMPEM of 16055 SOSIP.v8.3 in complex with r2463 polyclonal antibodies (as Fab)**.** Red arrows point towards the Fab-corresponding density.

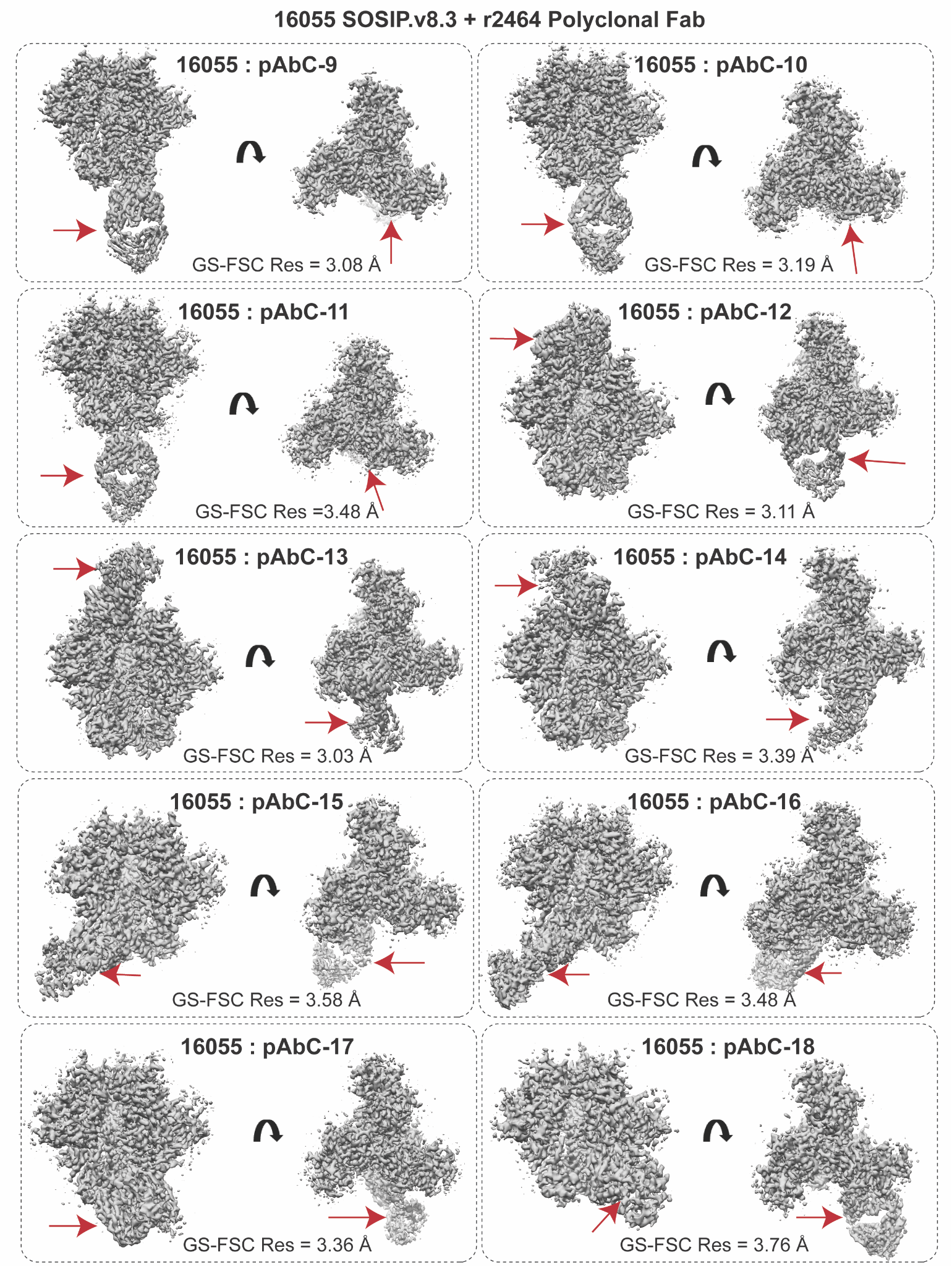

**Fig S8.** EM density maps recovered by cryoEMPEM of 16055 SOSIP.v8.3 in complex with r2464 polyclonal antibodies (as Fab)**.** Red arrows point towards the Fab-corresponding density.

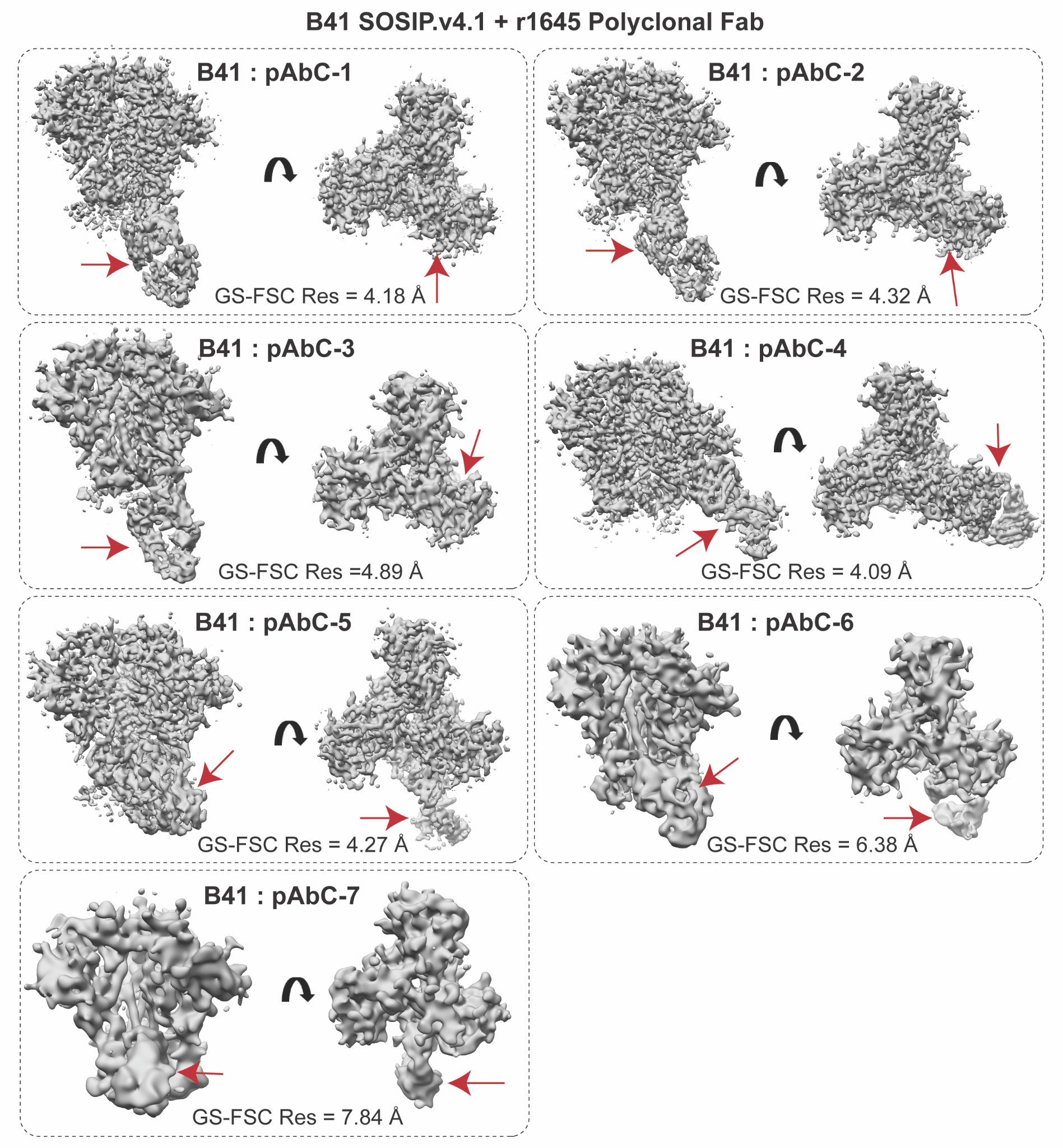

**Fig S9.** EM density maps recovered by cryoEMPEM of B41 SOSIP.v4.1 in complex with r1645 polyclonal antibodies (as Fab)**.** Red arrows point towards the Fab-corresponding density.

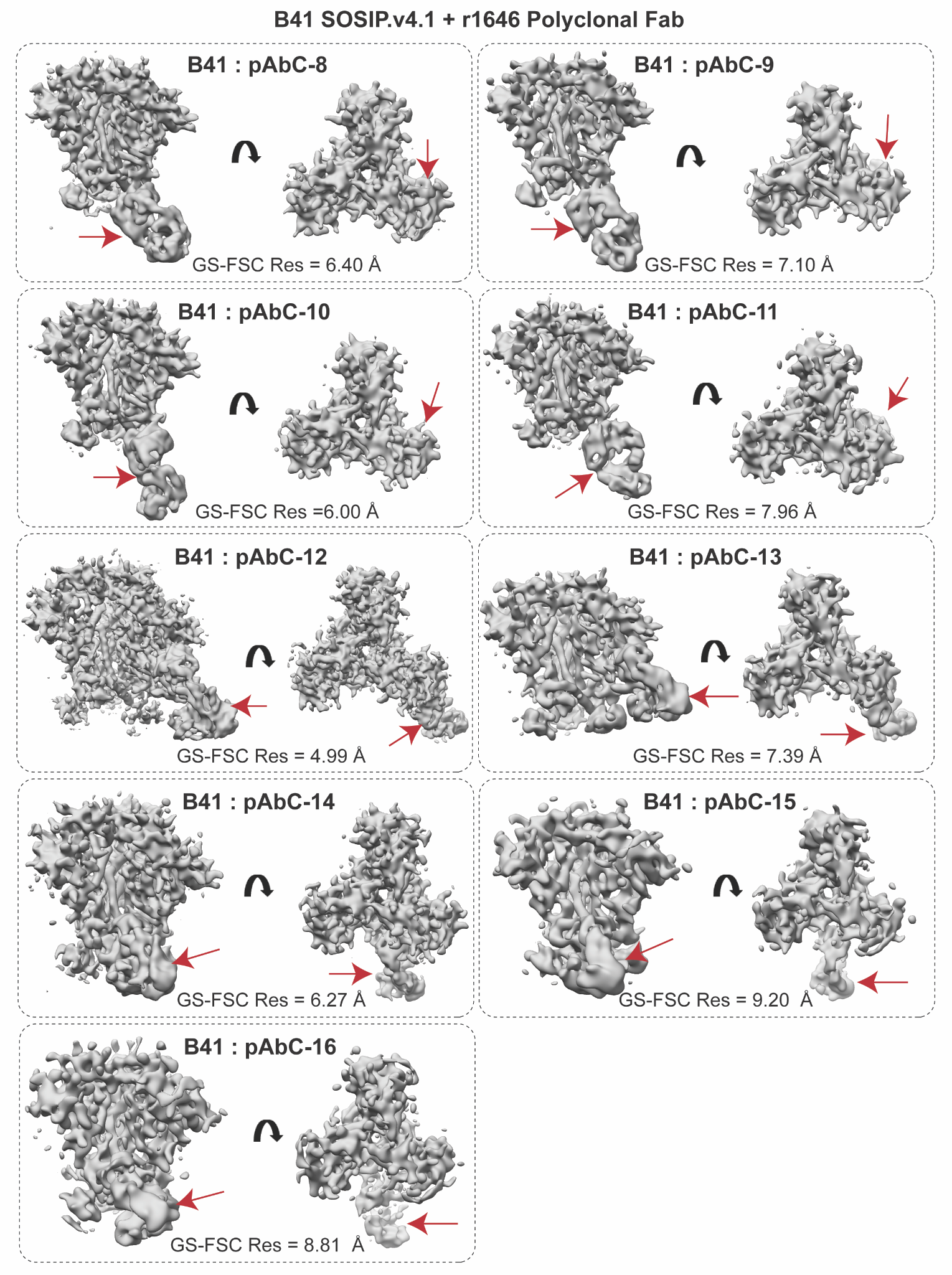

**Fig S10.** EM density maps recovered by cryoEMPEM of B41 SOSIP.v4.1 in complex with r1646 polyclonal antibodies (as Fab)**.** Red arrows point towards the Fab-corresponding density.

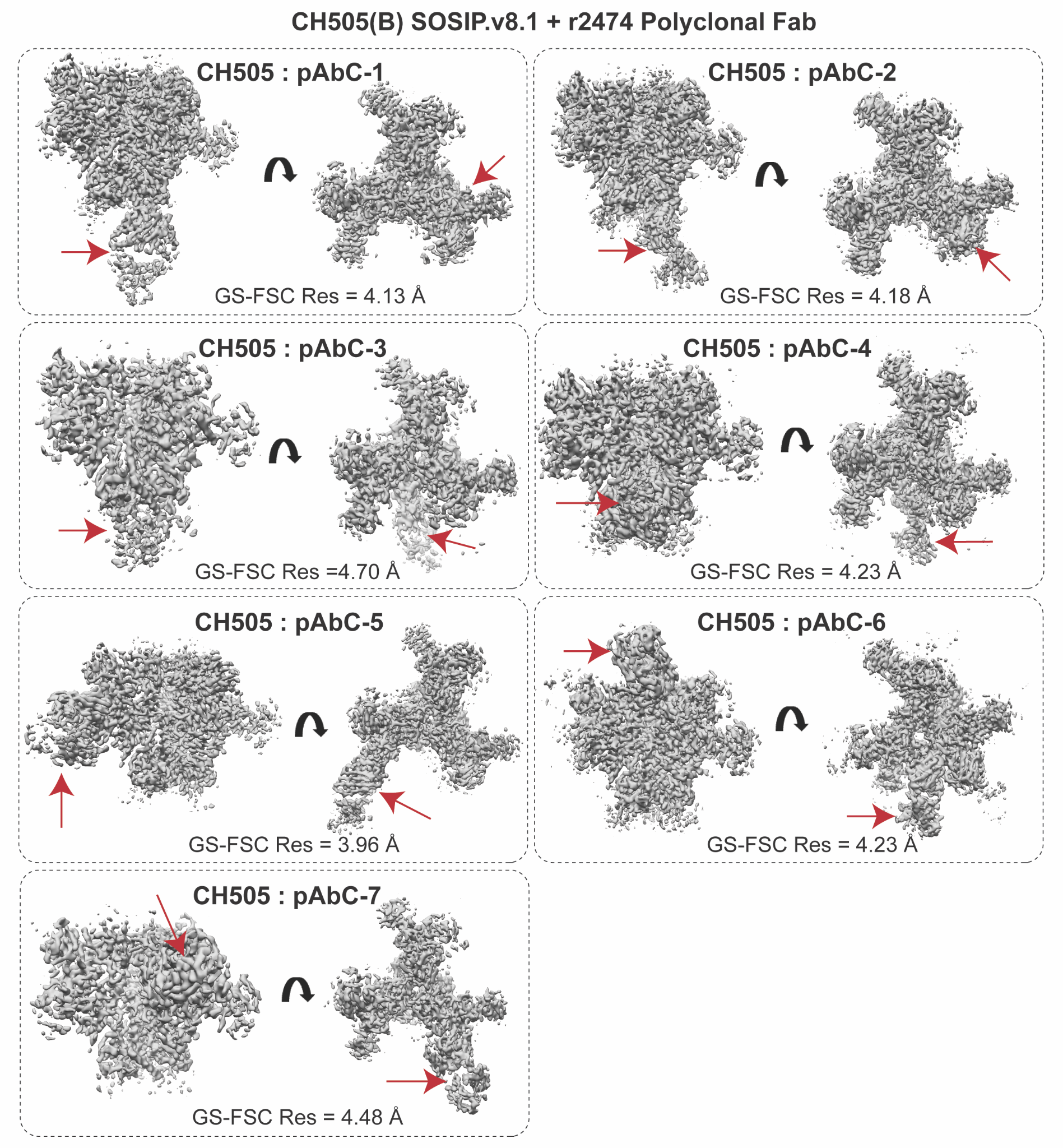

**Fig S11.** EM density maps recovered by cryoEMPEM of CH505(B) SOSIP.v8.1 in complex with r2474 polyclonal antibodies (as Fab)**.** Red arrows point towards the Fab-corresponding density.

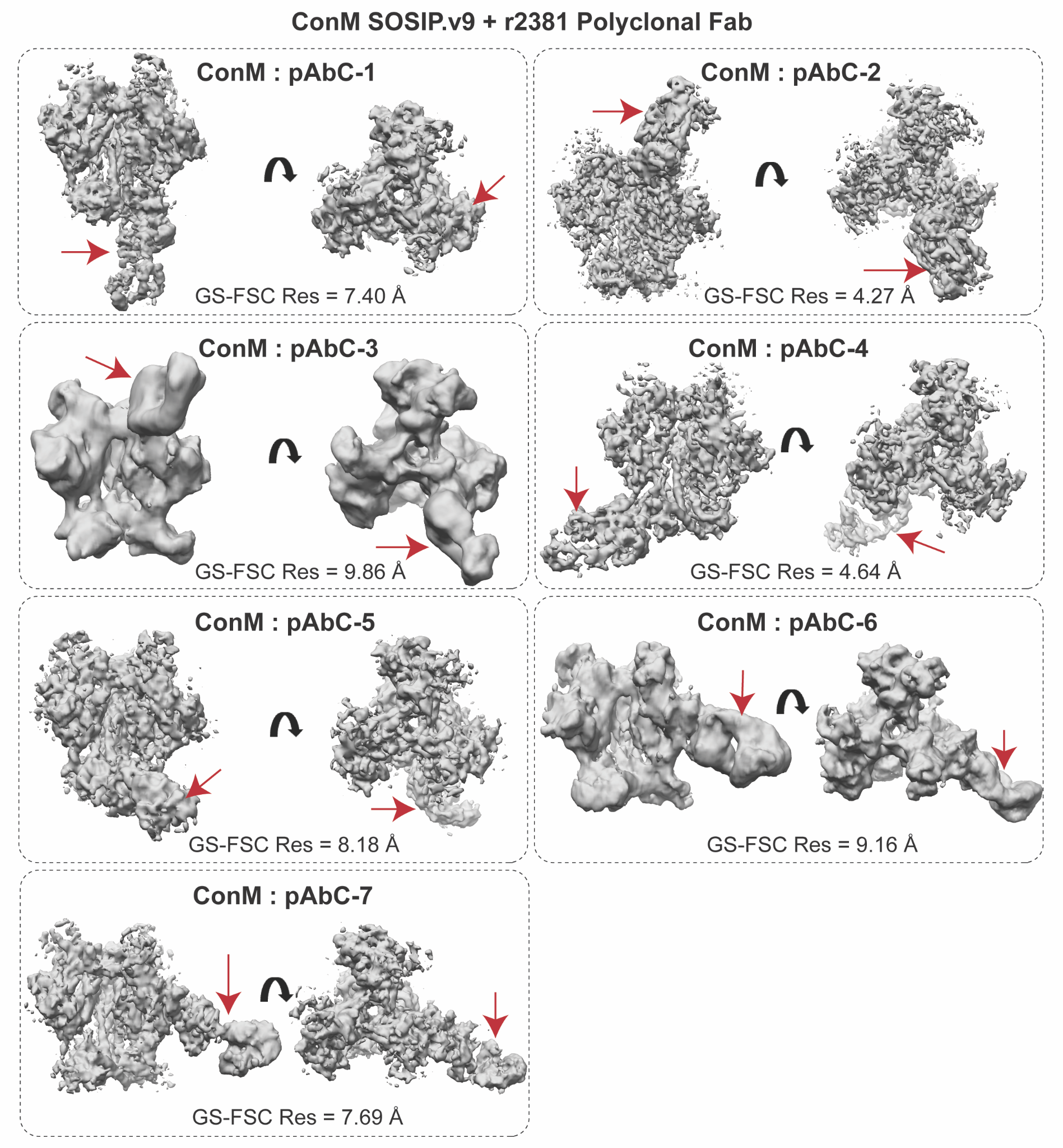

**Fig S12.** EM density maps recovered by cryoEMPEM of ConM SOSIP.v9 in complex with r2381 polyclonal antibodies (as Fab)**.** Red arrows point towards the Fab-corresponding density.

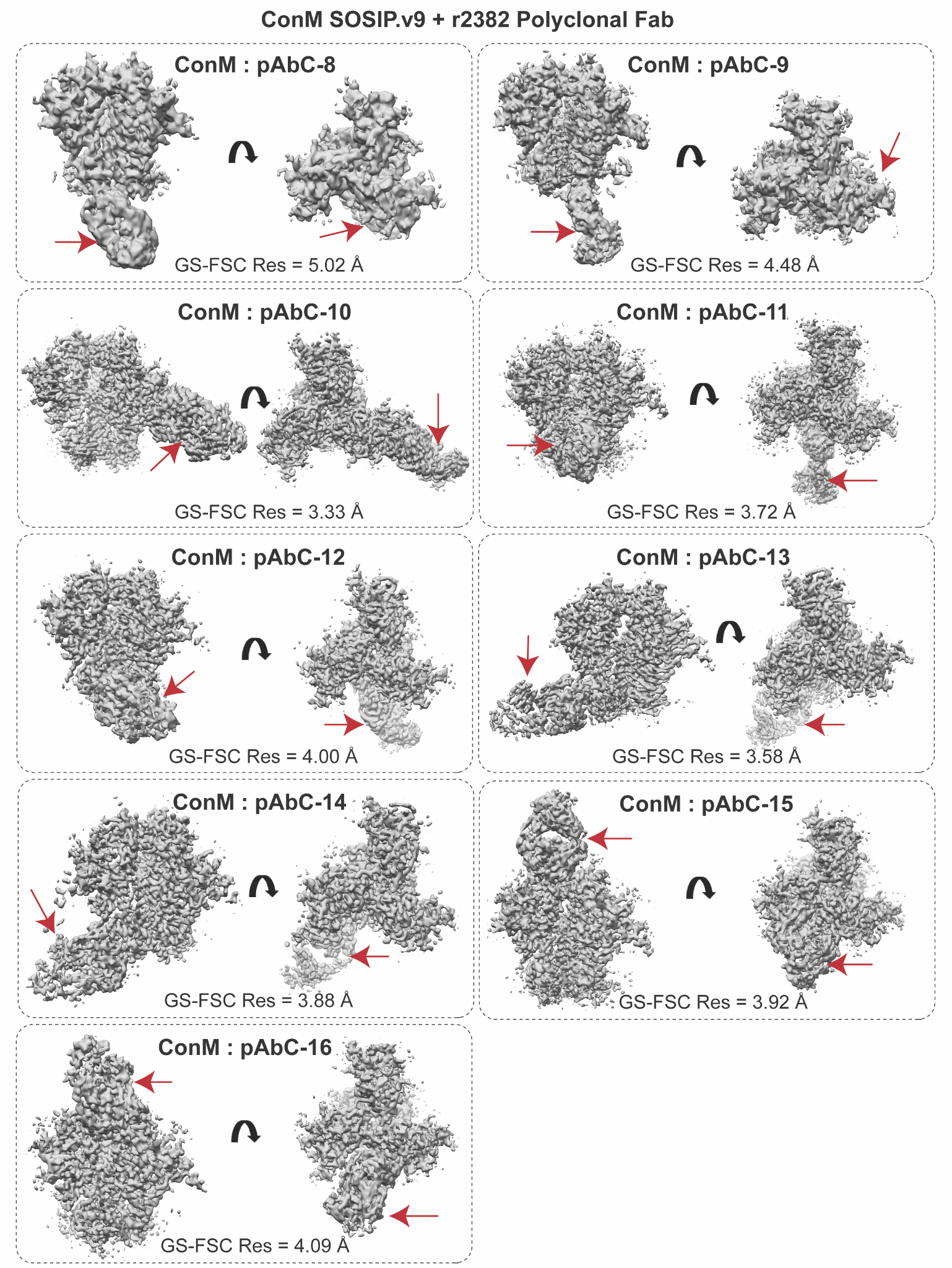

**Fig S13.** EM density maps recovered by cryoEMPEM of ConM SOSIP.v9 in complex with r2382 polyclonal antibodies (as Fab)**.** Red arrows point towards the Fab-corresponding density.

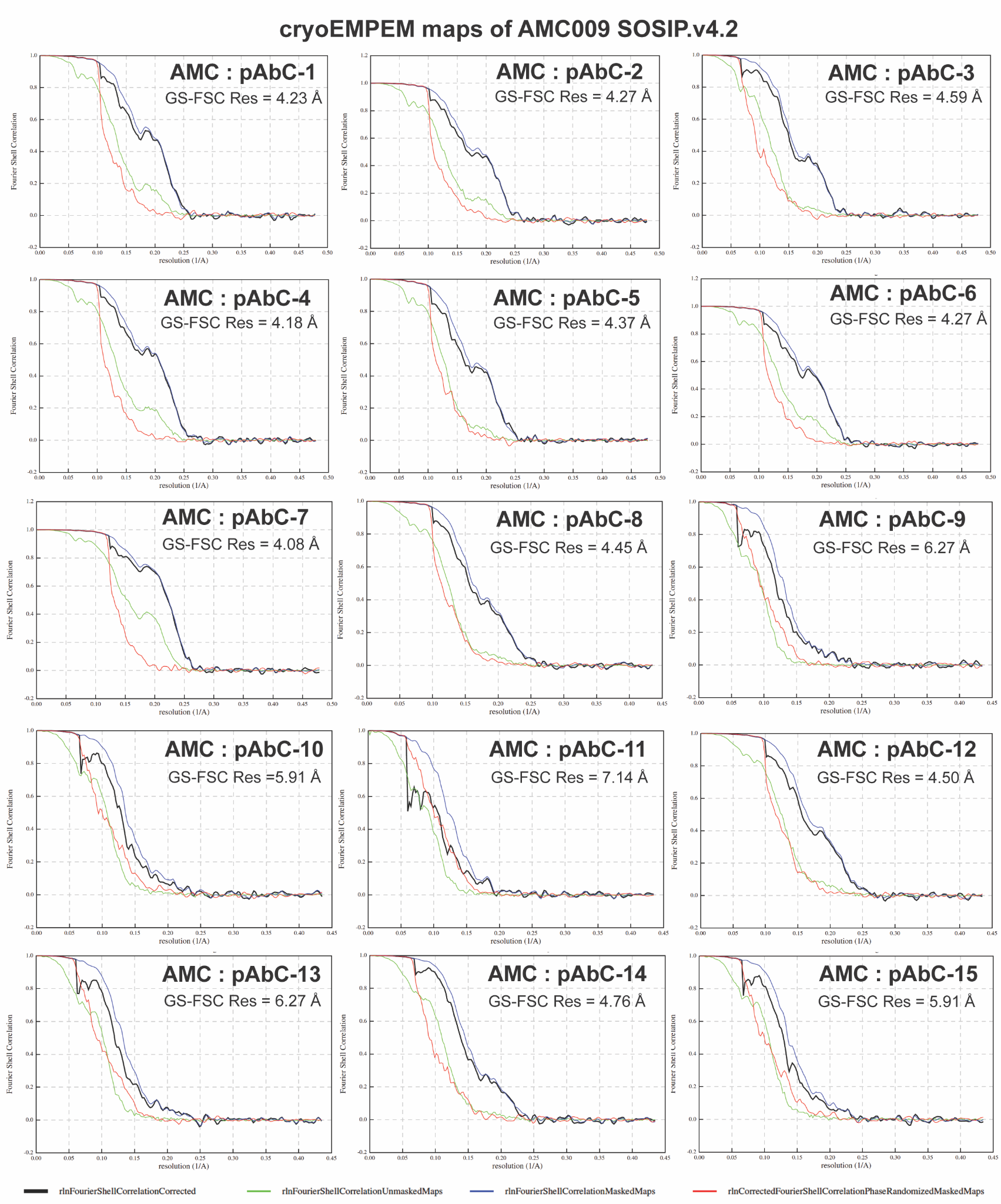

**Fig S14.** Fourier Shell Correlation curves of cryoEMPEM maps recovered for AMC009 SOSIP.v4.2 in complex with polyclonal antibodies from UA0062 and UA0065.

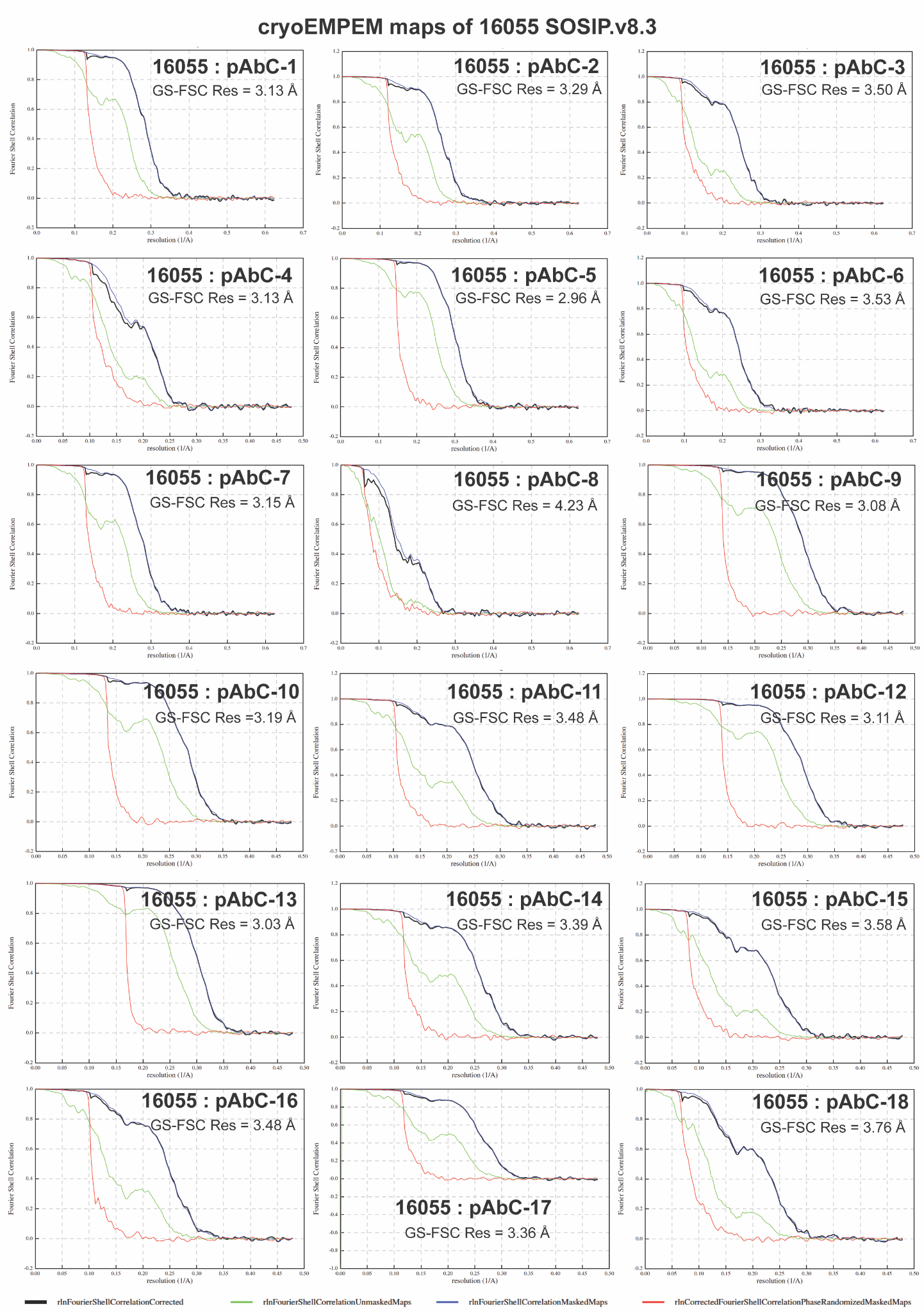

**Fig S15.** Fourier Shell Correlation curves of cryoEMPEM maps recovered for 16055 SOSIP.v8.3 in complex with polyclonal antibodies from r2463 and r2464.

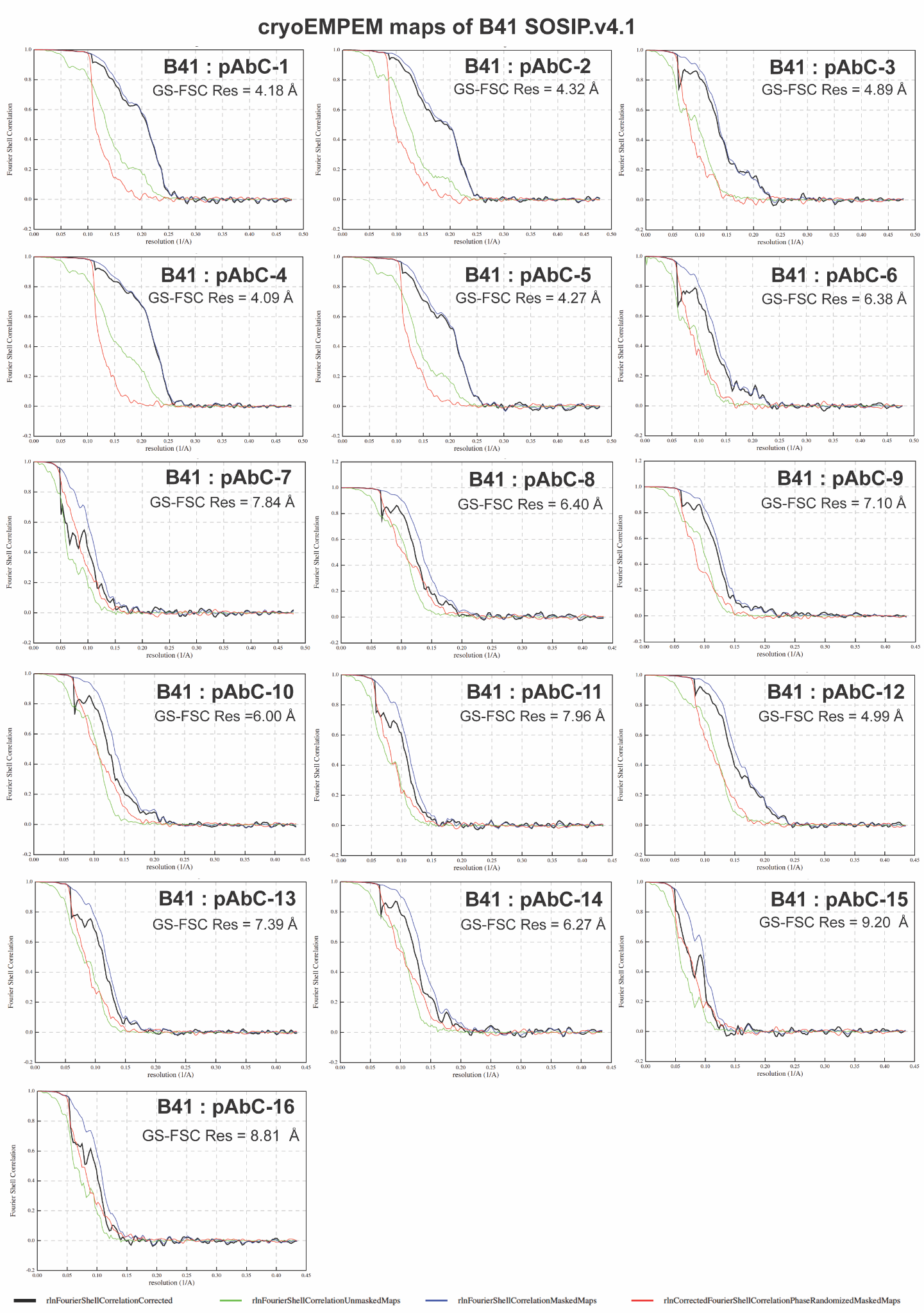

**Fig S16.** Fourier Shell Correlation curves of cryoEMPEM maps recovered for B41 SOSIP.v4.1 in complex with polyclonal antibodies from r1645 and r1646.

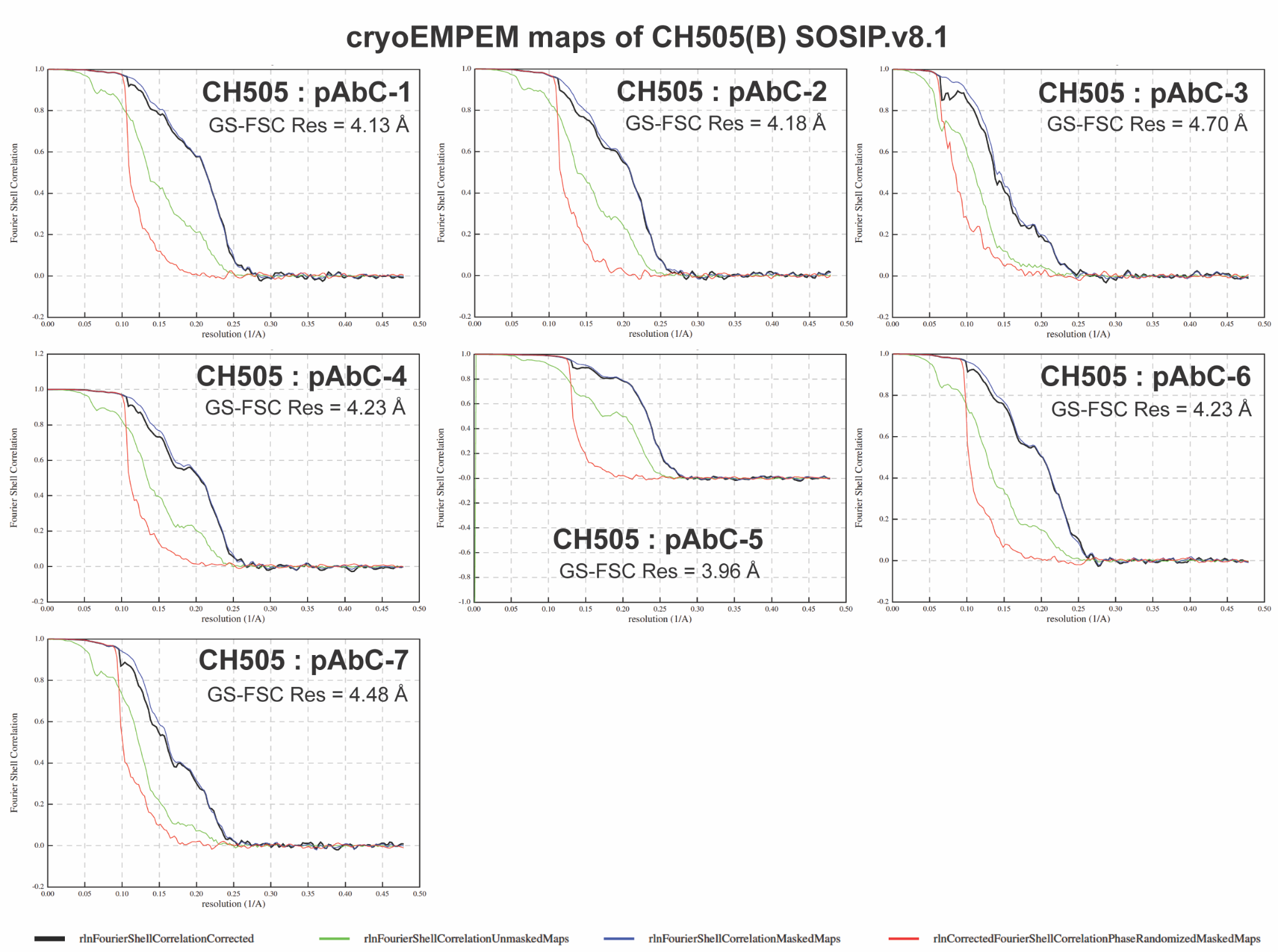

**Fig S17.** Fourier Shell Correlation curves of cryoEMPEM maps recovered for CH505(B) SOSIP.v8.1 in complex with polyclonal antibodies from r2474.

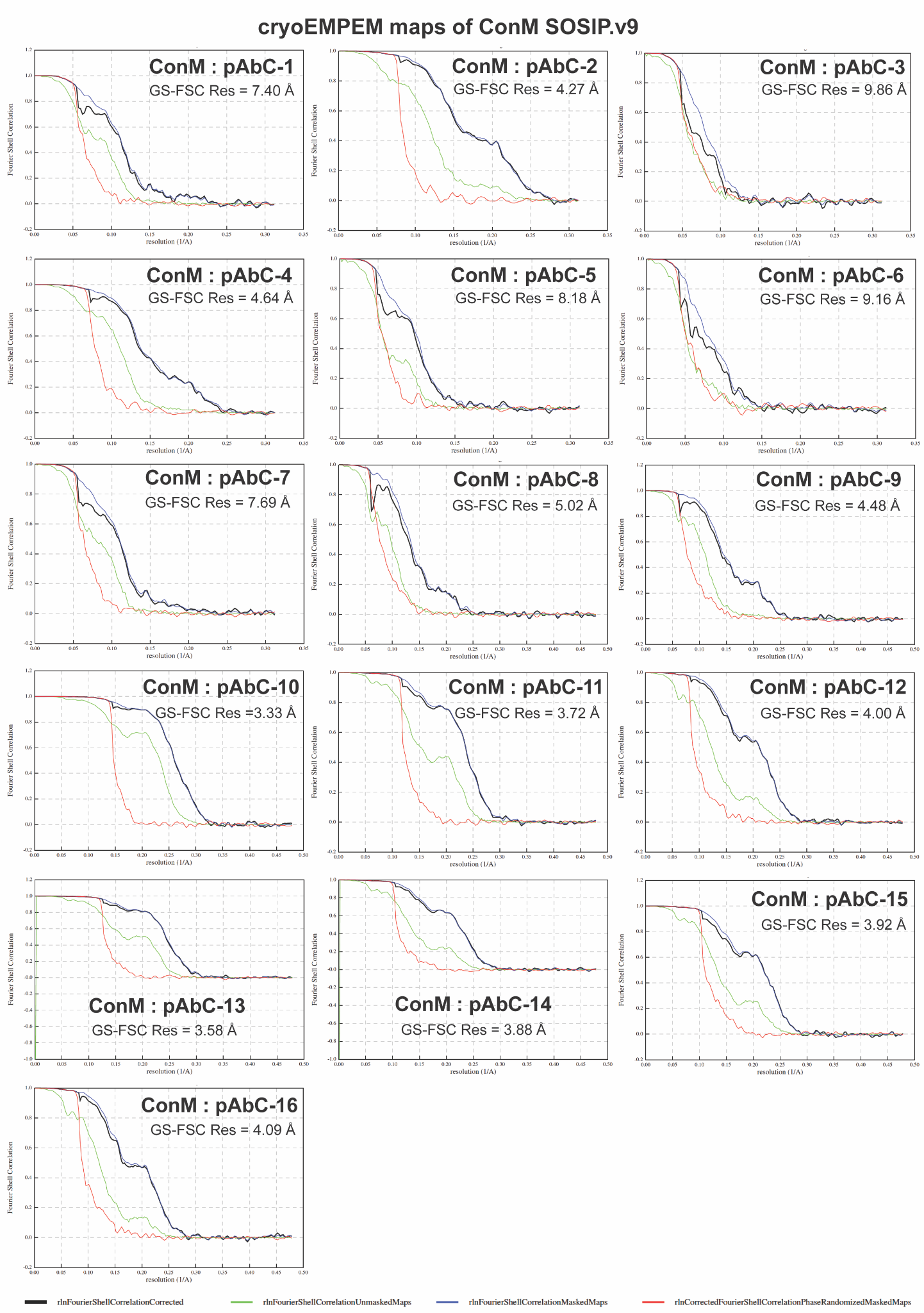

**Fig S18.** Fourier Shell Correlation curves for cryoEMPEM maps recovered for ConM SOSIP.v9 in complex with polyclonal antibodies from r2381 and r2382.

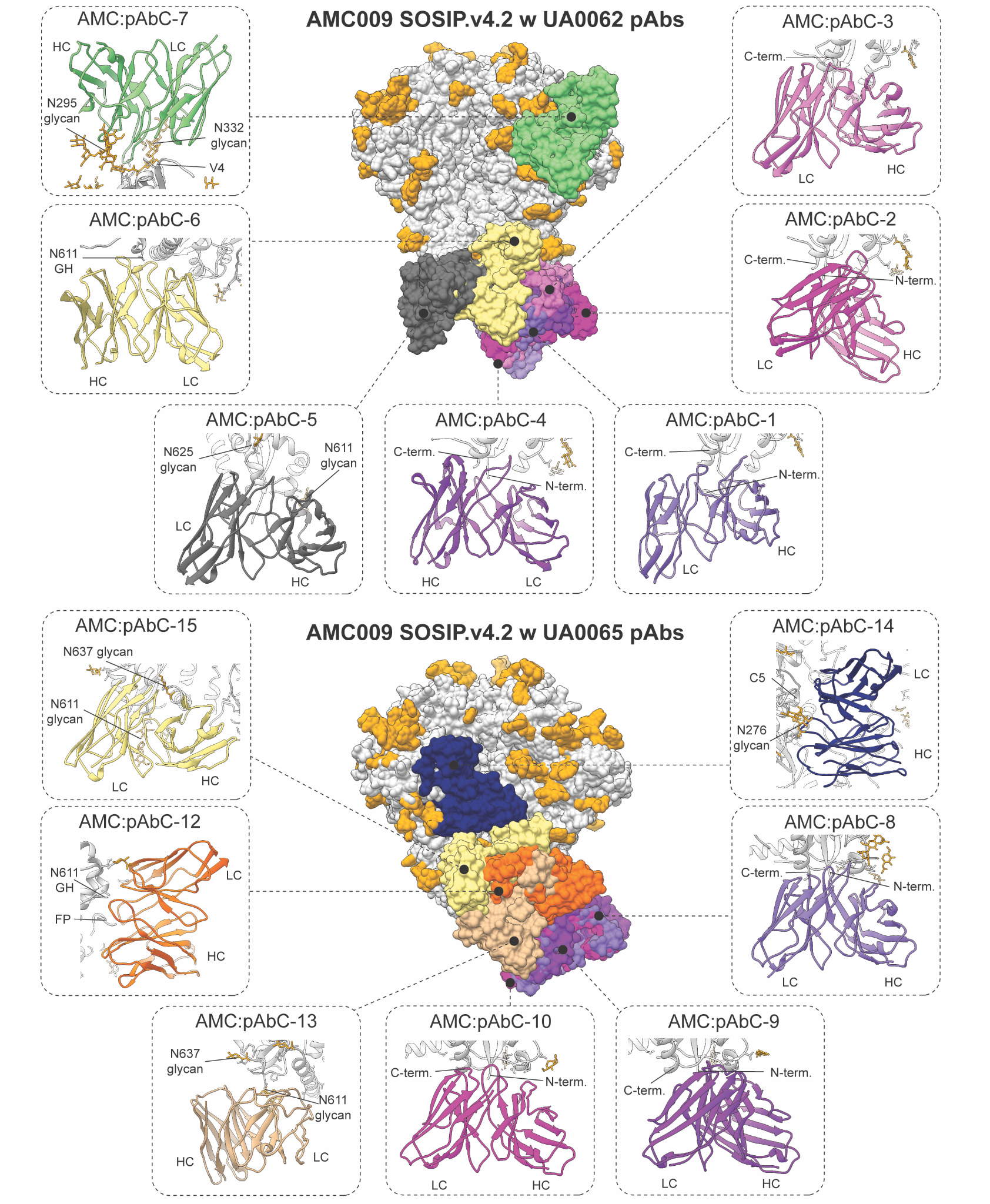

**Fig S19.** pAbC models recovered by cryoEMPEM of AMC009 SOSIP.v4.2 in complex with UA0062 and UA0065 polyclonal antibodies (as Fab)**.** Antigen is presented in gray with glycans in yellow. Antibodies are colored according to the epitope (see Fig. S2 for epitope color chart).

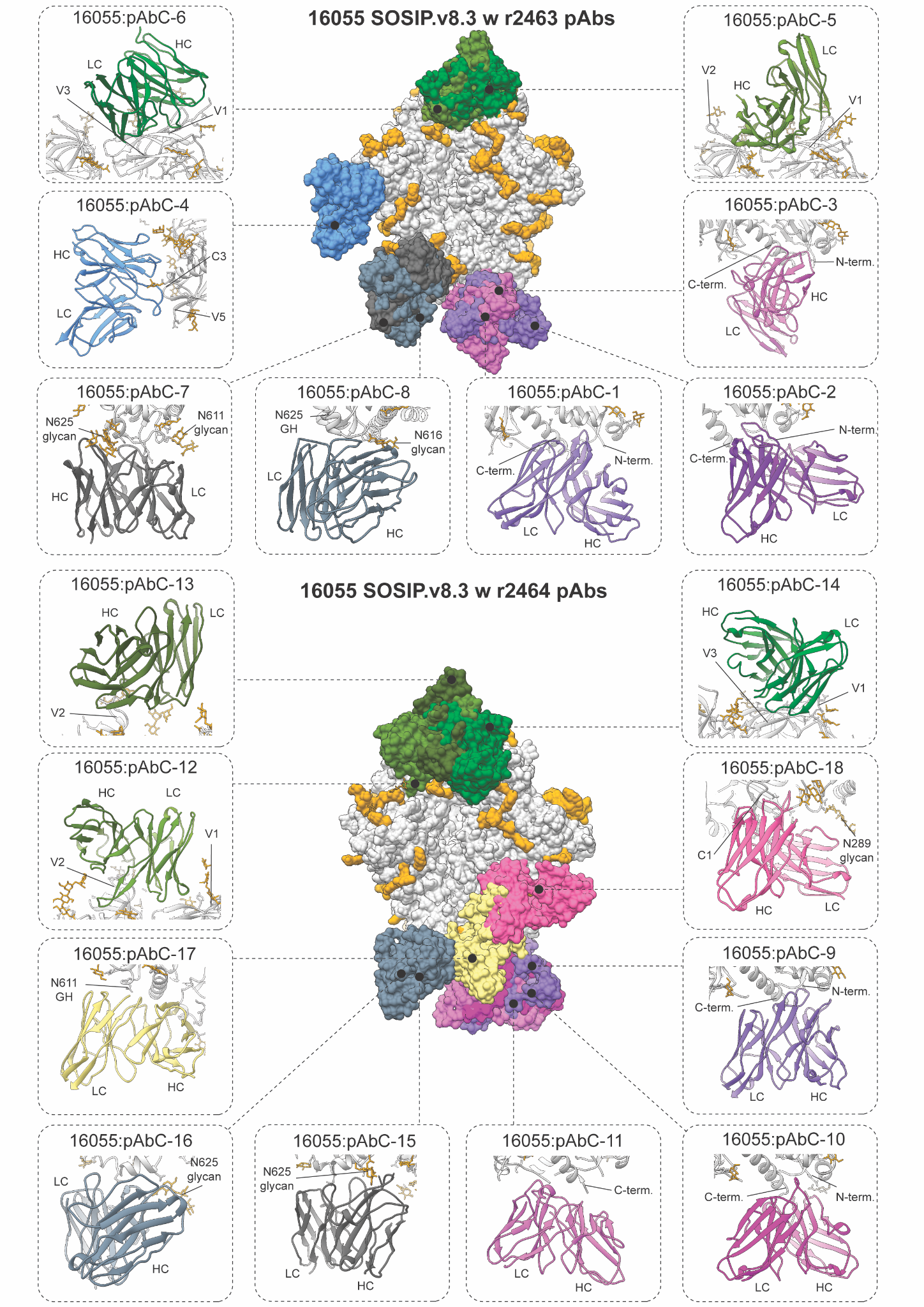

**Fig S20.** pAbC models recovered by cryoEMPEM of 16055 SOSIP.v8.3 in complex with r2463 and r2464 polyclonal antibodies (as Fab)**.** Antigen is presented in gray with glycans in yellow. Antibodies are colored according to the epitope (see Fig. S2 for epitope color chart).

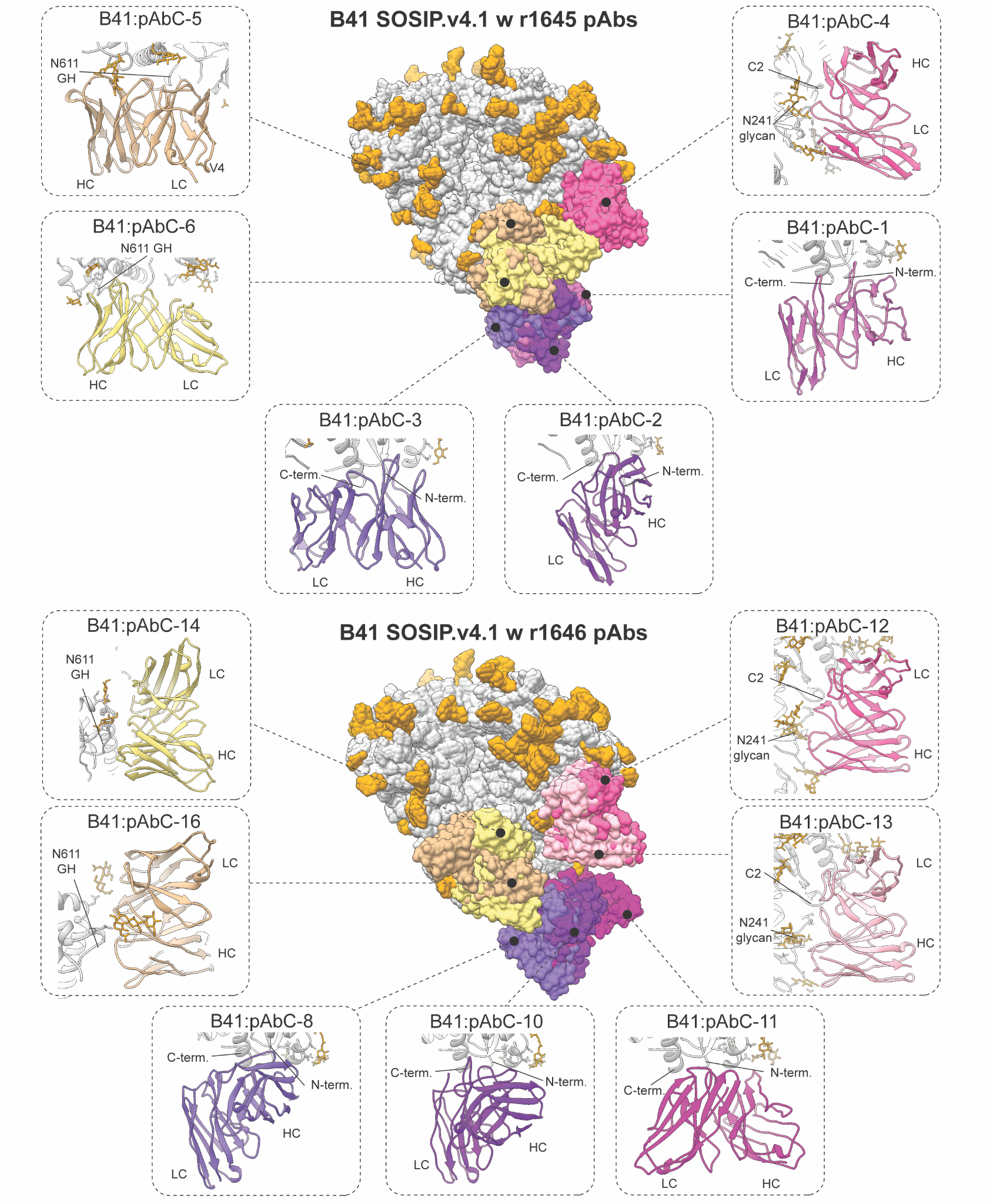

**Fig S21.** pAbC models recovered by cryoEMPEM of B41 SOSIP.v4.1 in complex with r1645 and r1646 polyclonal antibodies (as Fab)**.** Antigen is presented in gray with glycans in yellow. Antibodies are colored according to the epitope (see Fig. S2 for epitope color chart).

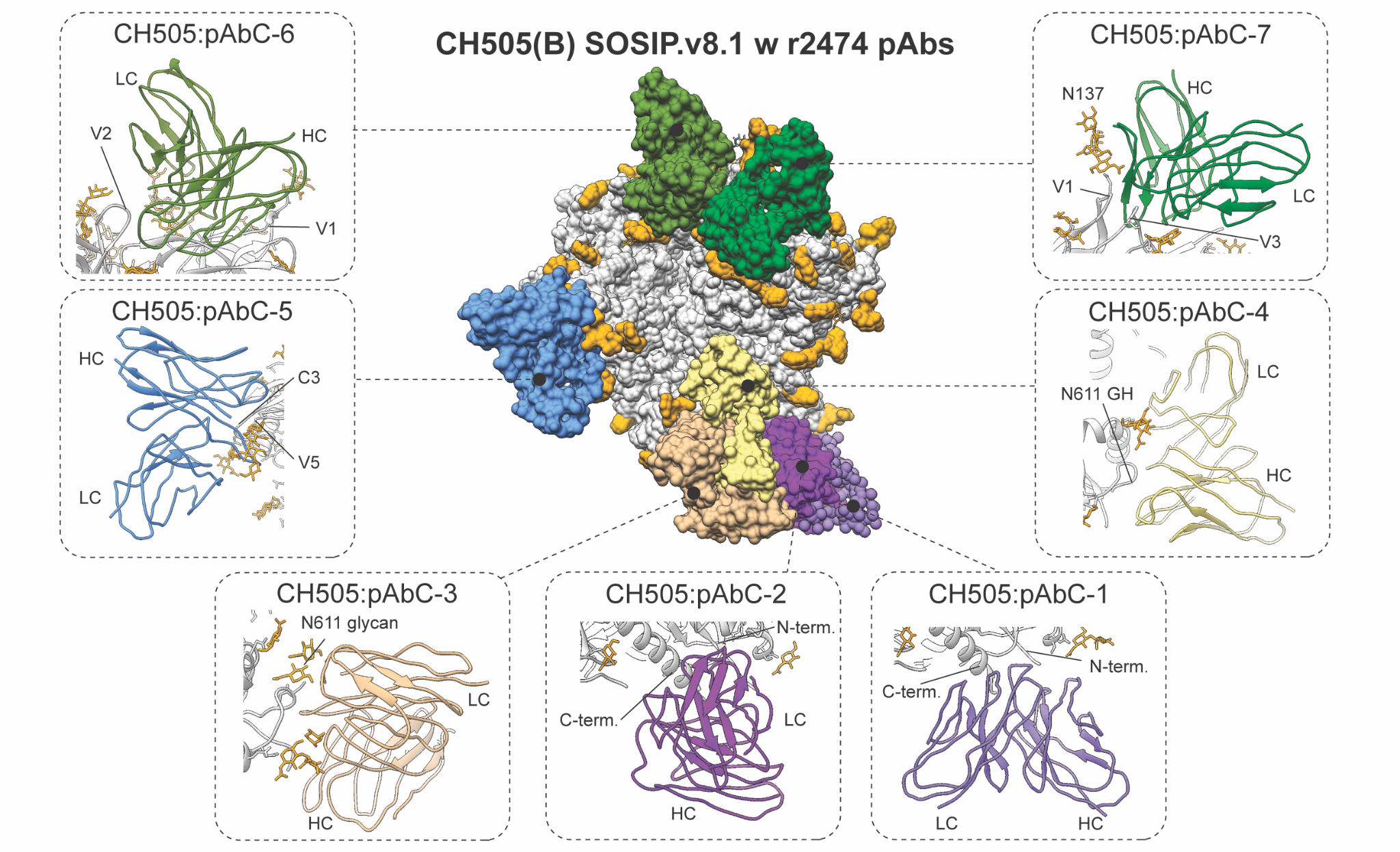

**Fig S22.** pAbC models recovered by cryoEMPEM of CH505(B) SOSIP.v8.1 in complex with r2474 polyclonal antibodies (as Fab)**.** Antigen is presented in gray with glycans in yellow. Antibodies are colored according to the epitope (see Fig. S2 for epitope color chart).

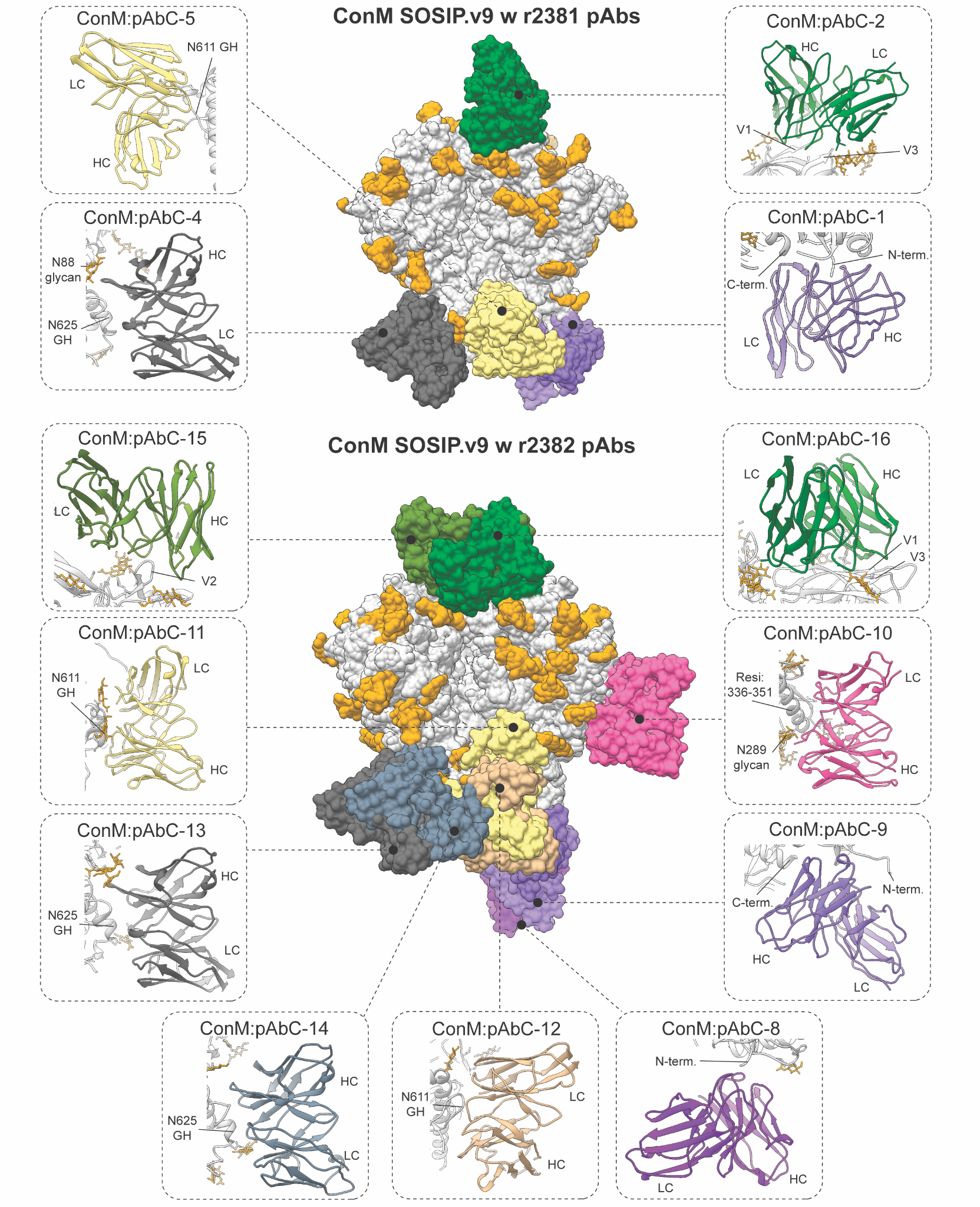

**Fig S23.** pAbC models recovered by cryoEMPEM of ConM SOSIP.v9 in complex with r2381 and r2382 polyclonal antibodies (as Fab)**.** Antigen is presented in gray with glycans in yellow. Antibodies are colored according to the epitope (see Fig. S2 for epitope color chart).

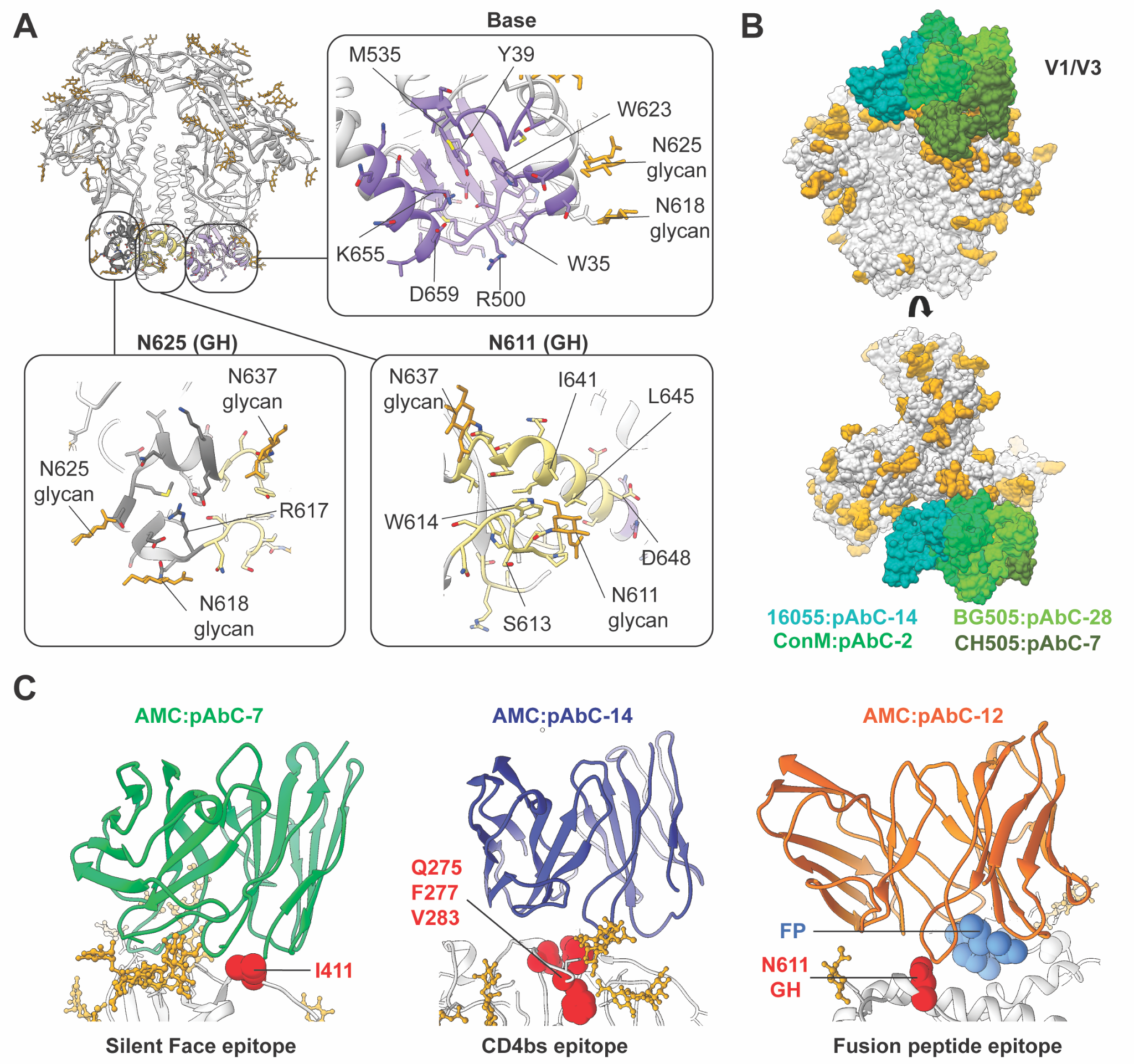

**Fig S24.** (**A**) Base-proximal epitopes on HIV-1 Env with key features (amino acids and glycans) labeled; the BG505 SOSIP sequence was used as reference. (**B**) Overlay of V1/V3-directed pAbCs recovered from different Env antigens. A surface model illustrates differences in angle of approach. The antigen is shown in gray with glycans in yellow, and antibodies are depicted in different shades of green. (**C**) Three pAbCs targeting bnAb-like epitopes on AMC009 SOSIP.v4.2. Antibody Fv domains are colored according to epitope; the antigen is shown as a gray cartoon with glycans in yellow. Key residues present in AMC009 but absent in other Envs are shown as red spheres. For AMC:pAbC-7 (left), I411 replaces the typical N411 PNGS. For AMC:pAbC-14 (middle), Q275, F277, and V283 are unique to AMC009. For AMC:pAbC-12 (right), N611 within a PNGS is unglycosylated, and interaction with the FP (blue) depends on this incomplete glycan occupancy (glycan hole).

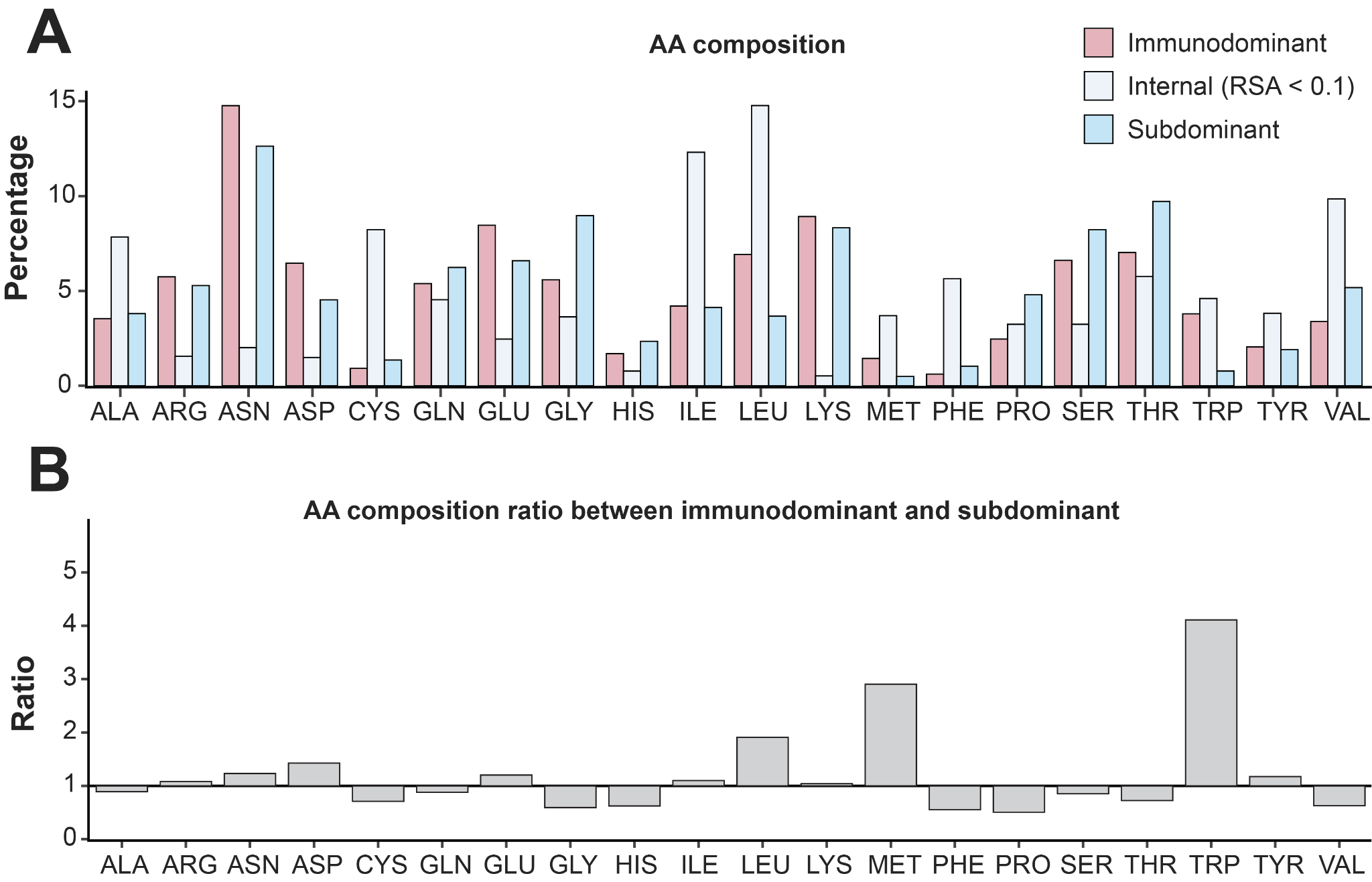

**Fig S25. (A)** Bar plots show the percentage contribution of each amino acid across immunodominant and subdominant surfaces, as well as all internal residues (defined based on having RSA < 0.1) across all Env-antibody complexes. (**B**) Bar graph showing the relative enrichment (ratio) of individual surface-exposed amino acids (RSA > 0.1) in immunodominant epitopes compared to subdominant.

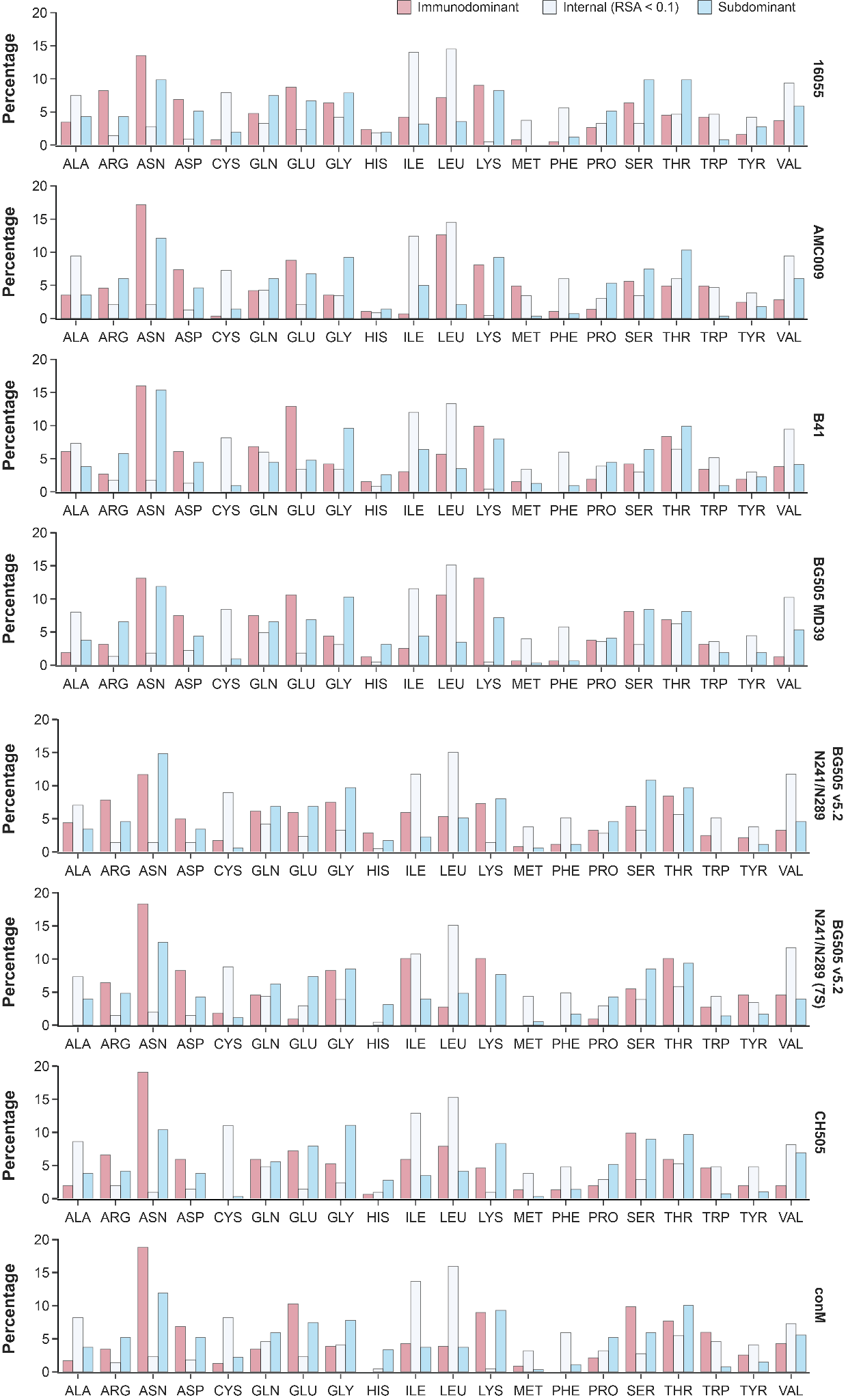

**Fig S26.** Bar plots showing the relative percentage of each amino acid across immunodominant and subdominant surface sites as well as all among internal residues (defined as RSA < 0.1). The results are shown for each antigen separately based on structural data recovered by cryoEMPEM.

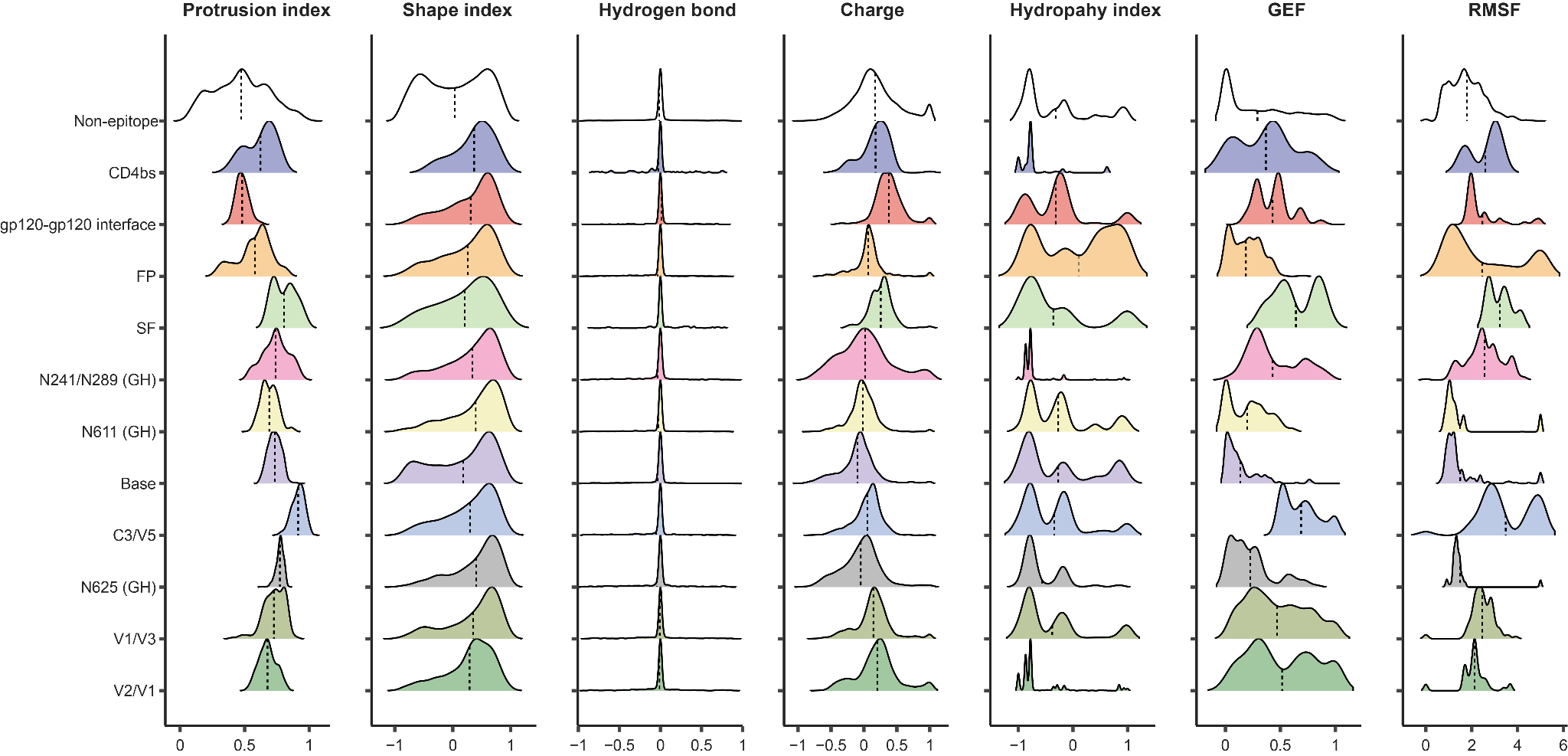

**Fig S27.** Violin plots showing the distribution of feature values split by epitope class.

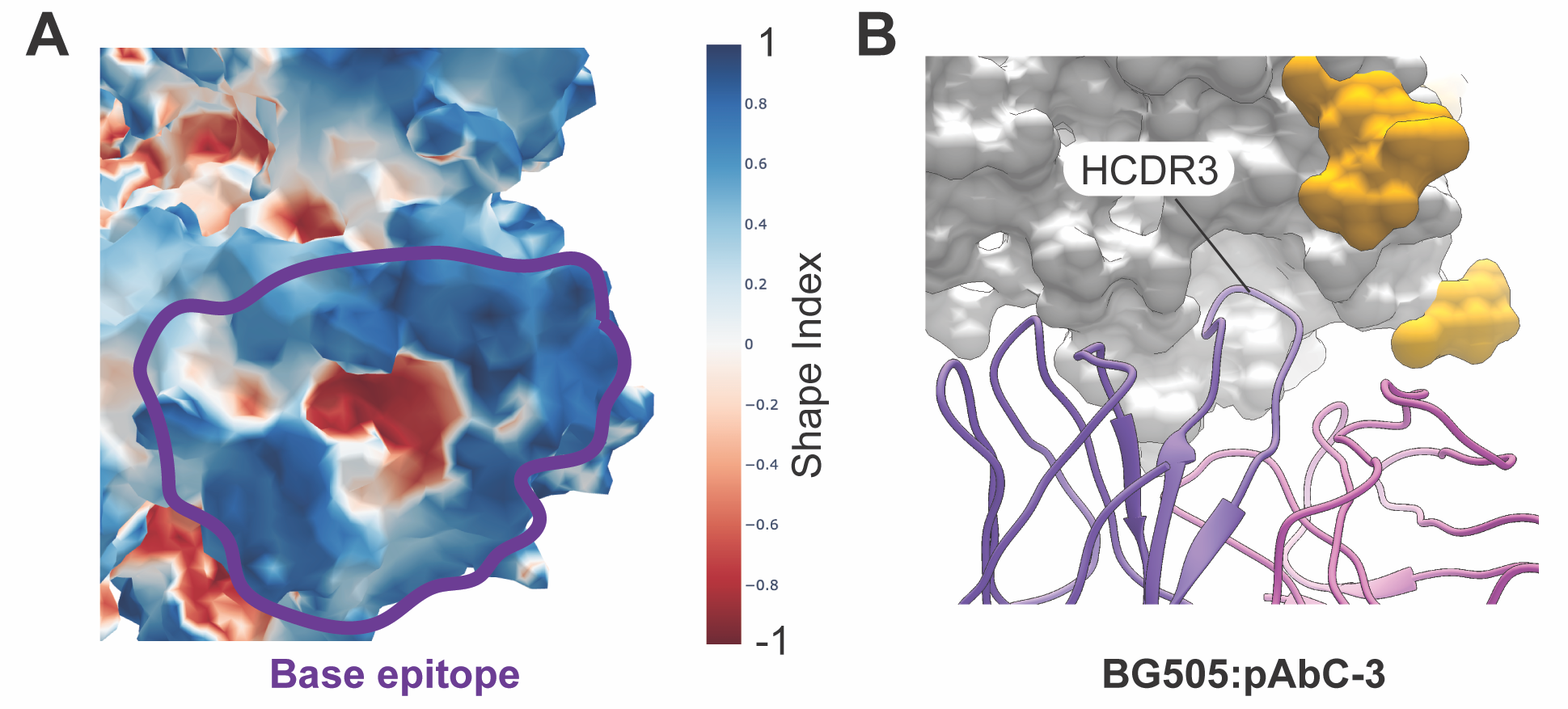

**Fig S28.** (**A**) Shape index mapped onto the surface of the base epitope, with the epitope boundary highlighted in purple. The central pocket is highly concave, whereas most of the surrounding epitope surface is convex. (**B**) Common binding mode of base-targeting antibodies, which typically involves insertion of a CDR loop (most often HCDR3) into the central pocket.

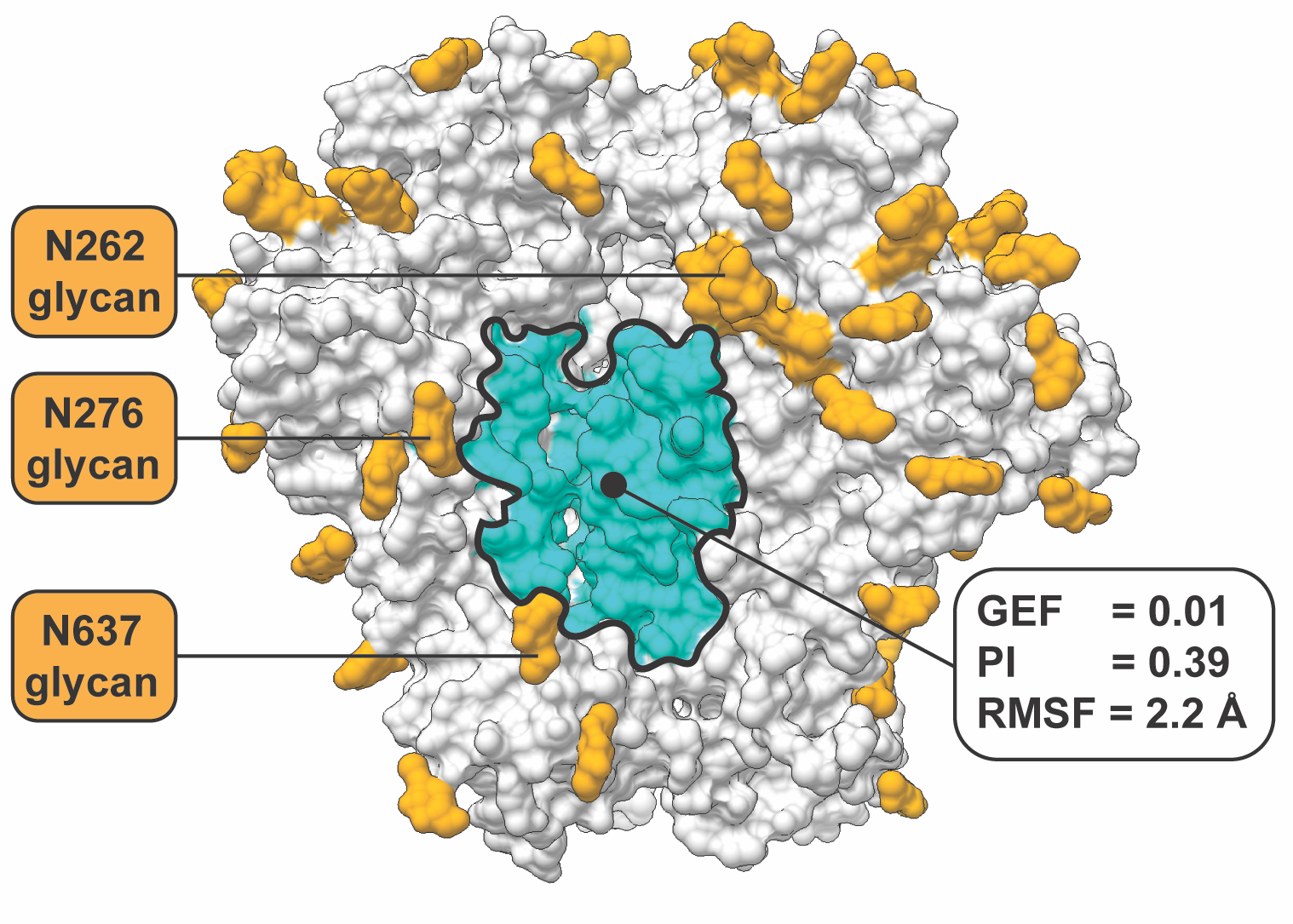

**Fig S29.** Subdominant region in the center of Env comprising segments at the interface of C1 and HR1. The flanking glycans are shown on the left while the most relevant surface metrics are shown on the right.

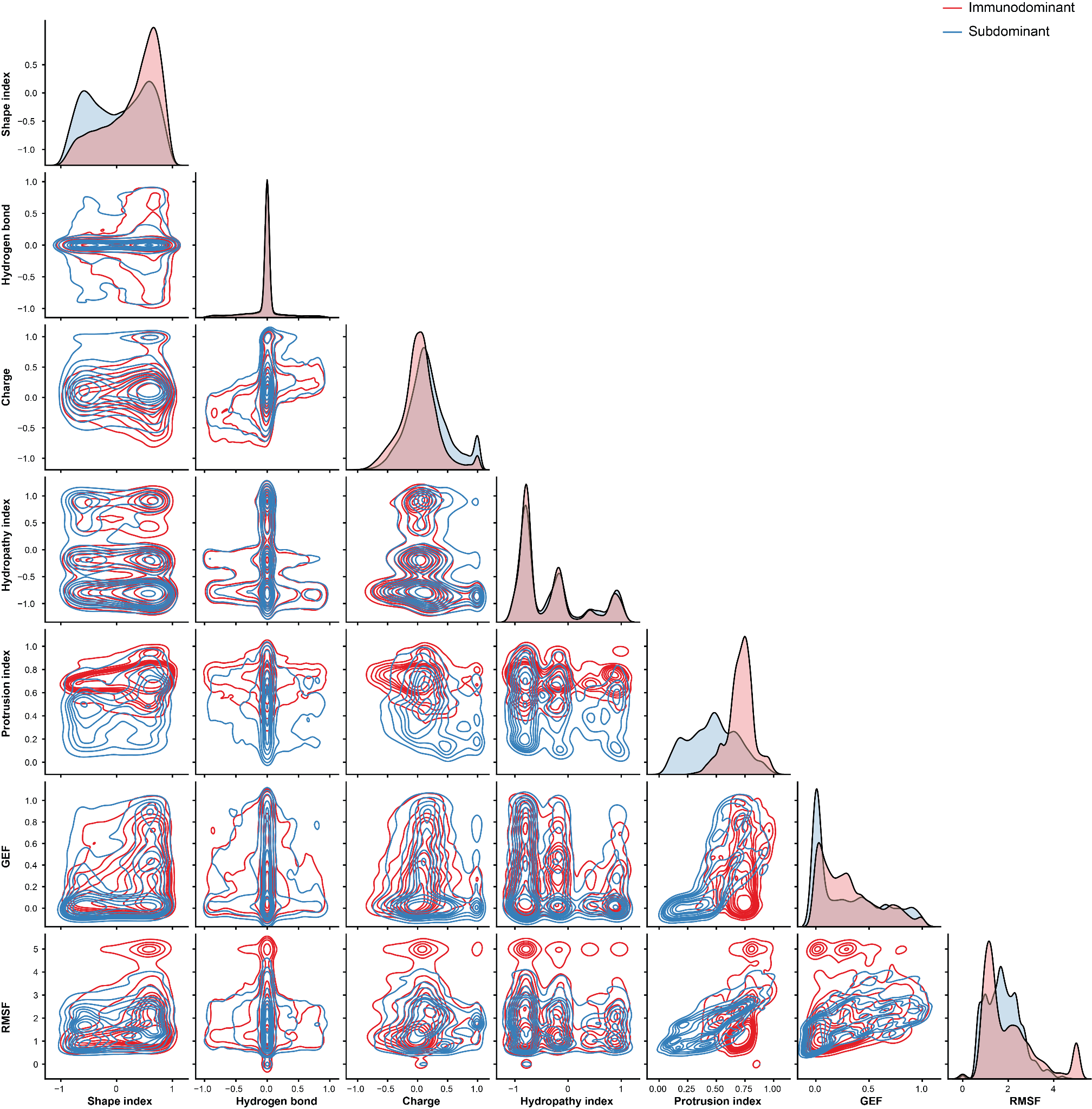

**Fig S30.** Matrix of bivariate scatter plots showing the correlation among features between immunodominant (red) and subdominant sites (blue).

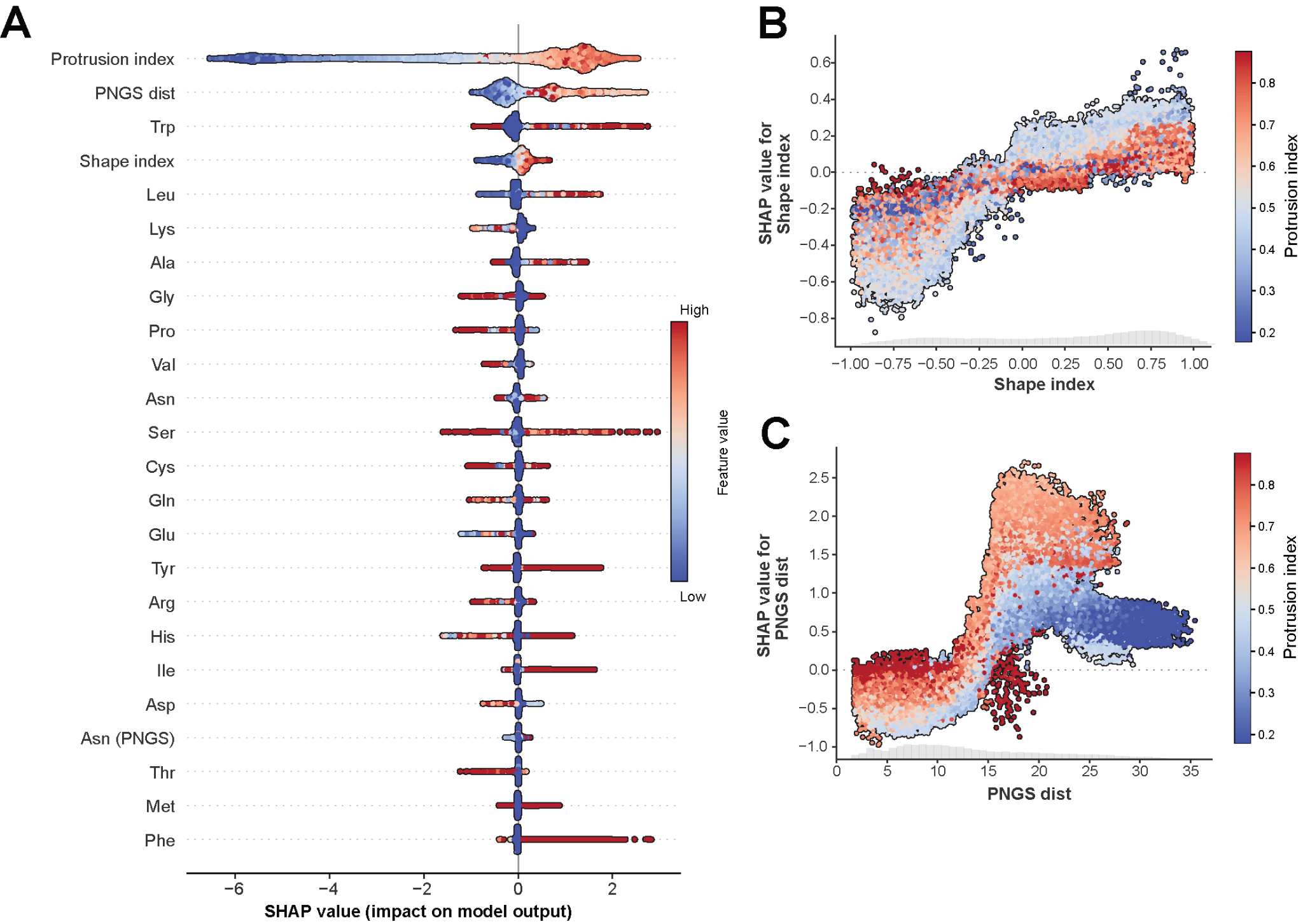

**Fig S31. (A)** Summary plot of SHAP value distributions for all features ranked by their importance. **(B)** SHAP dependence plots for SI, with each data point’s feature value on the x-axis and the corresponding SHAP value on the y-axis, colored by its PI value. **(C)** SHAP dependence plots for PNGS dist, with each data point’s feature value on the x-axis and the corresponding SHAP value on the y-axis, colored by its PI value.

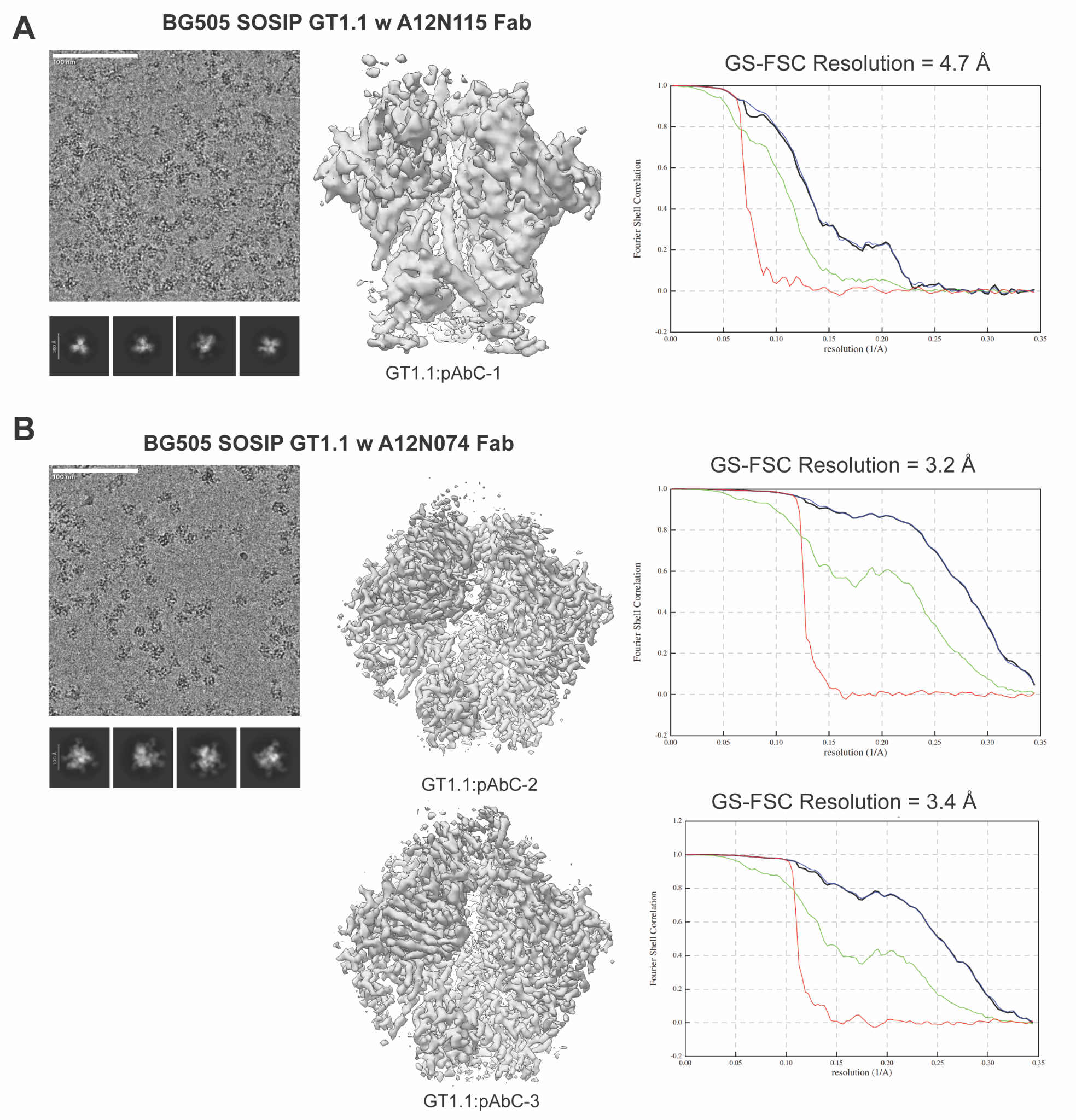

**Fig S32.** Representative raw micrograph, 2D class averages, reconstructed 3D maps and the corresponding Fourier Shell Correlation plots for BG505 SOSIP GT1.1 in complex with polyclonal antibodies from animal (**A**) A12N115 or (**B**) A12N074. Partial signal subtraction was employed on the A12N074 dataset due to high occupancy of non-CD4bs Fabs. Non-subtracted maps are available upon request.

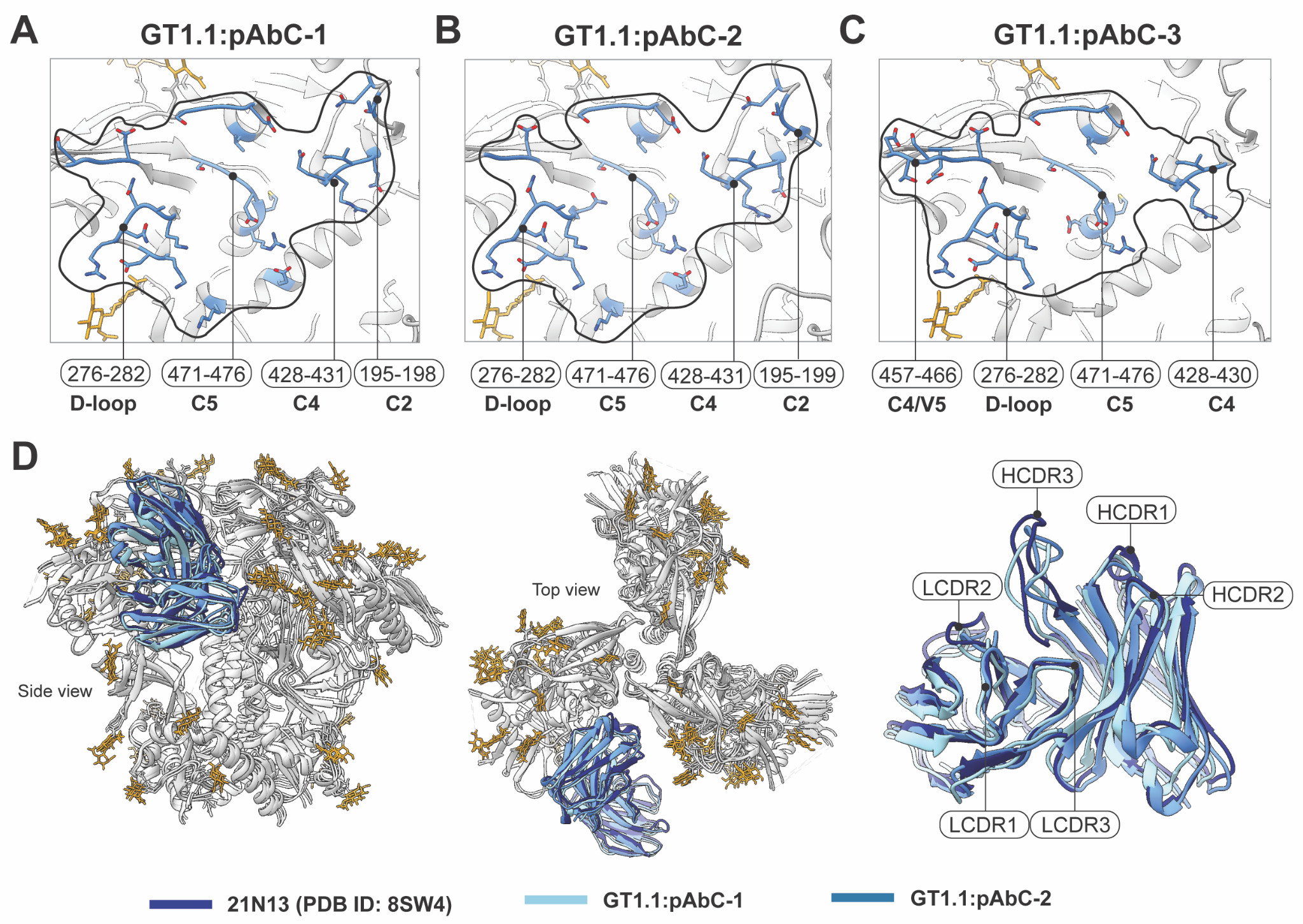

**Fig S33.** Binding footprints of cryoEMPEM-recovered polyclonal antibodies targeting the CD4bs, (**A**) GT1.1:pAbC-1, (**B**) GT1.1:pAbC-2, and (**C**) GT1.1:pAbC-3. The contact residues in each epitope are shown in blue, while the neighboring glycans are in yellow. The overall binding footprint is marked with black line. Note the high similarity in footprints of pAbC-1 and pAbC-2 and their combined difference when compared to pAbC-3 that relies more heavily on contacts in the C4/V5 region. (**D**) Overlay of the 21N13 mAb structure with GT1.1:pAbC-1 and GT1.1:pAbC-2 antibodies recovered by cryoEMPEM. On the left is the structural comparison of the entire complex including the antigen. On the right is the enlarged view of the Fv region of each antibody with different CDR loops illustrated.

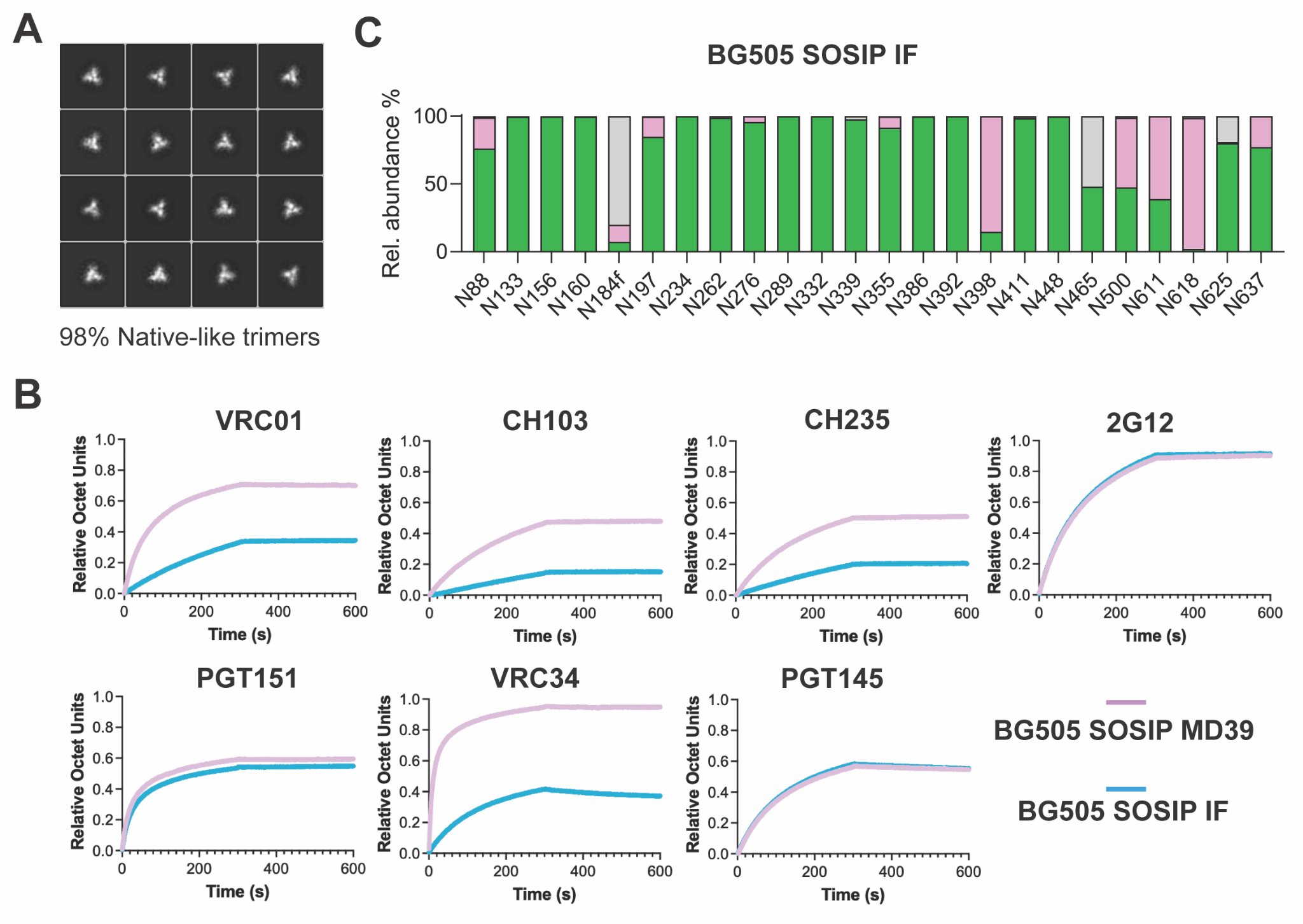

**Fig S34. (A)** 2D class averages of the nsEM experiment performed on BG505 SOSIP IF. The percentage is based on quantification of the number of particles in trimer-like and non-trimer classes. **(B)** BLI-based antigenicity evaluation of the BG505 SOSIP IF compared to BG505 SOSIP MD39, which lacks the immunofocusing (IF) mutations. Antibody name is indicated above the corresponding panel.  **(C)** The composition of N-linked glycan PNGS is displayed, with PNGS aligned to HxB2. Sites for which glycopeptide data of sufficient quality could not be obtained are not displayed. The abundance of oligomannose- and hybrid-type glycans is colored green, complex-type glycans are colored pink, and the proportion of unoccupied PNGS are displayed in gray.

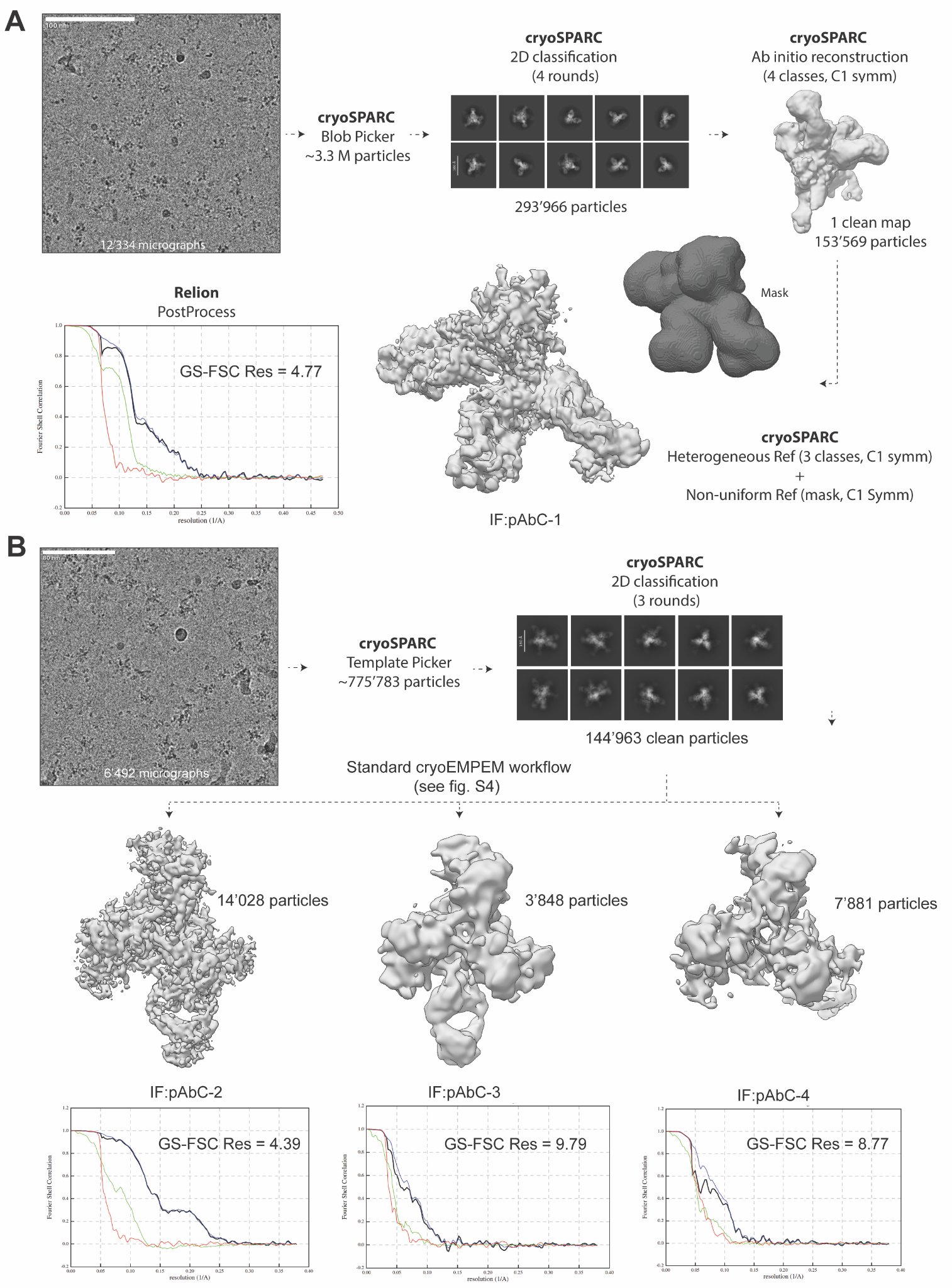

**Fig S35.** Workflow for cryoEMPEM processing of BG505 SOSIP IF in complex with NZWR-1/2 polyclonal antibodies. (**A**) In the first experiment the assembly extended to 18 h. (**B**) In the second experiment the immune complexes assembled for 2 h prior to purification and grid preparation. Representative data and key statistics are shown for each condition.

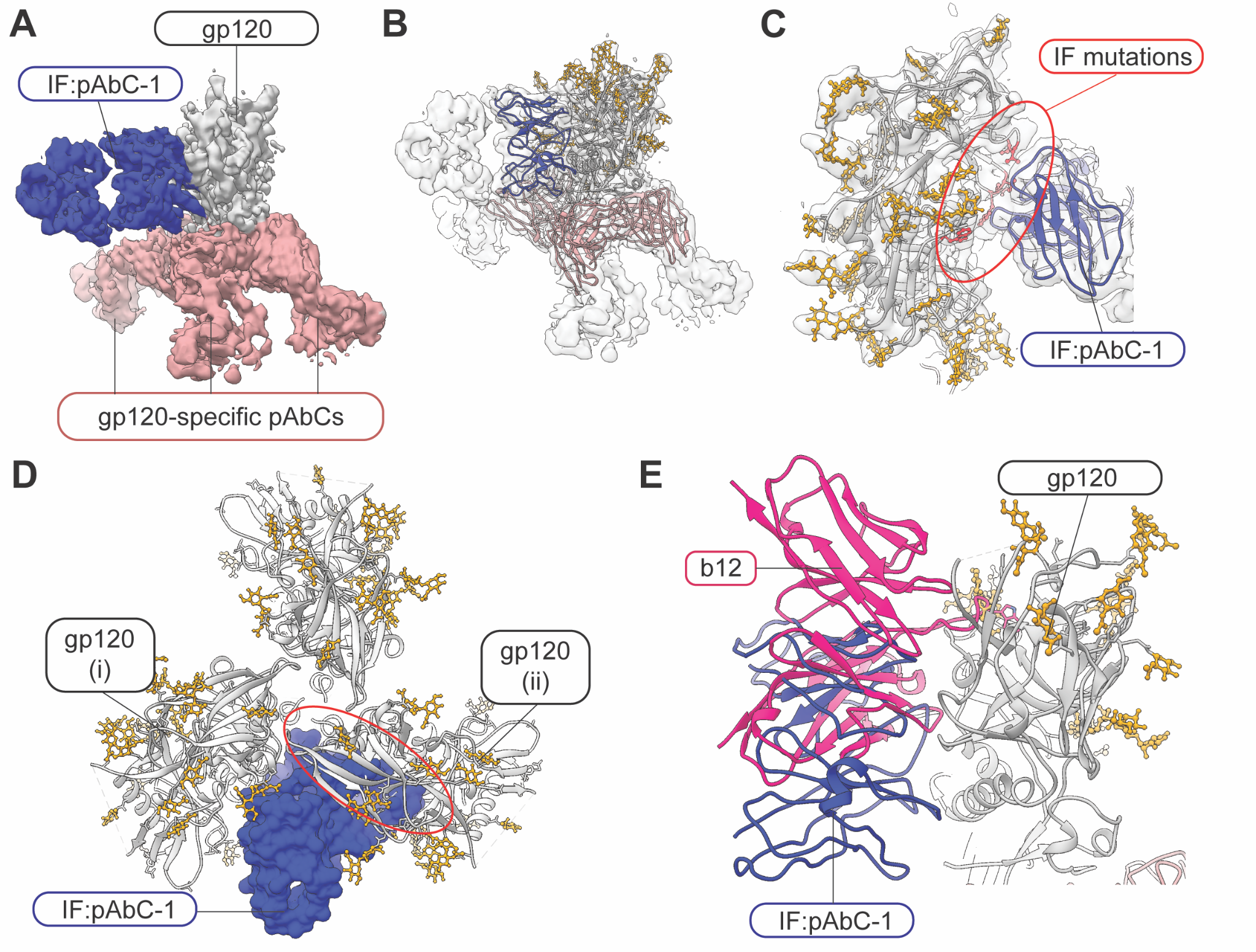

**Fig S36.** (**A**) Segmented map of the IF:pAbC-1 complex, with BG505 SOSIP IF in gray, the CD4bs antibody in purple, and three additional gp120-directed antibodies in salmon. (**B**) Backbone model of the immune complex, colored as in (**A**). (**C**) Close-up of the CD4bs interface, showing bridging density (transparent surface) to the IF mutation sites (red). (**D**) The IF:pAbC-1 antibody (blue surface) is sterically incompatible with trimeric Env (gray cartoon), indicating that binding requires or induces trimer dissociation. (**E**) Overlay of the b12 bnAb (magenta) and IF:pAbC-1 (blue) structures showing partial overlap of their epitopes on Env (gray).

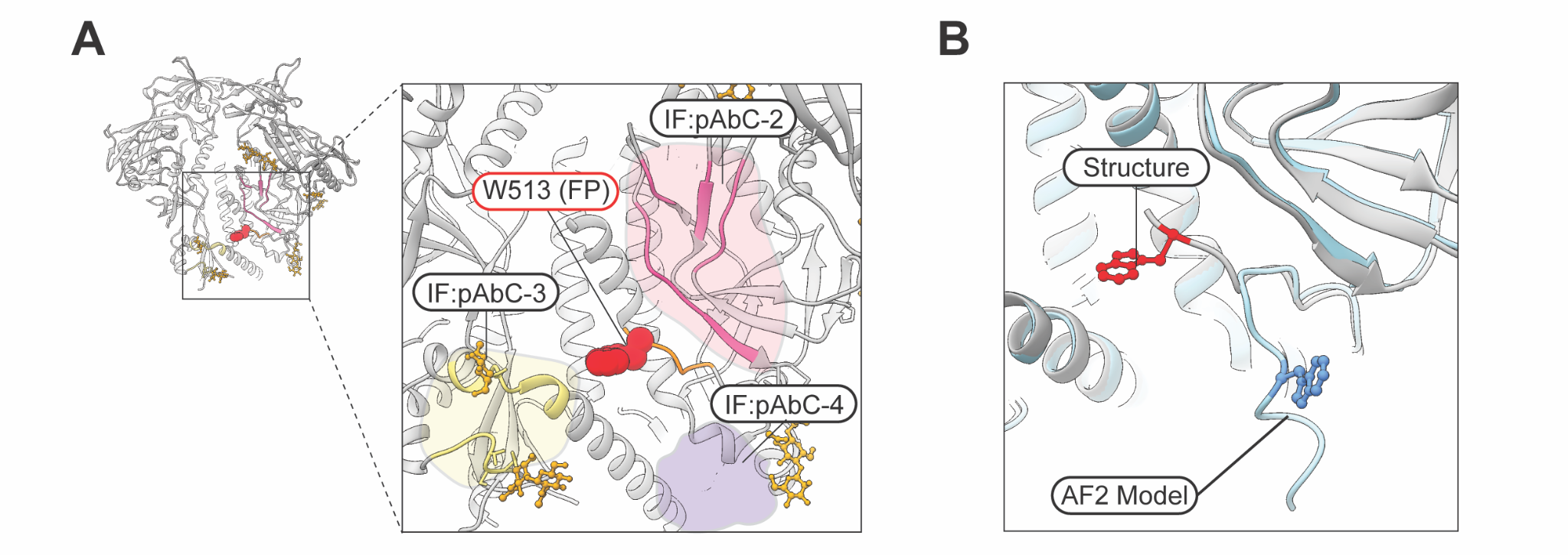

**Fig S37.** (**A**) Epitope footprints of IF:pAbC-2 (pink), IF:pAbC-3 (yellow) and IF:pAbC-4 (purple) mapped onto the surface of BG505 SOSIP (gray). The FP is shown in orange, with W513 highlighted as red spheres. (**B**) Close-up of the FP region in the original AF2 model of BG505 SOSIP IF (cyan, W513 in blue) compared with the experimental structure (gray, W513 in red).
